# Fate-resolved gene regulatory signatures of individual B lymphocytes in the early stages of Epstein-Barr Virus infection

**DOI:** 10.1101/2022.02.23.481342

**Authors:** Elliott D. SoRelle, Joanne Dai, Nicolás M. Reinoso-Vizcaino, Ashley P. Barry, Cliburn Chan, Micah A. Luftig

## Abstract

Epstein-Barr Virus (EBV) infection of B lymphocytes elicits diverse host responses via complex, well-adapted transcriptional control dynamics. Consequently, this host-pathogen interaction provides a powerful system to explore fundamental cellular processes that contribute to consensus fate decisions including cell cycle arrest, apoptosis, proliferation, and differentiation. Here we capture these responses and fates with matched single-cell transcriptomics and chromatin accessibility, from which we construct a genome-wide multistate model of early infection dynamics. Notably, our model captures a previously uncharacterized EBV^+^ analog of a multipotent activated precursor state that can yield early memory B cells. We also find that a marked global reduction in host chromatin accessibility occurs during the first stages of infection in subpopulations of EBV^+^ cells that display senescent and pre-apoptotic hallmarks induced by innate antiviral sensing and proliferation-linked DNA damage. However, cells in proliferative infection trajectories exhibit greater accessibility at select host sites linked to B cell activation and survival genes as well as key regions within the viral genome. To further investigate such loci, we implement a bioinformatic workflow (crisp-ATAC) to identify phenotype-resolved regulatory signatures. This customizable method applies user-specified logical criteria to produce genome-wide single-cell ATAC-and ChIP-seq range intersections that are used as inputs for *cis*-linkage prediction and ontology tools. The resulting tri-modal data yield exquisitely detailed hierarchical perspectives of the transforming regulatory landscape during critical stages of an oncogenic viral infection that simulates antigen-induced B cell activation and differentiation. We anticipate these resources will guide investigations of gene regulatory modules controlling EBV-host dynamics, B cell effector fates, and lymphomagenesis. To demonstrate the utility of this resource, this work concludes with the discovery of EBV infection dynamics in FCRL4^+^ / TBX21^+^ Tissue-Like Memory B cells, an unconventional subset with notable associations to numerous immune disorders.

## Introduction

Epstein-Barr Virus (EBV) is an oncogenic gammaherpesvirus present in >90% of adults (Rickinson and Kieff, 2007) and associated with up to 2% of human cancers (Cohen et al., 2011). Recent reports have also provided epidemiological and mechanistic evidence supporting an etiological role for EBV in multiple sclerosis (MS) (Bjornevik et al., 2022; Lanz et al., 2022). In its initial stages, EBV infection within primary B lymphocytes manifests an array of host and viral programs. Upon entry into the host cell, the linear dsDNA viral genome rapidly circularizes to form an episome that is retained within the nucleus (Lindahl et al., 1976; Nonoyama and Pagano, 1972). Within hours to days, host innate immune responses are generated to restrict viral progression (Lünemann et al., 2015; Martin et al., 2007; Smith et al., 2013; Tsai et al., 2011). Simultaneously, viral genes are expressed to counteract host defenses (Ressing et al., 2015), co-opt B cell-intrinsic activation and proliferation (Calender et al., 1987; Thorley-Lawson, 2001; Thorley-Lawson and Mann, 1985), and attenuate DNA damage and stress responses instigated by virus-induced growth (McFadden et al., 2016; Nikitin et al., 2010). A consequence of these intimately adapted host-pathogen dynamics is that EBV infection can precipitate diverse responses and outcomes for host B cells. These include unsuccessful infection routes resulting from effective antiviral restriction and DNA damage-induced growth arrest as well as successful infection leading to immortalization *in vitro* (Bird, 1981; Henle et al., 1967; Pope et al., 1968; Zhao et al., 2011) or lifelong latency *in vivo* within memory B cells (Babcock, 1998; Longnecker et al., 2013; Miyashita et al., 1997) that retain oncogenic potential (Raab-Traub, 2007; Thorley-Lawson and Gross, 2004).

Since its discovery in 1964 as the first human tumor virus (Epstein et al., 1964; Young and Rickinson, 2004), extensive research has revealed the molecular means by which EBV establishes infection and underlies various malignancies. The entire EBV genome is ∼172 kilobases and contains at least 80 protein-coding sequences including six EBV nuclear antigens (EBNAs); several latent membrane proteins (LMPs); and loci that encode replicative and transcriptional machinery as well as structural proteins. The EBV genome also contains functional non-coding RNAs: the BHRF and BART microRNAs and the EBV-encoding regions (EBERs) (Rickinson and Kieff, 2007; Young et al., 2007).

EBNAs are especially important in establishing distinct forms of latency depending on their combinatorial expression (Price and Luftig, 2015). EBNA1 is a transcription factor (TF) that is essential for viral genome maintenance and B cell transformation and ubiquitously binds and epigenetically regulates host chromatin (Altmann et al., 2006; Canaan et al., 2009; Dheekollu et al., 2021; Humme et al., 2003; Lamontagne et al., 2021; Lu et al., 2010; Lupton and Levine, 1985; Wood et al., 2007; Yates et al., 1985). EBNALP is another essential factor (Mannick et al., 1991; Szymula et al., 2018) that initiates host cell proliferation alongside its co-activated target, EBNA2 (Alfieri et al., 1991; Harada and Kieff, 1997; Sinclair et al., 1994), and interacts with several host proteins including TFs (Han et al., 2001; Ling et al., 2005; Matsuda et al., 2003). EBNA2 is likewise required for B cell immortalization (Cohen et al., 1989), notably through coordination with host TFs and their binding sites (Lu et al., 2016; Zhao et al., 2011) and with EBNALP to drive early cell proliferation and viral LMP1 expression (Peng et al., 2005). The EBNA3 proteins (EBNA3A, EBNA3B, and EBNA3C) mediate a delicate balance of anti-and pro-oncogenic processes (Allday et al., 2015; Banerjee et al., 2014; Parker et al., 1996; Tomkinson et al., 1993; White et al., 2010). These include epigenetic repression of host tumor suppressor genes (*BIM, p14, p16*) and viral promoters (Maruo et al., 2011; Paschos et al., 2012; Saha et al., 2015; Skalska et al., 2010; Styles et al., 2017), competitive binding of the EBNA2-interacting host factor RBPJ (Robertson et al., 1995; Wang et al., 2015), and inhibition of apoptosis (Price et al., 2017). Collectively, the EBNAs reshape the nuclear regulatory and transcriptional landscape of EBV^+^ B cells, effectively hijacking B cell-intrinsic activation, expansion, and differentiation programs. Thus, EBV co-opts antigen-responsive host immune mechanisms for the ulterior purposes of viral replication and propagation.

While the EBNAs engage cell proliferation machinery at the epigenetic and transcriptional level in the nucleus, the LMPs (LMP1, LMP2A, and LMP2B) do so at the cell membrane by simulating antigen-induced signal transduction pathways. The essential LMP1 promotes B cell activation through mimicry of a constitutively active CD40 receptor (Kilger et al., 1998; Uchida et al., 1999) and interacts with Tumor Necrosis Factor (TNF) receptor-associated factors (TRAFs) to activate NF-κB pathway signaling via IKK (Devergne et al., 1996; Eliopoulos et al., 2003; Greenfeld et al., 2015; Luftig et al., 2003). These interactions induce anti-apoptotic pathways, MHC-mediated immune recognition, pro-inflammatory responses, and cell migration. Downstream consequences include oncogenic proliferation and survival but also induction of pro-apoptotic responses (Devergne et al., 1998; Fries et al., 1999; Greenfeld et al., 2015; Henderson et al., 1991; Shair et al., 2008; Wang et al., 2017). Thus, as in antigen-induced B cell activation (and subsequent differentiation), adept regulatory control of NF-κB signaling (Hoffmann et al., 2002; Mitchell et al., 2018; O’Dea et al., 2007; Roy et al., 2019) is dispositive for the fate of a given EBV^+^ B cell. Although it is not essential for transformation, LMP2A promotes cell survival through mimicry of a stimulated B cell receptor (BCR), which activates signaling cascades complementary to those induced by LMP1 (Anderson and Longnecker, 2008; Fish et al., 2020; Guasparri et al., 2008; Portis and Longnecker, 2004). LMP2A expression further predisposes EBV^+^ B cells to survival by lowering antigen affinity selection thresholds *in vivo* (Minamitani et al., 2015). Thus, EBV latent membrane proteins play integral roles in B cell proliferation in the absence of antigen licensing and in avoiding replicative dead ends effected by antiviral sensing.

Clearly, key EBV gene products manipulate diverse host programs at early stages to achieve sustained latency (Mrozek-Gorska et al., 2019; Pich et al., 2019). Many such perturbations involve extensive rewiring of epigenetic and transcriptional regulatory modules. EBV researchers have used methods such as RNA-, ATAC-(Assay for Transposase-Accessible Chromatin), and ChIP-seq (Chromatin Immunoprecipitation) to study these changes at various levels in the gene regulatory hierarchy within early infected cells and transformed lymphoblastoid cell lines (LCLs) (Arvey et al., 2012; Jiang et al., 2017; McClellan et al., 2013; Mrozek-Gorska et al., 2019; Wang et al., 2019; Zhou et al., 2015). Recently, the epigenetic and transcriptional roles of EBNA1 were interrogated through time-resolved multi-omics (Lamontagne et al., 2021). While these and other studies provide indispensable insights regarding virus-induced genome-wide expression and regulation, they have relied on bulk ensemble sequencing. Such assays yield population-averaged measurements that obscure variation arising from intrinsic stochasticity (Raj et al., 2006; Raj et al., 2010; Raj and Van Oudenaarden, 2008), asynchronous behaviors, and heterogeneous cell subsets. Specifically, ensemble averaging fails to capture cell-matched measurements across genes, which precludes identification of coordinated expression programs or epigenomic regulatory patterns in specific phenotypes. By contrast, single-cell sequencing provides refined genome-wide views of expression and regulation that preserve the ability for the identification of heterogenous cell states with low bias (Buenrostro et al., 2015; Junker and van Oudenaarden, 2014; Shalek et al., 2013; Shapiro et al., 2013; Wills et al., 2013). Given the complexity of host-virus relationships, single-cell -omics approaches are essential to dissect the early stages of EBV infection and the distinct fate trajectories it comprises. We previously used single-cell RNA sequencing (scRNA-seq) to identify EBV-driven heterogeneity in LCLs (SoRelle et al., 2021). Recent advances in single-cell multimodal-omics methods have made it possible to integrate scRNA-seq with several levels of hierarchical regulation (Efremova and Teichmann, 2020), which can provide greater insight into the mechanistic origins underlying gene expression. These include techniques for obtaining cell-matched measurements of mRNA transcripts, chromatin accessibility, and DNA methylation status (Cao et al., 2018; Chen et al., 2019; Clark et al., 2018; Zhu et al., 2019), as well as other molecular levels. In this work, we leverage single-cell multiomics (scRNA-seq + scATAC-seq) to capture and explore the distinct gene expression and regulatory signatures that determine the course of EBV infection in primary human B lymphocytes.

## Results

### EBV asynchronously induces primary B cells into distinct phenotypic states early after infection

To interrogate chromatin accessibility and gene expression changes that occur upon EBV infection, we isolated primary human B cells from the peripheral blood of two donors and infected them with the B95-8 strain of EBV. Infections were performed at a multiplicity of infection (MOI) of 5 to ensure latent gene expression in every cell (Nikitin et al., 2010). We cryopreserved samples of infected cells at 2-, 5-, and 8-days post-infection in addition to uninfected cells (Day 0) from each donor sample following B cell enrichment. Cell samples from each donor and timepoint were simultaneously thawed, prepared to >90% viability, and processed into single-cell multiome libraries. Single-cell matched transcript and accessibility data were obtained through standard NGS, alignment, counting, and quality control (QC) methods (**Table S1**).

EBV infection induced broad transcriptomic changes in B lymphocytes at high efficiency, as evidenced by the near-complete loss of resting phenotypes (Day 0) within two days of infection. New states emerged between Day 2 and Day 5, while subtle shifts in state proportions defined the period between Day 5 and Day 8 (Figure 1A). Total and unique transcripts per cell increased, particularly between Day 0 and Day 2, while mitochondrial gene expression increased gradually (Figure 1B). Total transcript and mitochondrial distributions at Day 2 exhibited two modes, which was consistent with the presence of both non-proliferative and mitotic cells identified by S-phase and G2M-phase marker scoring (Figure 1C).

**Figure 1.**
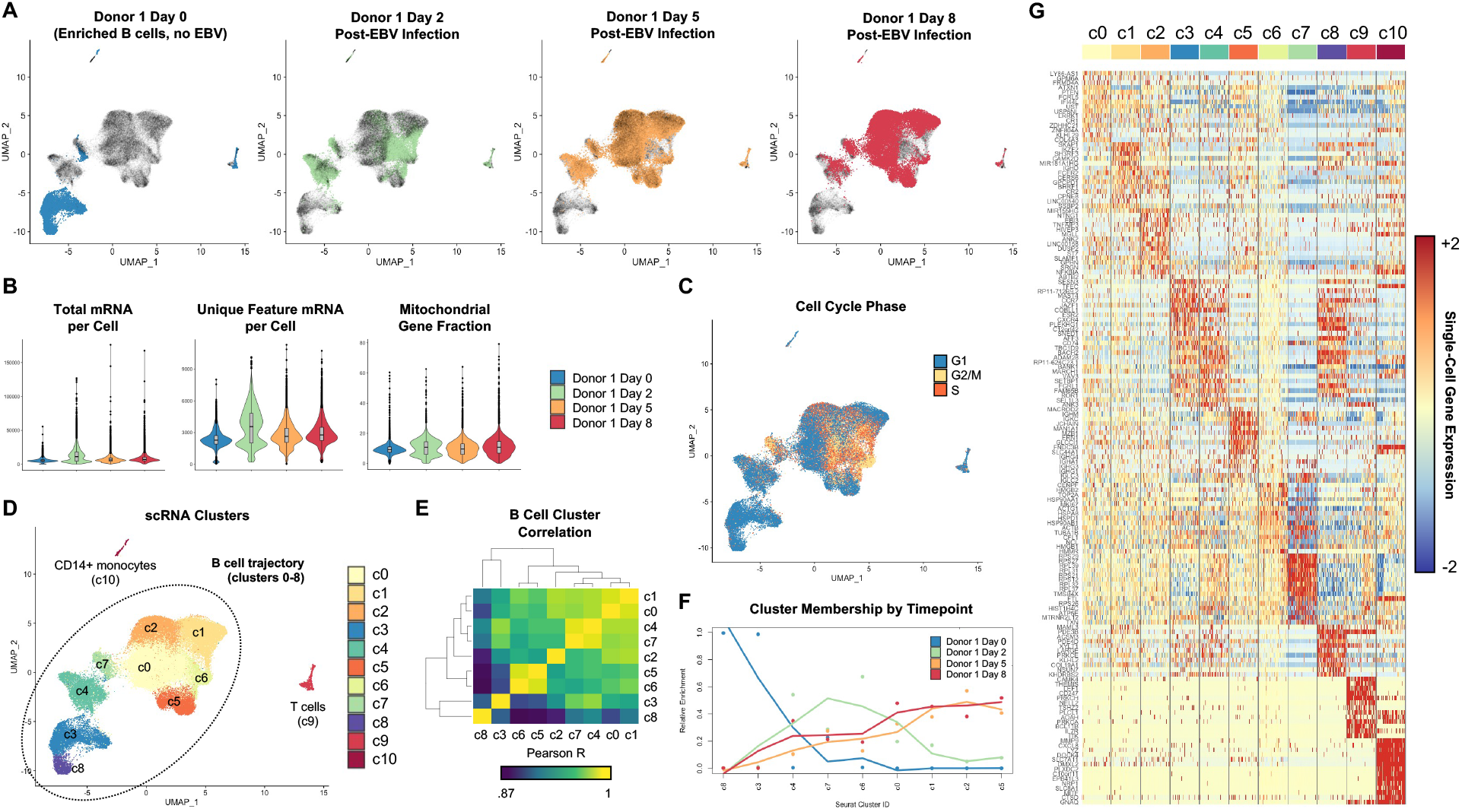
Time-resolved single-cell gene expression during early EBV infection of B lymphocytes. (A) Single-cell gene expression timecourse data from early EBV infection. (B) General expression trends during early infection. Total mRNA refers to all transcripts captured, while Feature mRNA refers to the number of unique transcripts (per cell). (C) Cell phase scoring of expression data after cell cycle marker regression. (D) Unsupervised clustering of early infected cell expression in merged timepoint data. (E) Pairwise correlation of identified clusters. (F) Cluster membership by timepoint. Fit lines show coarse changes in phenotype frequency over time. (G) Single-cell expression of the top 15 gene markers by cluster. *See also Figures S1-S4*

Unsupervised methods revealed subpopulations (clusters) in cell cycle-regressed aggregated scRNA-seq time courses. Two clusters corresponded to uninfected B cells (c3, c8); seven were post-infection B cell phenotypes (c0, c1, c2, c4, c5, c6, c7); and two were T cells (c9) and CD14^+^ monocytes (c10) carried over from PBMCs despite extensive B-cell enrichment (Figure 1D). Genome-wide expression correlation was higher among post-infection states relative to uninfected cells, and certain phenotypes were more strongly correlated (e.g., c0 with c1; c4 with c7, Figure 1E). Sorting cluster membership by day yielded coarse-grained dynamics of cell state transitions (Figure 1F). We determined top differential genes in each cluster (one-vs-all-others) to inform state identity annotations (Figure 1G). Identified clusters included many genes known to be modulated in EBV infection and were broadly consistent across both donors with respect to top marker genes, cell population frequencies, and temporal emergence (**Figures S1-S4**).

### Infected cell state heterogeneity is linked to antiviral and B cell-intrinsic responses

Cluster analysis deconvolved heterogeneous biological states within each sample and revealed phenotypes retained across multiple timepoints (Figure 2). At this resolution, we estimated time- and state-level trends in viral gene expression, variation in metabolic activity, and transcript diversity (Figure 2A). Overall, c0, c1, c2, c5, and c6 exhibited the highest levels of EBV transcripts and more unique transcripts than c3, c4, c7, and c8. Mitochondrial gene fraction and unique feature content were highest in c6 and lowest in c4 and c7, although c7 had a long-tailed distribution of mitochondrial expression (20-80%) prior to QC, indicative of (pre-) apoptotic cells. All clusters except uninfected B cells (c3, c8) displayed broad innate antiviral and interferon-stimulated gene (*ISG* and *IFI* member) expression. Antiviral gene expression was generally higher and exhibited greater variance in c7 than c4 and persisted at roughly uniform levels in c0, c1, c2, c5, and c6 (Figure 2B). Through joint consideration of cluster-resolved expression trends for viral, mitochondrial, and interferon-stimulated genes, we distinguished uninfected cells (c3, c8) and two classes of cells with the hallmarks of antiviral response: those with low proliferation and negligible viral expression (c4, c7) and those with viral and metabolic indicators of progressive EBV infection (c0, c1, c2, c5, c6).

**Figure 2.**
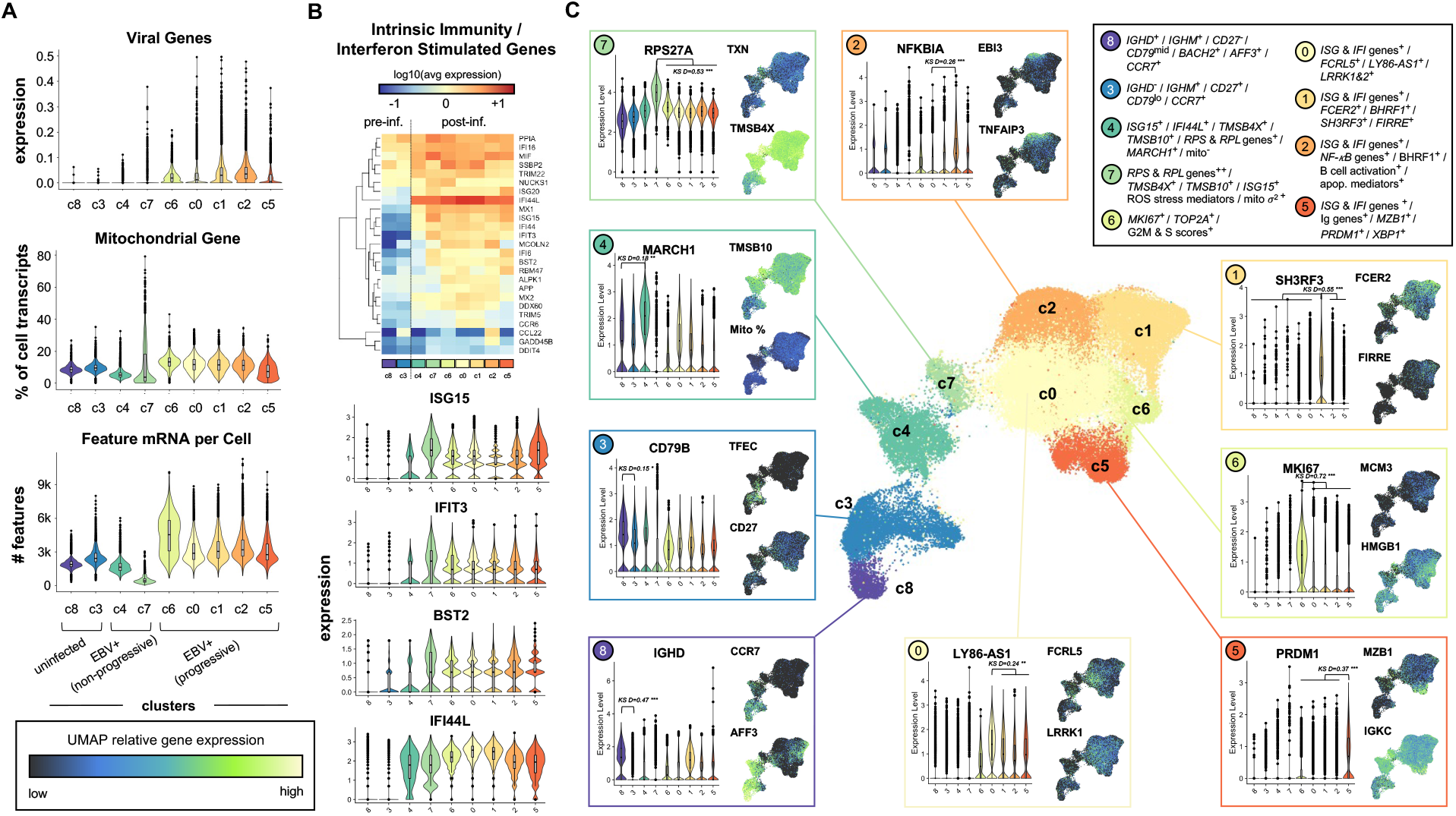
High-resolution dissection of infected B cell phenotypes. (A) Overview of global gene expression trends by phenotype. (B) Induction of interferon response genes in all EBV^+^ clusters. (C) Phenotype-resolved transcriptomic signatures in resting and EBV^+^ B cells. Select cluster-resolved comparisons of gene expression were evaluated via the Kolmogorov-Smirnov D statistic (KS D) and associated p value (* p < 1e-5; ** p < 1e-10; *** p < 1e-15) from 500 randomly sampled cells per cluster. *See also Figures S5-S11*

Next, we extensively analyzed differentially expressed genes (DEGs) among clusters and groups, including pairwise comparisons of all post-infection phenotypes (Figures 2C**, S5-S11**). The two resting cell phenotypes differed in their expression of *IGHD*, *IGHM*, *CD27*, and other markers that distinguish naïve (c8) from memory (c3) B cells. In addition to interferon response signatures, non-proliferating infected cells exhibited an overall reduction in gene expression and upregulated stress response markers. These included the highest overall expression of actin sequestration genes (*TMSB10*, *TMSB4X*) and, particularly within c7, numerous ribosomal subunit genes (e.g., *RPS27A*). Cells in c4 were distinguished by elevated expression of *MARCH1*, which encodes an E3 ubiquitin ligase that regulates the type I interferon response (Wu et al., 2020). Unlike c4, c7 cells also contained high transcript levels for genes involved in oxidative stress (*TXN*, *FTL*, *FTH1*), cytochrome oxidase subunits (e.g., *COX7C*), ubiquitin genes (*UBA52*, *UBL5*) and highly variable mitochondrial fractions. Among EBV^+^ cells with hallmarks of elevated respiration, those in c6 were most clearly consistent with proliferating cells based on upregulated cell cycle markers. Cells in c0 were distinguished by upregulation of *FCRL5* and *LY86-AS1*, an antisense RNA to a lymphocyte antigen (*LY86*) that mediates innate immune responses. Cells in this cluster also displayed markers consistent with the early stages of pre-germinal center activated B cells (e.g., *CCR6, CD69, POU2AF1, TNFRSF13B, PIK3AP1*). Notably, cells in c1 and c2 contained the highest levels of the EBV gene *BHRF1*. Between these two phenotypes, c2 was enriched for genes involved in NF-κB signaling and known markers of EBV-mediated B cell activation (*NFKBIA*, *TNFAIP3*, *EBI3*) while c1 appeared to be derived from naïve cells (based on *IGHD* and other carryover genes) and exhibited near-unique expression of *SH3RF3/POSH2* and *FIRRE*, a MYC-regulated long non-coding RNA (lncRNA). Finally, c5 displayed upregulation of immunoglobulin heavy and light chains (*IGHA1, IGHG1, IGHM, IGKC, IGLC1-3*) as well as genes involved in B cell differentiation (*MZB1, PRDM1/BLIMP1, XBP1*). Gene ontology (GO) networks were also generated for top DEGs from one-versus-all-other comparisons to facilitate phenotype annotations (**Figures S12-S16**).

### A map of B cell phenotypes and fate trajectories in early EBV infection

Graph-based pseudotime (Qiu et al., 2017) approximated EBV-induced state transitions when anchored from resting cells (Figure 3A). Pseudotime scoring was used to track state dynamics of the top 25 marker genes for each phenotype and four example expression trajectories are highlighted (Figure 3B). Collectively, flow cytometry for the B cell marker CD19 and CD23 (upregulated in EBV infection) at each timepoint (**Figure S17**), cluster-specific DEGs, network ontologies, and pseudotime led us to propose a multi-phenotype model for heterogeneous cell fate trajectories (Figures 3C**, S18-S19**) that manifest in early EBV infection *in vitro*. In this model, naïve (c8) and memory (c3) B cells infected with EBV either undergo antiviral response-mediated arrest (c4) or EBV-driven hyperproliferation (c6) within several days of infection. Hyperproliferating cells can subsequently enter one of several activated states (c0, c1, c2) or undergo growth arrest (c7). Further, differentiated B cells (c5) can develop following activation in a manner analogous to effector cell exit from the germinal center reaction. Among activated phenotypes, c2 matched classical EBV-mediated activation of NF-κB pathway genes, apoptotic regulators, and other known biomarkers (Cahir-McFarland et al., 2004; Messinger et al., 2019). Cells in c1 were consistent with a related activation intermediate that originated from EBV^+^ naïve cells. Despite the relatively short timecourse, c2 and c5 began to reflect the continuum of activation and differentiation phenotypes we previously characterized in LCLs (SoRelle et al., 2021), which are considered to be immortalized at 21-28 days post-infection (Nilsson et al., 1971). We confirmed these similarities by merging Day 8 and the LCL GM12878, for which scRNA-seq data was previously reported and analyzed (Osorio et al., 2019; Osorio et al., 2020; SoRelle et al., 2021) (**Figure S20**). Conceivably, EBV^+^ cells could also transition to a plasmablast phenotype (c5) from memory cells (c3) through hyperproliferation (c6) via division-linked differentiation (Hodgkin et al., 1996), effectively bypassing intermediate states.

**Figure 3.**
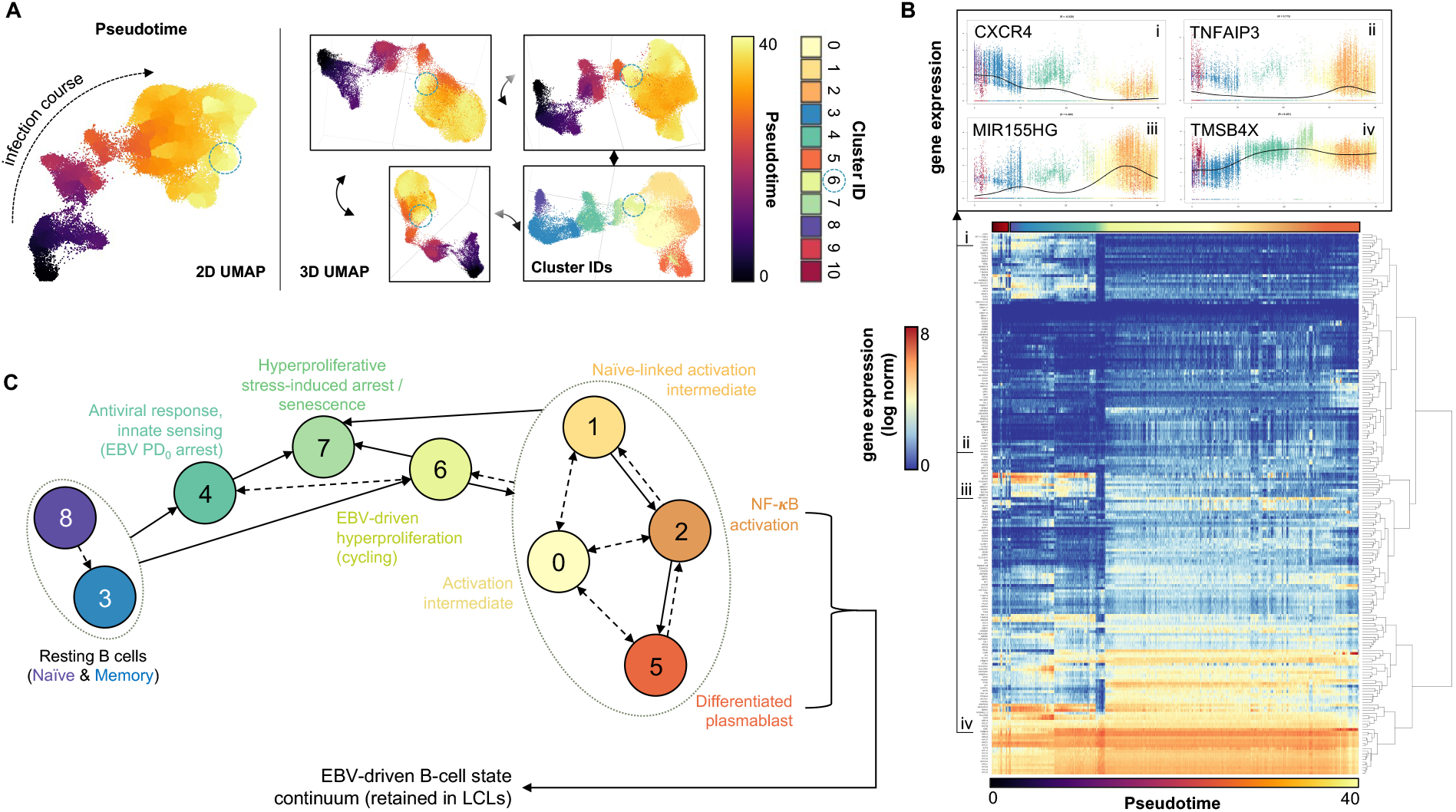
A model of B cell fate trajectories in early EBV infection. (A) Monocle3 pseudotime scoring of merged timecourse expression data relative to resting B cells (day 0). Unlike the 2D UMAP, 3D UMAPs depict closer proximity of c6 (first observed at day 2, blue dashed circle) to resting cells, consistent with the temporal emergence of the c6 phenotype prior to the c0, c1, c2, and c5 phenotypes. (B) Pseudotime-resolved expression dynamics of top differentially expressed genes (DEGs) across phenotypes. Genes are hierarchically clustered by pseudo-temporal expression pattern similarity. Spline interpolant fits are shown for expression of select genes in pseudotime (insets i-iv). After sorting cells by pseudotime score in ascending order, the average pseudotime score of every 25-cell interval was computed for efficient visualization (i.e., pseudotime for 52,271 cells at 25 cell resolution). (C) Annotated state model of EBV^+^ B cell phenotypes and fate trajectories. Empirically observed and putative directed state transitions are depicted in solid and dashed edges, respectively. Edges drawn to groups of phenotypes (dotted ovals) indicate transitions to/from each cluster within the group. *See also Figures S12-S19*

Conversely, cells in c7 highlighted diverse origins of EBV^+^ cell growth arrest, apoptosis, and senescence, which each provide host defenses against oncogenic malignancies (Bartkova et al., 2006; Nikitin et al., 2010). In addition to highly variable mitochondrial expression and the lowest transcript levels of any state, this phenotype was defined by broad upregulation of genes involved in ribosome biogenesis-mediated senescence (*RPS14*, *RPL29*, *RPS11*, *RPL5*) (Lessard et al., 2018; Nishimura et al., 2015) and stress-associated sequestration of actin monomers that favor G-actin formation (*TMSB4X*, *TMSB10*, *PFN1*) (Kwak et al., 2004). A subset of cells within c7 also contained elevated levels of cell cycle markers (*MKI67*, *TOP2A*, *CCNB1*, *CENPF*) carried over from pre-arrest hyperproliferation (**Figure S21**).

### Evidence for EBV induction of an activated precursor to early memory B cells (AP-eMBC)

We next sought to compare early infected phenotypes from our multistate model with cells isolated from secondary lymphoid organs. We acquired single-cell RNA-seq data from human tonsil tissue and identified germinal center (GC) cell subsets (Figure 4A), which we analyzed alongside early infection phenotypes of interest. EBV^+^ NF-κB activated cells (c2) clearly mimicked GC light zone (LZ) B cells; *MKI67*^hi^ cells (c6) matched actively cycling cells (including GC dark zone (DZ) B cells); and EBV^+^ differentiated cells (c5) matched plasmablasts and plasma cells (PB/ PC). Cells in c0 were most like pre-GC naïve and memory B cell (MBC) subsets (Figure 4B-D). Further, numerous c0 markers were consistent with both pre-GC activated B cells (*SELL, BANK1, CD69, GPR183 (EBI2)*) and memory B cell phenotypes (*SELL, BANK1, GPR183, PLAC8*) recently identified from scRNA-seq of tonsils in response to antigen challenge (King et al., 2021) (Figure 4C-D). Cells in c0 further exhibited upregulation of genes with essential roles in B cell activation (*TNFRSF13B/TACI*) (Wu et al., 2000) and germinal center formation (*POU2AF1/OCA-B*) (Kim et al., 1996; Luo and Roeder, 1995; Schubart et al., 1996) (**Figure S18**). Moreover, c0 displayed elevated *CCR6*, a marker of an activated precursor (AP) state that can generate early memory B cells (eMBCs) (Glaros et al., 2021; Suan et al., 2017) (Figure 5A).

**Figure 4.**
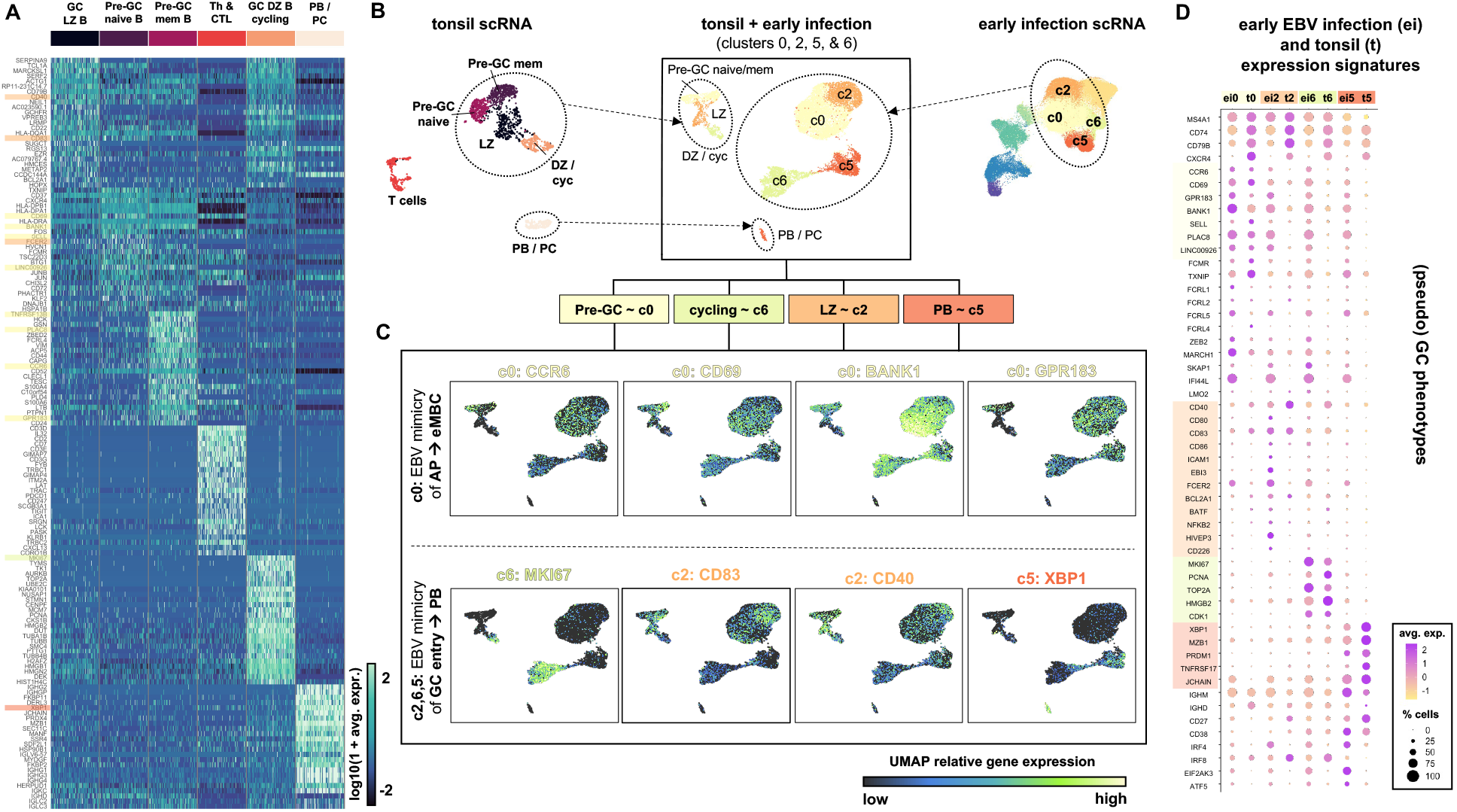
A subset of early infected cells exhibits hallmarks of a multipotent activated precursor to early memory B cells (eMBCs) (A) Top phenotype markers of healthy human tonsil subsets identified from scRNA-seq. (B) UMAP merging of tonsil and key early infection cluster scRNA-seq assays. Tonsil clusters are colored to match the closest corresponding cells from early infection. (C) UMAP Correspondence of key gene expression across tonsillar subsets and early infection phenotypes. Select markers of multipotent progenitors and eMBCs were informed by data from (Suan et al., 2017) and (Glaros et al., 2021). (D) Dot plot visualization of key genes across early infection (ei) c0, c2, c5, and c6 and their analogs within tonsils (t).

**Figure 5.**
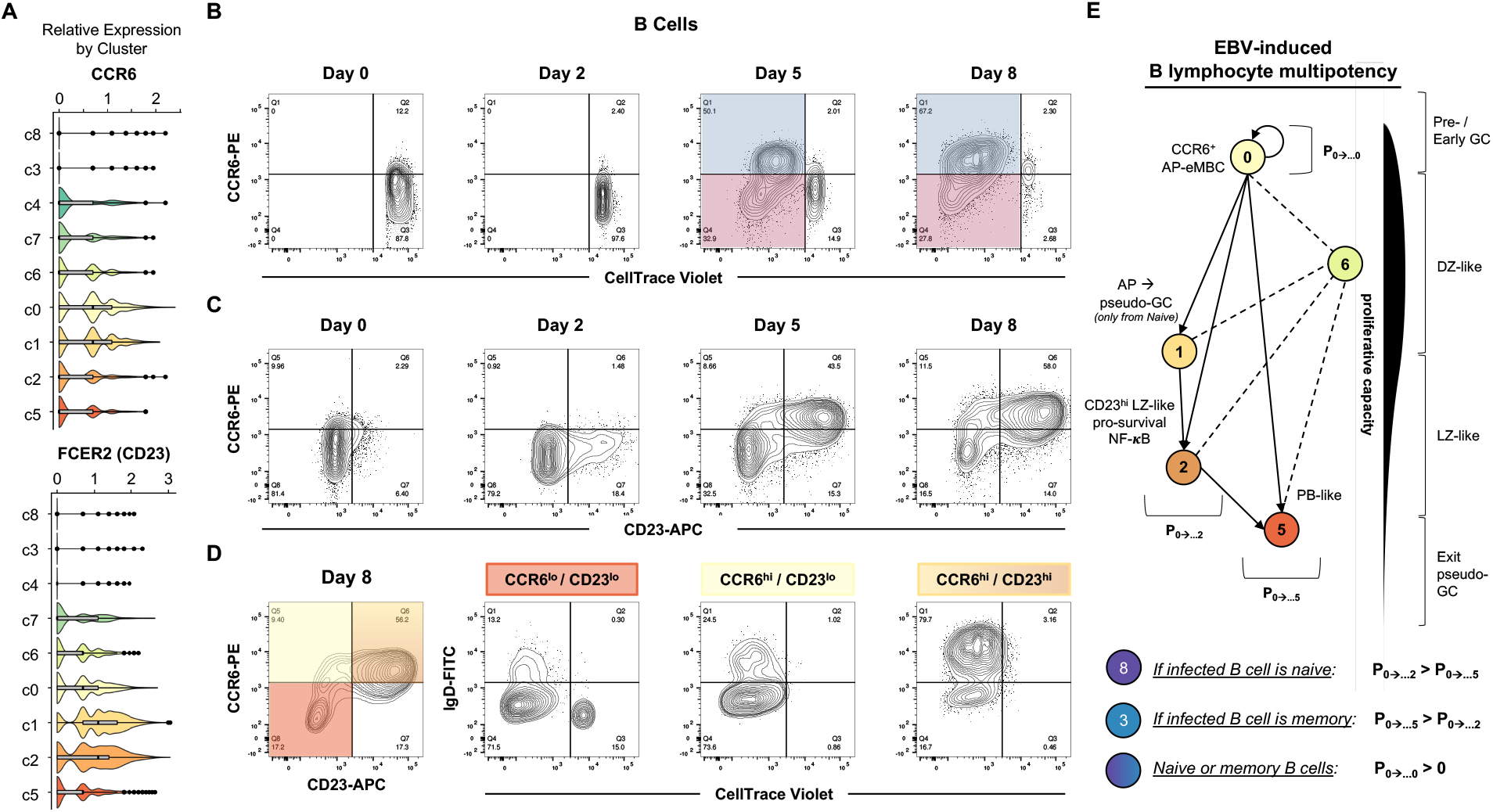
FACS validation of CCR6^+^ AP-eMBCs derived from naïve and memory B cells. (A) Relative expression of *CCR6* and *FCER2/CD23* by model phenotype determined by scRNA-seq. (B) CCR6 surface expression on uninfected and EBV-infected B cells by number of cell divisions (CellTrace Violet). CCR6^hi^ cells (blue gate) exhibits reduced proliferation relative to CCR6^lo^ (magenta gate) cells. (C) CCR6 and CD23 (FCER2) surface expression over the early infection timecourse. (D) Cell divisions and IgD status of cells at Day 8 gated by CCR6 and CD23 expression. Gated fractions are colored by approximate correspondence to scRNA-seq model phenotypes. (E) A fate model for EBV-induced AP-eMBC-like cells. P0à…n signifies the probability of the eventual transition of a cell from cluster 0 (AP-eMBC analog) to a given cluster n. Relative probability relationships for naïve and memory cells are proposed based on empirical findings from FACS. *See also Figures S22-25*

We subsequently validated the generation of CCR6^+^ AP-eMBC B cells in response to EBV infection through time-resolved FACS (Figures 5B-D**, S22-S24**). Resting B cells were CCR6^lo^ and remained so until between 2 and 5 days after infection. Further, we observed that the most proliferative cell fraction at day 8 was CCR6^lo^ and a moderately proliferative cell population was CCR6^hi^. While the most proliferative cells were CCR6^lo^/CD23^lo^, the proliferative CCR6^hi^ cells displayed variable CD23 levels (Figure 5B). Consistent with our scRNA time course, CCR6^hi^/CD23^hi^ and CCR6^hi^/CD23^lo^ populations respectively corresponded to c1/c2 and c0 and emerged within 5 days (Figure 5C). Based on CD27 and IgD status, these populations predominantly originated from naïve or non-switched memory versus switched memory cells, respectively; notably, cells from these different resting phenotypes were present in each population gated by CCR6 and CD23 status (Figure 5D**, S24C**). Rapidly proliferative CCR6^lo^/CD23^lo^/CD27^hi^/IgD^lo^ cells were consistent with infected memory B cells transitioning to plasmablasts (*c3àc6àc5* model trajectory; ∼72% of CCR6^lo^/CD23^lo^ cells). Marginally less proliferative CCR6^hi^/CD23^lo^/CD27^hi^/IgD^lo^ cells were consistent with stimulated AP-eMBCs (*c3àc6àc0* model trajectory; ∼74% of CCR6^hi^/CD23^lo^ cells). We also observed an IgD^hi^ naïve population that matched the pre-GC AP-eMBC phenotype (*c8àc6àc0*; (∼25% of CCR6^hi^/CD23^lo^ cells). Finally, an even less proliferative CCR6^hi^/CD23^hi^/IgD^hi^ population matched activated naïve (or non-switched memory) cells destined for GC BC (*c8àc6/c0àc1/c2*; ∼80% of CCR6^hi^/CD23^hi^ cells) and the minor subset (∼17%) of CCR6^hi^/CD23^hi^ cells that was IgD^lo^ was consistent with MBCs induced by EBV to undergo a pseudo-GC reaction (*c3àc6/c0àc2*) (Figure 5C-E). Intriguingly, a subset of CCR6^+^ cells displaying the AP phenotype apparently persists long after the early stages of infection based on scRNA-seq data from LCLs (**Figure S25**). Thus, c0 in our model matches a virus-induced common progenitor state from which PBs, GC BCs, and early MBCs have been shown to originate in response to antigen stimulation (Taylor et al., 2015). Our results further indicate that both naïve and memory B cells can achieve this multipotent state at different frequencies upon *in vitro* infection and that the AP-eMBC phenotype is perpetuated in EBV-immortalized B cells.

### Linked expression and accessibility illuminate regulatory mechanisms in phenotype trajectories

We next investigated potential regulatory mechanisms underlying DEGs observed across phenotypes. Expression data were jointly analyzed with cell-matched measurements from single-cell Assay for Transposase-Accessible Chromatin sequencing (scATAC-seq) and annotated by state (Figures 6A-B, **S26A-B**). While total and unique transcripts per cell increased through early infection, global chromatin accessibility decreased substantially upon infection. Resting (c3, c8) and hyperproliferative (c6) cells had the highest overall accessibility. There were significantly more peaks in the NF-κB activation state (c2) relative to other activation intermediates (c0 and c1; two-tailed t-test, p < 2.2×10^-16^ and 1.4×10^-14^, respectively) and differentiated cells (c5; p < 2.2×10^-16^). Similar accessibility reduction occurred in both donors in the first five days, with increased accessibility recovered between Day 5 and Day 8 (including to higher than resting levels in one donor) (Figures 6B**, S27**). This indicated that EBV-induced heterochromatinization is likely transient in successfully infected cells (i.e., those that evade innate- and damage-mediated arrest).

**Figure 6.**
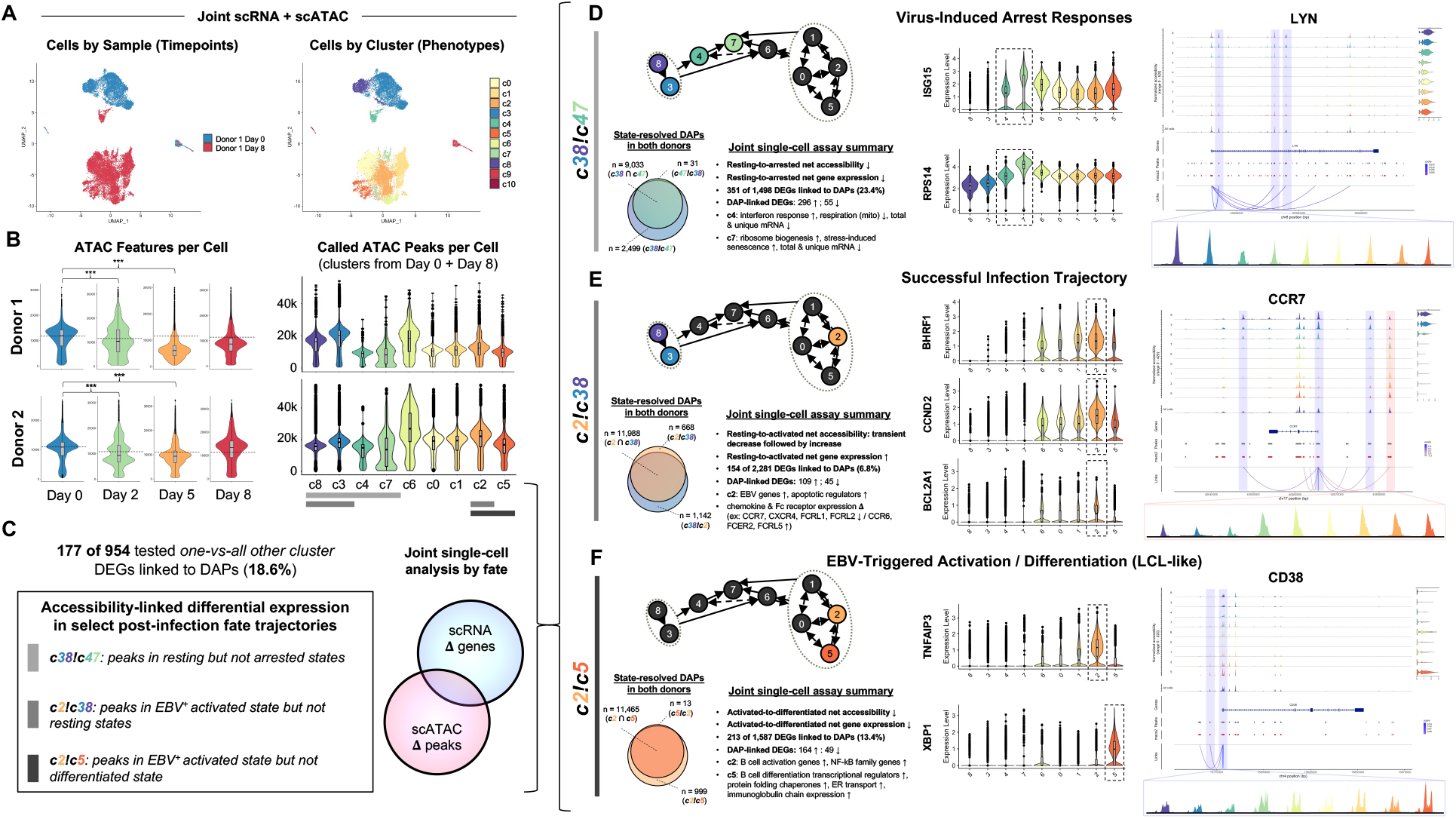
Cell-matched expression and chromatin accessibility cell fate trajectories. (A) UMAP visualization of scRNA + scATAC data generated using weighted nearest neighbors (WNN) multimodal integration (Hao et al., 2021; Stuart et al., 2020). Merged multimodal data from the first and last timepoints (day 0 & day 8) contain cells representative of all identified phenotypes. (B) Distribution of called ATAC peaks per cell by timepoint and phenotype in both donors (***p < 1e-15, one-sided Kolmogorov-Smirnov test). (C) Overview of global differentially accessible peak (DAP)-linked differentially expressed genes (DEGs) and cluster comparisons for major trajectories of interest. DAPs are identified by their presence in one or more peaks but not in (!) one or more other peaks. DAP-linked DEGs were explored in resting versus arrested cells (c38 vs c47), EBV^+^ activated versus resting cells (c2 vs c38), and EBV^+^ activated vs EBV+ differentiated cells (c2 vs c5). (D) Virus-induced arrest responses. State-resolved DAPs (*c38!c47*) and joint assay trend summaries are presented in addition to example DEGs and DAP linkages resolved by phenotype. (E) Successful infection trajectory. State-resolved DAPs (*c2!c38*), joint assay summaries, and trajectory-specific examples are presented as in (D). (F) EBV-induced B cell activation/differentiation continuum. State-resolved DAPs (*c2!c5*), joint assay summaries, and trajectory-specific examples are presented as in (D and E). *See also Figures S26-S38*

We found 954 linked feature genes derived from the top 100 marker genes for each cluster using the multimodal integration capabilities of the Signac package in R (Stuart et al., 2021). Of these 954 genes, 177 were significant DEGs with linkages to 476 differentially accessible peaks (DAPs). This translated to 18.6% of tested genes with potential DAP-linked regulation (Figures 6C**, S26C-D**). We identified genes linked (in *cis*) to DAPs to explore phenotype-associated gene regulatory relationships (**Figure S28A**). Joint analysis of DAP-linked DEGs yielded four regulatory patterns: higher accessibility with higher expression (+/+); lower accessibility with higher expression (-/+); lower accessibility with lower expression (-/-); and higher accessibility with lower expression (+/-). (**Figure S28B**). The +/+ and -/-patterns were characteristic of positive regulatory sites. The less frequently observed -/+ and +/- patterns were consistent with closure or opening of ATAC sites with negative regulatory functions, respectively. Analysis of genes of interest including *CCR7*, *CXCR4*, *RUNX3*, *BACH2*, *JCHAIN*, and *PRDM1* provided examples of each regulatory pattern and their variation among states (**Figures S28C, S29-S30**).

We developed joint scRNA + scATAC profiles for major infection fate trajectories in our model (Figure 6C-F). The path from resting cells to EBV^+^ arrested/senescent cells (*c38àc47*) was characterized by global reductions in accessibility and expression. 34.3% of all DEGs between resting cells and these non-proliferative EBV^+^ fates were linked to DAPs that become inaccessible after infection (*c38!c47* peaks). Top DAP-linked DEGs in innate arrested cells (c4) corresponded to upregulation of interferon-responsive genes and downregulation of mitochondrial genes, while stress- and damage-induced senescent cells (c7) were distinguished by their high expression of interferon-stimulated genes (e.g., *ISG15*) and ribosomal transcripts (e.g., *RPS14*). Cells in each of these clusters also displayed reduced expression of the proto-oncogenic tyrosine kinase gene *LYN* linked to closure of multiple regulatory sites following infection (Figure 6D).

The reduction in ATAC peaks within c7 was consistent with the formation of senescence-associated heterochromatin foci (SAHF) (Courtois-Cox et al., 2008; Di Micco et al., 2011; Lenain et al., 2017) (**Figure S31**). Because senescence can arise from diverse mechanisms such as innate immune sensing or growth-induced DNA damage, we used higher resolution clustering to reveal c7 subsets (7a and 7b). These subsets displayed DEGs involved in the cell cycle and antiviral sensing (**Figure S32**). Different *HMGB2* levels between 7a and 7b were notable, as this gene’s product mediates diverse roles in sensing (Yanai et al., 2009), double-stranded break repair (Krynetskaia et al., 2009), and p53 downregulation (Stros et al., 2002). Relative to resting B cells, *HMGB2* expression was strongly elevated in 7b (as in the hyperproliferative state, c6) but only mildly so in 7a (similar to c4, which could precede senescence (Glück et al., 2017)). Similarly, cell cycle markers were lower in 7a than 7b. Thus, 7a was consistent with EBV^+^ cells that arrest almost immediately via innate sensing and become senescent, whereas 7b matched a trajectory in which EBV^+^ hyperproliferative cells become senescent following replicative stress response induction. Notably, both 7a and 7b exhibited elevated levels of ribosomal subunit mRNAs (**Figure S32C-D**).

NF-κB activated EBV^+^ cells (c2) exhibited loss of accessibility at 1,142 sites present within resting cells in both donors (8.7% of all resting cell peaks). This reduction paralleled upregulated expression of the polycomb group repressor *EZH2* and a polycomb-interacting methyltransferase, *DNMT1* (**Figure S33**). However, EBV-activated cells possessed 668 peaks absent in resting cells (*c2!c38*) that were linked to 595 unique genes. 154 of these (25.9%) were DEGs between resting and activated cells (Figure 6E). These 154 *c2!c38* DAP-linked DEGs included 109 upregulated and 45 downregulated genes from *c38àc2*. Upregulated genes included regulators of apoptosis and tumor suppression (*BCL2A1, TNFRSF8, PDCD1LG2, ST7, IQGAP2, TOPBP1, CD86*); proliferation (*CDCA7, MKI67*); B cell signaling (*NFKBIA, MAPK6, TNIP1, TRAF3*); inflammation (*SLC7A11, RXRA, ZC3H12C*); oxidative stress (*SLC15A4, TXN*); and epigenetic remodeling (*AHRR, NCOR2*). The 45 downregulated *c2!c38* DAP-linked DEGs included *CCR7*, acetyltransferases (*EPC1, KAT6B*), apoptotic and stress response regulators (*STK39, STK17A/DRAK1, VOPP1, ZDHHC14*), negative regulators of B cell signaling (*CBLB*), and the tumor suppressor *ARRDC3* (**Figure S34**).

In a third example, we explored DAP-linked DEGs between EBV-induced activated (c2) and differentiated (c5) phenotypes. Activated cells exhibited 999 called peaks that were absent in differentiated cells (*c2!c5* DAPs) while only 13 new peaks emerged in differentiated cells (*c5!c2* DAPs) in both donors. This corresponded to a 15% net reduction in accessible peaks in the *c2àc5* transition. Notably, *c2!c5* DAPs found in both donors were linked to 13.4% of all DEGs identified between these states in the scRNA assay. Key *c2àc5* DAP-linked DEG dynamics included downregulation of NF-κB family genes and upregulation of plasmablast-specific transcriptional regulators, translation factors, and protein export machinery (i.e., facilitators of Ig synthesis, secretion, and protein folding chaperones) (Figures 6F**, S10**).

By mapping multiome reads to a concatenated reference (human + EBV), we were able to detect increased accessibility within the EBV genome over time after infection. We detected 21 unique viral ATAC peaks (20 of 21 common to both donors) including at TSSs for essential viral genes such as the *EBNAs* and *LMP1* (**Figure S35**). Quantification of episome peak-containing cells by phenotype revealed that EBV^+^ activated and hyperproliferative cells had the greatest number of accessible loci relative to other post-infection phenotypes. These sites included the C promoter (Cp) for *EBNA1*, *EBNA2*, and *EBNA3A-C*; the *LMP1* TSS; the TSS for *BMRF1*, a DNA polymerase accessory protein; and the *BHLF1* locus, which was recently recognized as a facilitator of latency and B cell immortalization (Yetming et al., 2020) (**Figure S36A-B**). Innate arrested cells (c4) exhibited the lowest frequency of cells with accessible episomal loci, followed by growth-arrested cells (c7) (**Figure S36C**).

### Post-infection cell fates exhibit differential enrichment of TF motifs

To further investigate regulatory differences by phenotype, we assayed TF binding motif enrichment by state (**Figure S37**). We identified variable motif enrichment linked to resting cell phenotypes and among non-arrested post-infection states (**Figure S37A**). Variation in accessible motifs broadly aligned with phenotypic gene expression with respect to antiviral response induction, promotion of cell growth, and oncogenesis (**Figure S37B-C**). Activated B cell (c2) ATAC peaks were enriched in binding sites for proto-oncogenic TFs including members of the REL (cREL, RelA, RelB), AP-1 (FOS, FOSB, JUNB, JUND), and EGR (EGR1-4) families. Enhanced accessibility at NF-κB family binding sites within activated cells was noteworthy, given the observed concurrent upregulation of NF-κB pathway gene expression. Similar phenotypic consistency was observed within differentiated cells (c5), which were enriched in accessible motifs for IRF4, IRF8, and XBP1. Globally, both resting B cell phenotypes and the innate sensing arrest state shared the greatest motif correlation with each other (R>0.75) and the lowest correlation with EBV-activated and hyperproliferating cells (0.55<R<0.7) (**Figure S38**).

### An informatics pipeline to infer phenotype-resolved TF signatures and gene regulatory elements

The prevalence of DAPs linked to DEGs known to be modulated in *trans* by EBV gene products led us to interrogate phenotype-specific TF signatures genome-wide. To do so, we employed a bioinformatic workflow to obtain ChIP-seq referenced inferences of single-cell phenotypes from scATAC data, which we termed “crisp-ATAC” (**Figures S39, S40**). We expected that ensemble-averaged ChIP data from an appropriate reference cell type would contain TF binding (and epigenetic) data from a superposition of cell phenotypes at high coverage, thus maximizing chances to identify overlaps with comparatively sparse scATAC cluster data. We further reasoned that phenotypic variation in TF binding site accessibility would have biological consequences.

We sought to predict viral EBNA and LMP1-mediated NF-κB accessible sites at promoters, enhancers, and actively transcribed genes in each state of our model. To do so, we applied crisp-ATAC recipes to intersect peaks from each scATAC phenotype with ChIP-seq peaks for viral EBNAs, NF-κB/Rel TFs, H3K4me1, H3K4me3, H3K27ac, H3K36me3, and RNA Pol II (Jiang and Mortazavi, 2018) (**Figure S39C**). Hyperproliferative cells (c6), EBV-activated cells (c2), and resting memory B cells (c3) exhibited up to 3-fold more enhancers and promoters at known EBNA2 binding sites relative to naïve B cells (c8), other activation intermediates (c0, c1), plasmablasts (c5), arrested states (c4, c7), and non-B cells (c9, c10). Similar patterns were found for EBNA3C and EBNALP sites (**Figure S39C**, left column) as well as Rel family TF binding sites (cRel, RelA, and RelB). Enhancers, promoters, and actively transcribed genes were consistently enriched in c2, c3, and c6 and depleted in c4 and c7, with intermediate levels present in c0, c1, c5, and c8. By accounting for peaks conserved in both biological replicates, we demonstrated low-noise measurements of DAPs for use with crisp-ATAC and characterized DAP frequencies and interval length distributions across all pairwise phenotype comparisons (**Figure S40**).

### crisp-ATAC finds TF-linked expression signatures that vary across distinct EBV^+^ cell fates

We applied crisp-ATAC to capture regulatory variation among infection phenotypes with respect to key viral transcriptional co-activators. We first compared the innate sensing arrest (c4) and NF-κB (c2) states (Figure 7A), as these represent starkly different post-infection fates. Peak data were extracted and gated to obtain *c2!c4* DAPs present in both donors (n=1,873), which yielded linked gene predictions (n=1,514) (Figure 7B-C). The *c2!c4* linked gene ontology network was enriched for innate defense (inflammation, antimicrobial processes) and EBV-induced responses (lymphocyte activation, regulation of apoptosis) (Figure 7D). Taking a macroscopic view, we found that predicted *c2!c4* linked genes included 42.5% (71 of 167) of known EBV super-enhancer (EBVSE) site-linked genes (Zhou et al., 2015). Consequently, 41-55% of EBNA-associated *c2!c4* DAPs also overlapped a peak for the super-enhancer-associated host TF RelA. Of the 71 EBVSE-linked genes identified for *c2!c4*, 19 (27%) were linked to EBNALP ChIP-seq peaks; 22 (31%) to EBNA2 ChIP-seq peaks; and 15 (21%) to EBNA3C ChIP-seq peaks. EBVSE-linked genes were enriched in EBNA-associated DAP-linked DEGs relative to size-matched random samples of genes in the captured transcriptome (Figure 7E).

**Figure 7.**
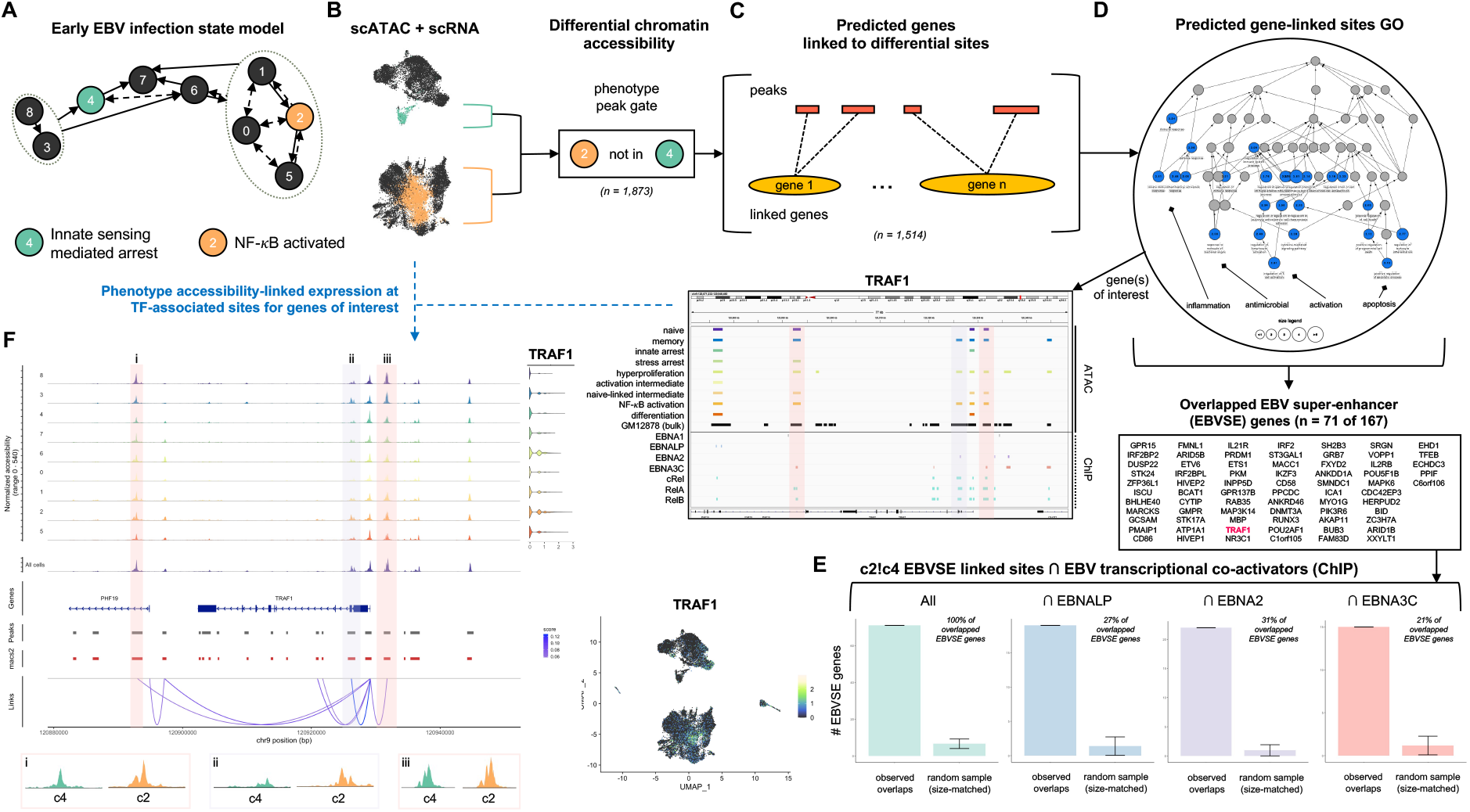
crisp-ATAC analysis of DAP-linked DEGs in activated versus innate arrested EBV^+^ B cells. (A) Schematic of NF-κB activation (c2) and innate arrest (c4) model phenotypes. (B) Multimodal assay gating to extract *c2!c4* DAPs. (C and D) Prediction of *cis*-regulatory linkage. All *c2!c4* peak intervals (n=1,873) are used as inputs to the GREAT (McLean et al., 2010) to predict *c2!c4* DAP-linked genes (n=1,514). (E) Occurrence of EBVSE-linked genes identified as *c2!c4* DAP-linked DEGs associated with select EBNA binding sites relative to expected frequency due to random overlap (n=100 simulation trials, error bars depict mean +/- standard deviation). Random samples were size-matched relative to each EBNA-associated gene list. (F) Example gene of interest analysis for *TRAF1*, an EBVSE-linked gene identified as a *c2!c4* DAP-linked DEG with phenotype-variable accessibility at multiple EBNA binding sites. *See also Figures S39-S42, S44-S47*

We analyzed specific genes of interest based on 1) EBVSE membership, 2) GO process involvement, and/or 3) empirically demonstrated importance to EBV infection. The NF-κB activated gene and signal transducer *TRAF1*, whose gene product interacts with viral LMP1, was identified through all three of these routes (Devergne et al., 1996; Eliopoulos et al., 2003; Fries et al., 1999; Greenfeld et al., 2015; Sandberg et al., 1997). We found *c2!c4* DAPs associated with one or more EBNA at −3kb, +2kb, and +37kb relative to the *TRAF1* TSS, each with significant positive correlation to *TRAF1* expression (p<0.05 for correlation z-score). Notably, these EBNA-associated regulatory loci exhibited reduced accessibility in stress arrest (c7), activation intermediate (c1) and differentiated (c5) states compared with c2 (Figure 7F).

We used a grouped crisp-ATAC comparison (*c256!c38*) to study changes associated with the trajectory for successful EBV-induced B cell immortalization (**Figure S41**). We analyzed viral co-activator-associated DAPs between proliferative (c6) and LCL-like phenotypes (c2, c5) versus resting B cells (c3, c8). Despite the net reduction in accessibility after infection, we identified 245 unique genes linked to 1,824 peaks present in all tested EBV^+^ states (c256) but absent in both resting phenotypes (c38) (**Figure S41A-C**). 166 of the 245 genes were linked to a binding site for at least one EBNA, and 18 of these genes overlapped with EBVSE targets (7.3% of predicted genes, 10.8% of known EBVSE genes). Only 31 GO process terms were shared across the top 100 terms for each tested EBNA, accounting for 15% of unique terms (**Figure S41D**). We selected the *c256!c38*∩EBNA targets *TNFRSF8* (CD30), *CD274* (PD-L1), and *PDCDL1G2* (PD-L2) based on their therapeutic relevance to EBV-associated lymphomas. For each gene, we confirmed the presence of EBNA-associated DAP-linked DEGs by phenotype. These included three *EBNA2_c256!c38_* sites near the *TNFRSF8* TSS (−17kb, −12kb, and +16kb) and a shared multi-EBNA site −17kb from the *PDCDL1G2* TSS and +43kb from the end of the *CD274* gene (**Figure S41E**). These loci were enriched for Rel sites and activating histone marks in LCL reference data.

In a final example, we evaluated activated versus differentiated EBV^+^ phenotypes (*c2!c5*) sites with known viral transcriptional co-activator binding in both donors to explore regulatory relationships that distinguished the phenotypes present in LCLs (SoRelle et al., 2021) (**Figure S42A**). 519 of 999 identified *c2!c5* peaks intersected with at least one EBNA binding site, from which 247 unique genes were predicted. 29 of 110 *c2!c5*∩EBNA2 site-linked genes (26.3%) were *c2!c5* DEGs in the scRNA assay, as were 34 of 115 *c2!c5*∩EBNALP site-linked genes (29.6%) and 20 of 125 *c2!c5*∩EBNA3C site-linked genes (16.0%). 20 site-linked genes were identified from all three viral co-activator recipes (**Figure S42B**), including the EBVSE-linked G protein coupled receptor *GPR137B*, a lysosmal transmembrane receptor that regulates mTORC1 activity and autophagy (Gan et al., 2019; Gao et al., 2012). *GPR137B* was also identified as a *c2!c38* DAP-linked DEG, indicating inaccessible regulatory loci within resting cells as well. We identified two regulatory DAPs with significant positive correlation to gene expression at +14kb and +18kb relative to the *GPR137B* TSS. One of these sites (+14kb) coincided with EBNALP and EBNA3C binding sites but did not exhibit a *c2!c5* DAP. However, the second site (+18kb) overlapped with all three intersected EBNAs and was a *c2!c5* DAP. Both sites also intersected with Rel family TFs (cRel, RelA, and RelB) (**Figure S42C**). Other genes involved in lysosome-mediated processes including autophagy and antigen presentation regulation were also identified from the *c2!c5* comparison (*TFEB*, *LAMP3*), albeit with modestly elevated but significant differential expression (Martina et al., 2012; Nagelkerke et al., 2014; Settembre et al., 2011). We also found numerous other genes involved in immune activation signaling, apoptosis, and transcriptional regulation that displayed EBNA-associated *c2!c5* DAP-linked DEGs (**Figure S42D-E**). Collectively, these vignettes of post-infection cell trajectories highlight the genome-wide dynamics of diverse EBV-induced B cell responses captured within the single-cell multiomics dataset.

### Single-cell imputation reveals transcriptomic dynamics of EBV infection of FCRL4^+^/TBX21^+^ B cells

The EBV early infection model presented herein captures robust B cell phenotypes, each of which are exhibited by >1000 cells in the assay. However, we did not preclude the possibility that distinct rare populations of B cells may also be represented within the dataset. To explore this possibility with sufficient sensitivity, we implemented a recently reported method that minimizes technical read noise from transcript dropout in scRNA data (Linderman et al., 2022). Briefly, this method (ALRA) adaptively thresholds a low-rank approximation of the single-cell expression matrix in order to preserve biological zeros (true negatives) for gene expression and impute probable values in place of technical zeros (false negatives from dropout). Unsupervised clustering of ALRA-imputed data from Day 5 of the early infection timecourse revealed a small (1.5% of cells) yet distinctive population of B cells (Figure 8A). Notably, this subset expressed *FCRL4*, which encodes an inhibitory receptor that blocks BCR signaling (Davis, 2007; Sohn et al., 2011); *TBX21 / T-bet*, a gene for a canonical Th1 subset homeobox TF (Szabo et al., 2000) that is also essential to an unconventional memory B cell subset (Johnson et al., 2020; Rubtsova et al., 2013; Wang et al., 2012); and *CXCR3*, which encodes a chemokine receptor that mediates migration in a subset of MBCs (Muehlinghaus et al., 2005). While these *FCRL4^+^ / TBX21^+^ / CXCR3^+^* cells displayed robust expression of lineage markers (e.g., *CD19*, *MS4A1/CD20*) and genes involved in the early stages of B cell activation (e.g., *CCR6*, *CD69*), they notably lacked expression of GC reaction gene signatures or plasmablast differentiation (Figure 8B). Rather, the most prominent feature of this population was broad upregulation of *FCRL* family genes including *FCRL5* and *FCRL6*, which was originally found to be expressed on NK and T cells (Wilson et al., 2007) but has recently been identified in B cell progenitors (Honjo et al., 2020). While these cells expressed markers of double-negative (DN & DN2) B cell populations (*TBX21, ITGAX, CXCR3*), such cells are typically FCRL4^-^ (Jenks et al., 2018; Scharer et al., 2019). This population was most consistent with Tissue-Like Memory (TLM) B cells based on the expression of the markers discussed above in addition to being *CD21^-^ / FCRL5^+^ / SOX5^+^ / RTN4R^+^,* although differential FCRL4 expression in TLM B cells has been reported (Li et al., 2016; Rakhmanov et al., 2009). Top markers of the *FCRL4^+^* TLM-like phenotype in our dataset were also enriched for genes with roles mediating innate immunity and inflammatory responses; proto-oncogenes including *FGR*, which has been shown to be induced by EBNA2 during EBV infection (Cheah et al., 1986; Knutson, 1990); and an array of genes whose overexpression contributes to cell migration, metastasis, plasticity, and the potential for self-renewal in a variety of cell types.

**Figure 8.**
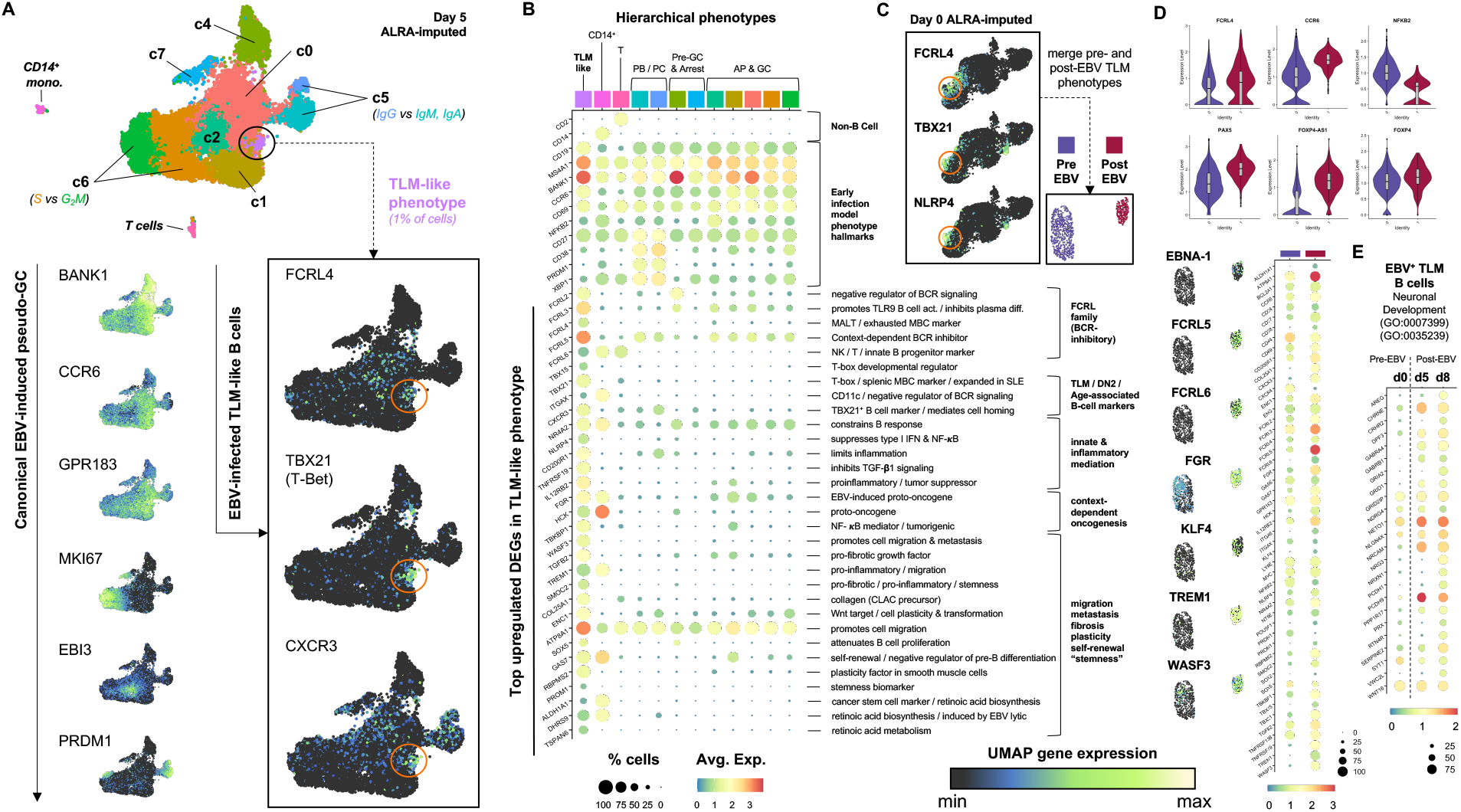
Discovery and characterization of EBV-driven transcriptomic dynamics in FCRL4^+^ / TBX21^+^ Tissue-Like Memory (TLM) B cells. (A) Unbiased clustering of Day 5 scRNA assay processed using biological zero-preserving imputation by adaptive thresholding of low-rank approximation (ALRA; see (Linderman et al., 2022)). Early infection model cluster numbering (top panel) and key markers of the EBV-induced pseudo-GC reaction are depicted (bottom-left column). A non-GC cluster of *FCRL4^+^ / TBX21^+^ / CXCR3^+^* B cells, consistent with TLM B cells, was also identified (bottom-right column). (B) Hierarchically ordered phenotypes from imputed Day 5 data based on expression representative model state genes and top markers of the TLM B phenotype. With respect to the depicted genes, the TLM B state shares greater similarity to non-B cell lineages than other infected B cell states. Annotations are provided for TLM B cell markers genes, which include previously identified lineage markers (e.g., *ITGAX/CD11c*). (C) Identification of the TLM B phenotype in ALRA-imputed Day 0 scRNA assay. Cells matching this phenotype from Day 0 and Day 5 were extracted and merged to evaluate differential gene expression pre- and post-EBV infection. (D) Differential gene expression in TLM B cells in response to EBV infection. (E) Expression of genes involved in nervous system development (GO:0007399, FDR = 0.0075) and tube morphogenesis (GO:0035239, FDR = 0.035) within TLM-like B cells before (d0) and after (d5, d8) EBV infection. *See also Figure S43, S44*

We confirmed the presence of a Day 0 (EBV^-^) precursor to the Day 5 TLM phenotype, and we focused on this Day 0 TLM B cell cluster to assess pre- and post-infection differential gene expression (Figure 8C). Although TLM B cells were *FCRL4^+^* at Day 0, the expression of *FCRL4* increased following infection, consistent with our prior finding that FCRL4 is a host biomarker of the EBV Latency IIb program (Messinger et al., 2019). *CCR6* expression in these cells increased following infection, indicating early B cell activation. However, key NF-κB pathway genes associated with subsequent GC B cell activation events were downregulated after infection in TLM B cells. Although upregulation of genes in this canonical pro-survival pathway was not observed, we found evidence for a *PAX5* / *FOXP4-AS1* / *FOXP4* axis, which has been reported to promote proliferation and anti-apoptotic signaling (Figure 8D**, violin plots**) (Wu et al., 2019). Critically, we observed EBV latent transcripts within TLM B cells at Day 5 and Day 8 in addition to upregulation of numerous genes with roles in cellular plasticity, tumorigenesis, migration, and self-renewal (Figure 8D**, dot plot**). In addition to *FCRL4*, *TBX21*, *CXCR3, ITGAX/CD11c, and FCRL5,* an unexpected set of genes known to mediate neural development, adhesion, and signaling were conserved and/or upregulated markers of this cell subset during the early infection timecourse (Figures 8E**, S43-S44**). These genes included *NRCAM, DPF3, PPP1R17, PCDH1, GABRA4,* and *GRID1*, which were identified through GO Process enrichment within TLM-like B cells (GO:0007399, FDR = 0.0075, 62 genes and GO:0035239, FDR = 0.035, 23 genes). Notably, several of these were also observable in subsets within the GM12878 LCL. Many of these and other genes upregulated within *FCRL4^+^* TLM-like B cells following EBV infection were also found in GM12878, which was derived from an independent biological donor (**Figure S43, S44**).

## Discussion

Our data reveal heterogeneity in coordinated gene expression and chromatin accessibility dynamics within individual cells during the critical early stages of a viral infection. By capturing the initial phases of EBV infection with high resolution -omics, we discern the gene regulatory environments associated with diverse infected cell fates and their respective developmental trajectories. These include genome-wide expression and chromatin signatures associated with effective host antiviral response, virus-triggered oncogene-induced senescence, and the path to sustained EBV latency and host cell immortalization, which is accessed via simulated B cell activation. Moreover, we develop a bioinformatic workflow to characterize post-infection outcomes through gene- and peak-level investigations at loci matching specific epigenetic patterns as well as host and viral transcription factor binding profiles. The combined high resolution multiomics data and integrative analytical framework reported herein yield a vividly detailed representation of the genome-wide interplay of host and virus.

We captured diverse post-infection B cell fate trajectories that, due to their asynchronous parallel emergence, cannot be resolved by ensemble sequencing methods. Remarkably, we find that large-scale euchromatin-to-heterochromatin transitions (20-40% reductions in genome-wide accessibility) can occur in post-infection trajectories and fates, including several that display increases in transcript levels. The scATAC data implicate Simpson’s paradox with respect to heterogeneous chromatin accessibility dynamics following infection, since the total number of unique peaks increases in early infected populations while the peaks per cell can, in fact, decrease (Simpson, 1951; Trapnell, 2015) (**Table S1**).

Global epigenetic silencing via SAHF formation (Di Micco et al., 2011) is most prominent both in cells that rapidly undergo innate sensing-mediated arrest and in cells that evade this response but arrest due to DNA damage response activation during virus-induced hyperproliferation. Our data also indicate a possible role for ribosome biogenesis in the transition from virus-induced arrest to senescence, likely through a p53-MDM2 axis (Deisenroth and Zhang, 2010), which critically regulates EBV transformation (Forte and Luftig, 2009). Intriguingly, the trajectory of successful infection is distinguished by increased accessibility at key sites against the larger backdrop of heterochromatin formation. A substantial number of these sites (many associated with viral TFs) have predicted *cis* linkages to genes enriched for regulation of apoptosis, tumor suppression, inflammation, and chromatin remodeling, all of which are determinants of successful EBV infection. Moreover, viral episome ATAC profile heterogeneity across infected phenotypes indicates that latency establishment depends on retained accessibility to viral genes within the repressive host milieu.

The identification of an EBV-infected analog of an AP-eMBC phenotype is consistent with results from *in vitro* and *in vivo* antigen stimulation experiments (Suan et al., 2017; Taylor et al., 2015), as well as previous work from our lab that identified CCR6 as an EBV Latency IIb program biomarker that becomes downregulated in the transition to Latency III in LCLs (Messinger et al., 2019).The development of this state *in vitro* implies that EBV may gain access to the memory pool *in vivo* via progenitors that have limited involvement in the GC reaction. In the context of normal antigen stimulation, this subset of eMBCs undergoes early exit from the cell cycle and GC reaction as a consequence of restricted access to or engagement with cognate antigen (Glaros et al., 2021). It is thus conceivable that EBV-infected B cells may differentially develop into GC BCs, PBs, or eMBCs from the AP state in a manner dependent on the extent to which the LMPs, EBNAs and other viral gene products induce mimicry of BCR activation and signaling. This model accommodates a surprising possibility – that EBV may access long-term persistence and survival not only within post-GC high-affinity MBCs but also via GC-independent eMBCs that avoid extensive proliferation.

The activated B cell and plasmablast phenotypes that developed within 5 days are generally consistent with our findings in LCLs (SoRelle et al., 2021) and resemble *in vivo* tonsil cell subset transcriptomes. Furthermore, differentiated plasmablasts exhibited fewer accessible sites in both host chromatin and viral episomes relative to activated cells. Notably, prior studies found that only 50% of EBV-infected cells that secrete immunoglobulin go on to become immortalized (Tosato et al., 1985). Additional work within LCLs demonstrated that EBV^+^ cells with upregulated Ig production exhibited reduced DNA synthesis and EBNA downregulation (Wendel-Hansen et al., 1987). Collectively, these findings support a model for continuous EBV-driven B cell differentiation *in vitro*, wherein plasmablasts are continually generated through activation-induced maturation yet selected against by their reduced proliferation. While this disadvantage limits viral replication via cell division, the reduced MYC levels, increased XBP1, and endoplasmic reticulum stress in these cells may support EBV lytic reactivation (Guo et al., 2020; Laichalk and Thorley-Lawson, 2005; Sun and Thorley-Lawson, 2007; Taylor et al., 2011).

Depending on the phenotype comparison, roughly 10-35% of all DEGs were correlated with DAPs. This range was similar to the frequencies of both differential accessibility of expressed TF binding motifs (23%) and DEGs associated with differentially accessible promoters (25%) identified in dexamethasone-treated A549 cells profiled with sci-CAR (Cao et al., 2018). The observation of similar DAP-linked DEG frequencies in response to diverse stimuli (e.g., chemically induced glucocorticoid receptor activation, viral infection) raises intriguing questions regarding the fundamental frequency of genes regulated, at least in part, by accessibility changes. The observed proportion implies that most DEGs may be regulated by higher order chromatin interactions and differential recruitment of transcriptional activators and/or RNA Pol II.

The crisp-ATAC method provides a simple, flexible informatic approach to map ChIP-seq profiles to cell phenotypes discovered from scATAC-seq. Thus, it provides a potential route for effectively bootstrapping scATAC resolution to datasets from other -omics modalities. When data from suitable reference cell types and/or states are available, crisp-ATAC can be used to predict phenotype-resolved gene regulation by evaluating simple or complex combinations of factor binding sites, histone modifications, and/or nuclear chromatin compartmentalization across all differentially accessible sites. In our case, cell-matched empirical scRNA-seq data can be cross-referenced with crisp-ATAC outputs for methodological validation and identification of genes of interest for future studies. This method is adaptable to comparisons of phenotype-resolved ATAC profiles in contexts such as other infections, development, and cell responses to stimuli or therapies. We expect this approach to be particularly powerful for exploring TF-associated regulatory changes in time-resolved studies of cellular behaviors.

The identification and characterization of genome-wide EBV infection dynamics within *FCRL4^+^* TLM B cells (as well as other B subsets including *FCRL4^+^ / TBX21^-^* and *FCRL4^-^ / TBX21^+^* B cells evident within our data) provides an intriguing road map for subsequent research into EBV pathogenesis in unique B cell niches and their potential role(s) in human disease. Specifically, the tissue-homing capacity, innate immune mediation roles, and progenitor-like features of TLM B cells make them a high-interest target cell type in a range of cancers and autoimmune conditions. Notably, TBX21 and CXCR3 have separately been found at high frequencies in specific B cell lymphoma subtypes including Chronic Lymphocytic Leukemia (CLL), splenic marginal zone lymphoma (SMZL), extranodal marginal zone lymphoma (EMZL), precursor B-cell Lymphoblastic Leukemia, and Hairy Cell Leukemia (Dorfman et al., 2004; Jones et al., 2000). Generally, it has been observed that expansion of B cells with a base expression profile of low CD21 (*CR2*) and inhibitory receptors including FCRL4 is a fundamental feature of chronic infections and (auto) immune disorders (Freudenhammer et al., 2020). Expanded atypical MBC populations expressing FCRL4 and ITGAX have been identified in patients with chronic *Plasmodium falciparum* infections living in regions with endemic malaria (Weiss et al., 2009) and in exhausted B cell populations (also CXCR3^+^) described in the context of HIV viremia (Moir et al., 2008). FCRL4^+^ / TBX21^+^ / ITGAX^+^ B cells have also recently been identified as a pathogenic subset in primary Sjögren’s Syndrome (pSS) with lymphomagenic potential (Verstappen et al., 2020). FCRL4^+^ / CD20^hi^ B cells expressing RANKL/TNFSF11 have been reported as a subset that contributes to inflammation in rheumatoid arthritis (RA) (Yeo et al., 2015) – this matches an additional phenotype that we observed in ALRA-imputed data from Day 8 and in the GM12878 LCL. Notably, RANKL/TNFSF11 was not expressed in these cells until after EBV infection (data not shown). A variety of B cells phenotypes have been found to be clonally expanded in systemic lupus erythematosus (SLE). These include CXCR3^+^/CD19^hi^ B cells (Nicholas et al., 2008), FCRL4^-^ DN2 cells (Jenks et al., 2018) and notably, a recent large cohort study identified IL-21 stimulation drove expansion of tissue-homing CD11c^+^ / T-bet^+^ (*ITGAX^+^ / TBX21^+^*) with significantly elevated levels of FCRL3, FCRL4, and FCLR5 (Wang et al., 2018). In the context of autoimmunity, comprehensive analysis of distinct B cell subsets – including how each of these may be affected by EBV infection – is especially relevant to resolving the etiology of multiple sclerosis (MS). While one report found clonally expanded B cells from the cerebrospinal fluid (CSF) of MS patients with upregulated TBX21, CXCR3, and SOX5, EBV reads were not identified in the samples (Ramesh et al., 2020). However, two recent reports that provide epidemiological (Bjornevik et al., 2022) and serological (Lanz et al., 2022) data supporting a causal role for EBV in at least some MS cases have ignited efforts to explore the mechanistic foundations of this causality. That effort, along with research into other diseases associated with EBV-induced B cell dysregulation, will benefit from understanding the nuances of viral pathogenesis in distinct B cell niches. The data developed and analyzed herein provide a comprehensive portrait of *de novo* EBV infection within the canonical GC model and offer a tantalizing initial glimpse into non-canonical infection dynamics in at least one atypical yet critical B cell phenotype. These non-canonical responses to EBV infection are exemplified by the apparent lineage-inappropriate expression of genes that mediate neural cell development, adhesion, and signaling within a lymphoid compartment. By combining high resolution, scale, and modalities, we hope this resource will facilitate advances in our understanding of the gene regulatory diversity intrinsic to peripheral B cell subsets and how that heterogeneity underlies previously obscured complexities of host-EBV dynamics.

### Limitations of the study

The reported single-cell multiomics data have several limitations. They do no capture aspects of host-virus dynamics acting at other molecular levels. Examples include epigenetic modifications (e.g., DNA methylation status), three-dimensional chromatin architecture changes, modulation of translation and protein abundance, post-translational modifications, protein-protein interactions, and signaling cascades (e.g., phosphorylation status). While we present a method for inferring DAP-linked TF binding and epigenetic modifications based on empirical scATAC data, we do not have direct ChIP-seq measurements at the single-cell level.

The reported bioinformatic methodology (crisp-ATAC) also has notable constraints. This approach is limited by the availability of ChIP-seq data from an appropriate cell type and target reference state. Moreover, regulatory site predictions must be empirically tested to validate potential functions in gene expression control. Finally, distance limits imposed for identifying *cis*-regulatory linkages preclude identification of distal gene regulatory elements formed via 3D nuclear conformation.

## Acknowledgments

The authors wish to thank members of the Duke University Molecular Genomics Core (MGC) and the Duke Center for Genomic and Computational Biology (GCB), especially Emily Hocke, Karen Abramson, Dr. Simon Gregory, and Dr. Nicolas Devos. Special thanks are also due to the anonymous donors, without whose blood donations this work would not have been possible. E.D.S. wishes to acknowledge funding support from the Department of Molecular Genetics and Microbiology Viral Oncology Training Grant (NIH T32 # 5T32CA009111-42). This work was supported by funding from the National Institute of Dental and Craniofacial Research (NIDCR award #R01DE025994-06).

## Experimental Methods

### PBMC isolation and B lymphocyte enrichment

Whole blood (50 mL each from two anonymous donors; TX1241/Donor 1 & TX1242/Donor 2) was obtained from the Gulf Coast Regional Blood Center (Houston, TX). Upon receipt, peripheral blood mononuclear cells (PBMCs) from each donor sample were isolated via Ficoll gradient separation (Histopaque®-1077, Sigma #H8889), resuspended at 10×10^6^ cells/mL in RPMI 1640 + 15% heat-inactivated fetal bovine serum (FBS, v/v, Corning) (R15 media), and incubated overnight at 37°C and 5% CO_2_. The next day, CD19^+^ B cells were enriched from donor PBMCs via negative isolation (BD iMag Negative Isolation Kit, BD Biosciences #558007) and resuspended at 2×10^6^ cells/mL in R15 supplemented with 2 mM L-glutamine, 100 units/mL penicillin, 100 µg/mL streptomycin, and 0.5 µg/mL cyclosporine A (R15^+^ media). Roughly 45×10^6^ B cells were recovered per donor post-enrichment. Following CD19^+^ validation (see below: Flow cytometry), enriched B cell aliquots (1-2 mL at 3×10^6^ cells/mL) were viably frozen in 90% FBS + 10% DMSO and stored in liquid N_2_.

### EBV infection and cell culture

The type 1 EBV strain B95-8 was obtained in-house as viral supernatant from the inducible B95-9 Z-HT cell line as reported previously (Johannsen et al., 2004). Immediately after withholding and cryopreserving uninfected enriched B cells for each donor (day 0 samples), the remaining cells in culture were infected with B95-8 via resuspension in viral supernatant (100 µL per 1×10^6^ cells) for 1 h at 37°C and 5% CO_2_. Infected B cells from each donor were rinsed with 1x PBS, resuspended in R15^+^ media, and incubated at the conditions described above throughout the course of infection. Aliquots were taken from each infected donor culture at 2-, 5-, and 8-days post-infection and viably frozen as described for uninfected day 0 samples.

### Flow cytometry

The extent of B cell enrichment was quantified for each donor using flow cytometry. Following negative isolation, cell samples (2×10^5^ per donor) were rinsed with FACS buffer (1x PBS + 2% FBS), stained with phycoerythrin (PE)-conjugated mouse anti-human CD19 (BioLegend, clone HIB19; catalog #302208; lot #B273508) in the dark for 30 min at room temperature, then rinsed again prior to analysis. Cell samples at each timepoint were prepared as described and co-stained with PE-anti-CD19 and allophycocyanin-conjugated mouse anti-human CD23 (APC-anti-CD23, BioLegend, clone EBVCS-5; catalog #338513; lot #B273489) to validate successful EBV infection. To validate the AP-eMBC phenotype (c0), enriched resting B cells from two additional donors (TX1253 and TX1254) were labeled with CellTrace Violet (ThermoFisher / Invitrogen, Cat #34571) and stained with one of the following combinations at days 0, 2, 5, and 8: CCR6/Memory panel (FITC-anti-CD27, PE-anti-CCR6, and APC-anti-CD23); Naïve/Memory panel (FITC-anti-IgD, PE-anti-CD19, and APC-anti-CD27); or CCR6/Naïve panel (FITC-anti-IgD, PE-anti-CCR6, and APC-anti-CD23). FITC-anti-CD27, FITC-anti-IgD, and APC-anti-CD27 were purchased from BioLegend (Cat #302806, #348206, and #356410, respectively) and PE-anti-CCR6 was purchased from Invitrogen (Cat #12-1969-42). Compensation matrices were calculated from single-stain controls for FITC and PE and applied to all samples for analysis. All cytometry measurements were acquired with a BD FACSCanto II (BD Biosciences) and analyzed using FlowJo version 10 (Ashland / BD Biosciences).

### Human tonsil sample preparation

Tonsillar B cells were isolated from discarded, anonymized tonsillectomies from the Duke Biospecimen Repository and Processing Core (BRPC; Durham, NC). Tonsil tissue samples were manually disaggregated, filtered through a cell strainer, and isolated by layering over a cushion made from Histopaque-1077 (H8889; Sigma-Aldrich). Harvested lymphocytes were washed three times with FACS buffer (5% FBS in PBS) prior to scRNA library preparation.

### Preparation of scRNA and scATAC libraries

Cryopreserved samples from each early infection timepoint of interest were simultaneously thawed by donor and purified to > 90% viable cells by Ficoll gradient separation. Viable cells from each timepoint and donor were then prepared as single-cell matched gene expression (scRNA) and chromatin accessibility (scATAC) libraries by the Duke Molecular Genomics Core staff with the 10x Chromium Next GEM Single Cell Multiome ATAC + Gene Expression Kit (10x Genomics, Pleasanton, CA) (Satpathy et al., 2019; Zheng et al., 2017). Briefly, nuclei were isolated from each sample and subjected to transposition at accessible chromatin sites. Next, transposed nuclei, barcoding master mix, and gel beads containing unique barcode sequences were prepared into single-cell GEMs (Gel bead emulsions) using the Chromium Controller and Chip J. Within each GEM, poly-adenylated (poly-A) mRNA transcripts from individual nuclei are captured by barcoded, indexed poly-T primers and reverse transcribed into cDNA. Simultaneously, a separate barcoded sequence containing a spacer and Illumina P5 adaptor sequence is added to transposed regions within the captured nucleus. Barcoded multiomes were then purified, pooled, and pre-amplified by PCR prior to library construction. The scATAC library for each sample is generated by PCR amplification and incorporation of sample index and Illumina P7 adaptor sequences. Separately, pre-amplified gene expression cDNA is further PCR amplified, fragmented, and size selected. The scRNA library for each sample is then constructed using PCR to incorporate the P5 and P7 sequencing adaptors. Two biological replicates of tonsillar lymphocytes were prepared as scRNA libraries using the 10x Genomics Next GEM 3’ Gene Expression kit with v3 chemistry (10x Genomics, Pleasanton, CA), and sequenced, processed, aligned, and analyzed as described above for early infection scRNA samples.

### Sequencing, alignment, and count matrix generation

The 8 paired-end scATAC libraries (4 timepoints per 2 donors) were pooled onto two lanes of an Illumina S2 flow cell and sequenced at a target depth of 25,000 reads per cell on an Illumina NovaSeq (Illumina, San Diego, CA). The 8 paired-end scRNA libraries were similarly pooled and sequenced at a target depth of 50,000 reads per cell. Tonsil scRNA libraries were likewise pooled and sequenced at 50,000 reads per cell. All sequencing runs were performed by staff at the Duke Center for Genomic and Computational Biology. Raw base calls for each assay were prepared as sample-demultiplexed FASTQ files using *cellranger-arc mkfastq* (Cellranger, 10x Genomics), a wrapper of the Illumina *bcl2fastq* function. Next, sample-matched scRNA and scATAC reads were aligned against genomic references to produce multiome count matrices using *cellranger-arc count*. One set of count matrices was generated by mapping reads to a concatenated genomic reference constructed from the human genome (GRCh38) with the ∼172 kB type 1 EBV genome (NC_007605) included as an extra chromosome. These outputs were used for downstream RNA-only analyses to capture host and viral gene expression. A second set of count matrices generated by mapping to GRCh38 only was used for downstream joint RNA and ATAC analyses. Compatible reference packages were assembled from the relevant genome (.fa) and annotation (.gtf) files using *cellranger-arc mkref*.

### Data QC and scMultiome analysis

All direct analysis of scRNA and scATAC data was conducted in R using Signac (Stuart et al., 2021), an extension of Seurat (Macosko et al., 2015; Satija et al., 2015; Stuart et al., 2019). Following read mapping and counting, linked scRNA and scATAC data were obtained from between 8,934-20,000 cells per sample. After QC filtering by mitochondrial expression (n < 20%), total transcripts (n < 25,000), unique transcripts (n > 1,000), and minimum cells expressing a given feature (n > 3), data from between 8,376-19,310 cells per sample were analyzed (see **Table S1**). The mitochondrial gene expression threshold was selected based on the high metabolic activity of early-infected B cells and the high cell viability observed in each sample (> 90%) immediately prior to library preparation to preserve biologically relevant phenotypes (Osorio and Cai, 2021). After QC filtering, a total of 52,271 and 44,920 cells were analyzed across the infection timecourse for donors TX1241 and TX1242, respectively. For RNA-only analysis, gene expression data (host and viral) across all timepoints for a given donor were merged into a single object, log normalized, scored for cell cycle markers, and scaled with cell cycle scoring regressed out. The top 2,000 differentially expressed features over the early infection timecourse data were identified and used for principal component analysis (PCA). The top 30 principal components were further dimensionally reduced via uniform manifold approximation projection (UMAP, (McInnes et al., 2018)), and clustering was performed to identify biologically distinct cell subpopulations. Merged scRNA dataset pseudotime trajectories were calculated using Monocle3 (Qiu et al., 2017), and were mapped along with cluster identities to 3D UMAP coordinates for visualization (Qadir, 2019; Qadir et al., 2020). For joint ATAC and RNA analysis, host gene expression and chromatin accessibility were analyzed for each separate timepoint and for a merged object containing Day 0 and Day 8 multiome data. Nucleosome signal and transcription start site (TSS) enrichment were calculated and used for QC filtering (Nucleosome.signal < 2, TSS.enrichment > 1). ATAC peaks were called using macs2 (Liu, 2014) with hg38 annotations. Gene expression data in the joint analysis was processed as described for the RNA-only analysis with the exception of using Signac’s SCTransform function instead of log normalization for expression counts. Top differential features in each assay (‘peaks’ and ‘SCT’) were determined, and multimodal neighboring and UMAP were performed for integrated data visualization. Cluster identities defined in the RNA-only assay were mapped to this merged joint dataset, which contained cells representing all identified subpopulations. Peaks with significant (anti-) correlation (p < 0.05 for z-score of correlation coefficients) to differentially expressed genes were identified using the LinkPeaks function in Signac, which was informed by SHARE-seq (Ma et al., 2020).

### crispATAC workflow and reference data curation

ChIP-referenced inference from single-cell phenotype ATAC (crispATAC) was developed to predict subpopulation-resolved gene regulatory features. In a typical workflow, cluster-level chromatin accessibility tracks are cross-referenced against ChIP-Seq (Chromatin Immunoprecipitation Sequencing) profiles for epigenetic marks and TFs of interest measured from a reference cell phenotype (in this study, lymphoblastoid cell lines such as GM12878). In this study, cluster-specific called peaks from the joint scATAC + scRNA dataset were extracted and prepared as simplified genomic range files (3-column .bed file format). Next, the desired ChIP-Seq datasets for the reference phenotype were downloaded and, where applicable, converted to .bedgraph format to be used as input for peak calling with the *macs2* function *bdgpeakcall*. The ChIP (and Hi-C) datasets used for crispATAC in this study (**Table S2**) are all publicly available from the National Center for Biotechnology Information (NCBI) Gene Expression Omnibus (GEO). Once all ATAC-seq and ChIP-seq peak files were generated, all were used as inputs to a single call of the bedtools (Quinlan and Hall, 2010) function *multiinter*, which output a matrix of all genomic range intersection intervals where at least one input file exhibited a peak. This intersection matrix was imported to R as a data frame and analyzed to identify common and/or differential intervals (matrix rows) among scATAC cluster phenotypes, epigenetic marks, and TFs using Boolean logic gating by dataset (matrix columns). For a given crispATAC recipe (e.g., peaks in scATAC cluster 1 not in scATAC cluster 2 intersected with EBNA2 ChIP peaks = [c1 ∩ !c2] ∩ EBNA2), the genome intervals matching the gating criteria were returned and converted to .bed files. Lists of differentially accessible, transcription-factor associated sites generated in this way were subsequently analyzed with the Genomic Regions Enrichment of Annotations Tool (GREAT) (McLean et al., 2010) to identify potential *cis*-regulated genes within 1 megabase of each query site. As a final step, output lists of potential linked genes were intersected with the top marker genes identified from the corresponding cluster-wise comparison in the scRNA assay, thus integrating direct single-cell RNA and ATAC measurements with subpopulation-resolved regulatory inferences from ensemble ChIP profiles. In a similar but separate approach, scATAC and ChIP peaks were intersected with topologically associated domain (TAD) boundaries (prepared using hicExplorer, (Ramírez et al., 2018; Wolff et al., 2018; Wolff et al., 2020)) and nuclear subcompartments from GM12878 Hi-C data to study differentially accessible TF-associated sites in the context of 3D nuclear architecture.

### Visualization of crispATAC outputs, gene ontologies, and networks

Data for genes of interest identified from crispATAC recipes were explored using dimensionally reduced (UMAP) expression maps and cluster-level accessibility tracks (Signac, (Stuart et al., 2021)), called peaks aligned with TFs and epigenetic marks (IGV, (Robinson et al., 2011)), and local neighborhoods in Hi-C contact maps (Juicebox, (Durand et al., 2016)). Cluster-resolved gene ontologies were generated and quantified by GREAT (McLean et al., 2010). Top scRNA assay cluster markers and GREAT output gene lists were also visualized as annotated networks using Cytoscape (Shannon et al., 2003).

## Notes

### Competing Interest Statement

The authors have declared no competing interest.

### Summary of Updates

We have identified a unique sub-population of human B cells within our scRNA-seq dataset akin to tissue-like memory (TLM) B cells using a recently reported method for zero-preserving imputation (ALRA) reported by Lindermann et al (Nat Comm, 2022). Unexpectedly, we find evidence for upregulated expression of gene sets within this sub-population associated with stemness/plasticity, migration, and neuronal cell development, adhesion, and signaling following EBV infection.

