## Supplementary Figures for "Fate-resolved gene regulatory signatures of individual B lymphocytes in the early stages of Epstein-Barr Virus infection"

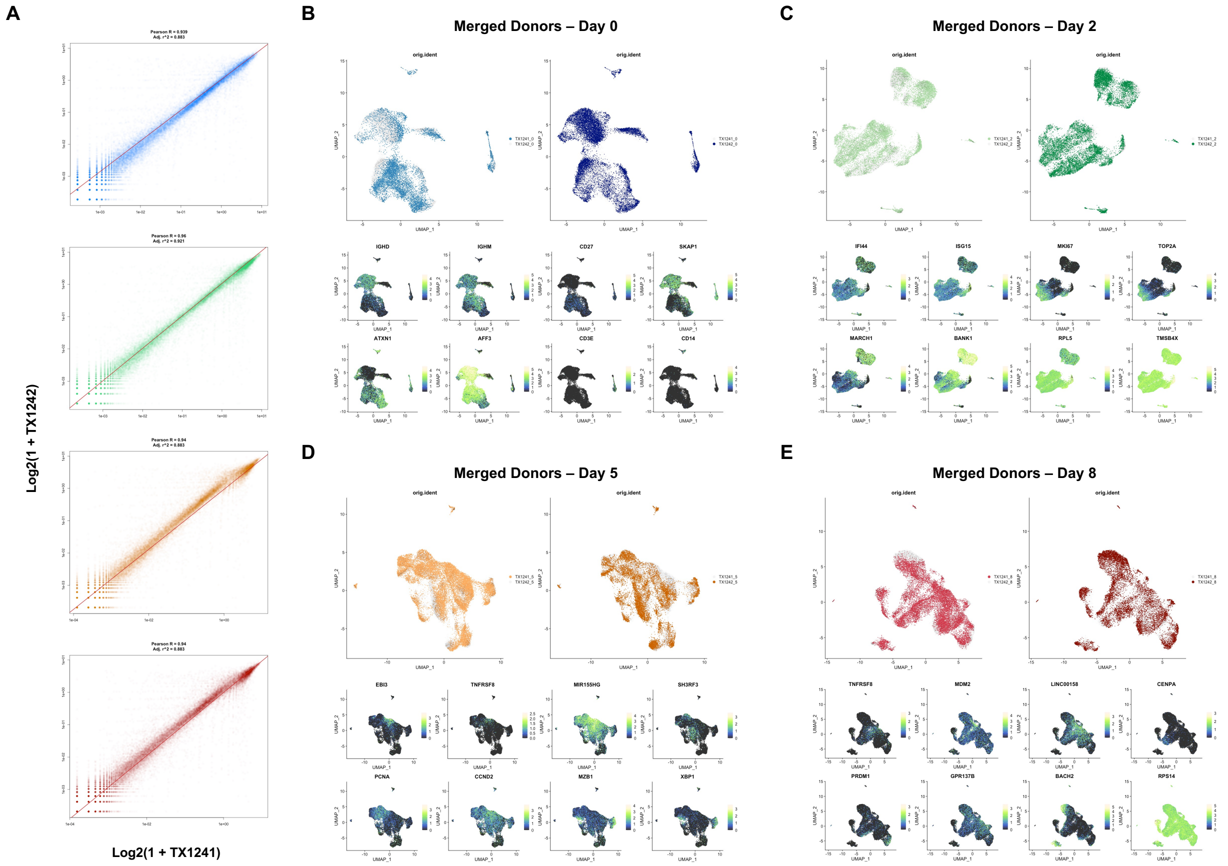


**Figure S1. Concordance of single-cell early infection data across two independent donors (TX1241 and TX1242)**

(A) Genome-wide expression correlation by day between donors.

(B) Donor-merged data from Day 0 samples. Cells from TX1241 (light blue) and TX1242 (dark blue) exhibit similar colocalization in dimensionally reduced projections and similar patterns of differentially expressed genes. In these examples, data from both donors are presented without cell cycle marker regression.

(C) Donor-merged data from Day 2 for TX1241 (light green) vs TX1242 (dark green), as in (A).

(D) Donor-merged data from Day 5 for TX1241 (light orange) vs TX1242 (dark orange), as in (A and B).

(E) Donor-merged data from Day 8 for TX1241 (light red) vs TX1242 (dark red), as in (A, B, and C).

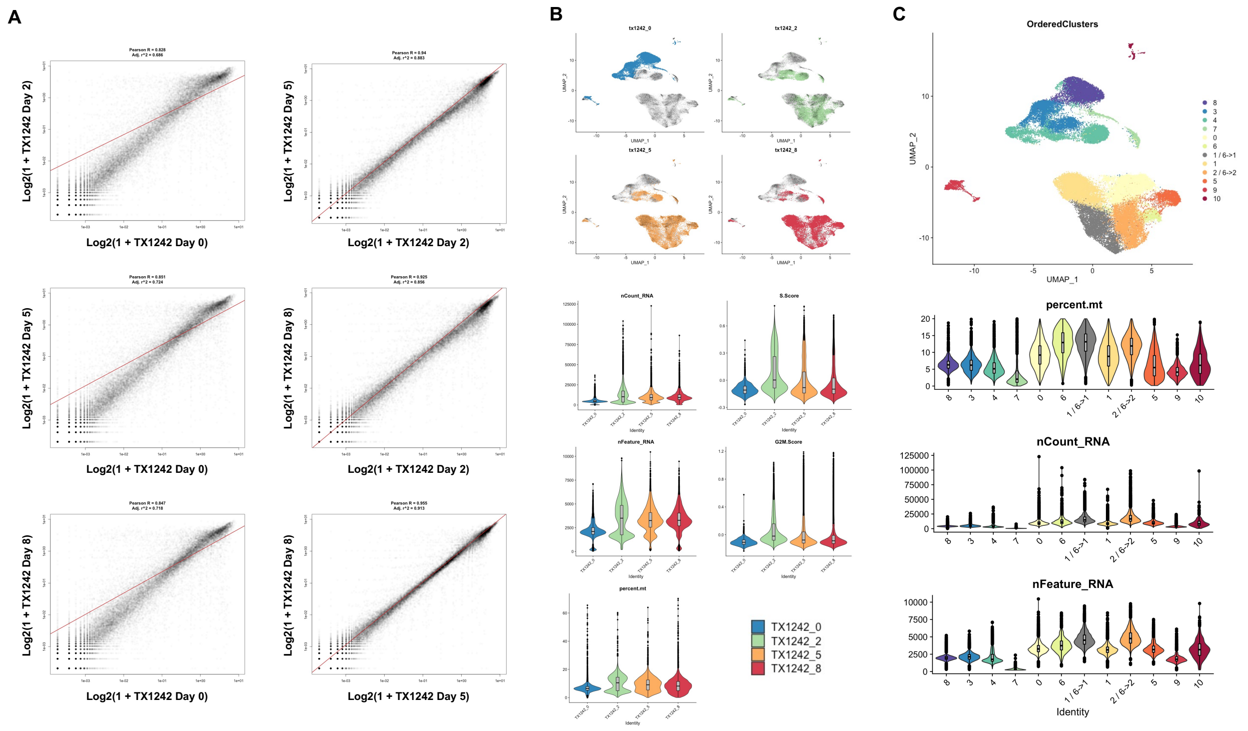


**Figure S2. Merged scRNA timecourse for a second donor (TX1242)**

(A) All pairwise timepoint genome-wide correlations for TX1242. The highest inter-day correlation is observed for Day 5 vs Day 8.

(B) TX1242 scRNA dimensionally reduced data and QC summary by sample timepoint.

(C) TX1242 cell cluster identification and general trends by phenotype. Data are presented without cell cycle marker regression, leading to the observation of two transitional phenotypes.

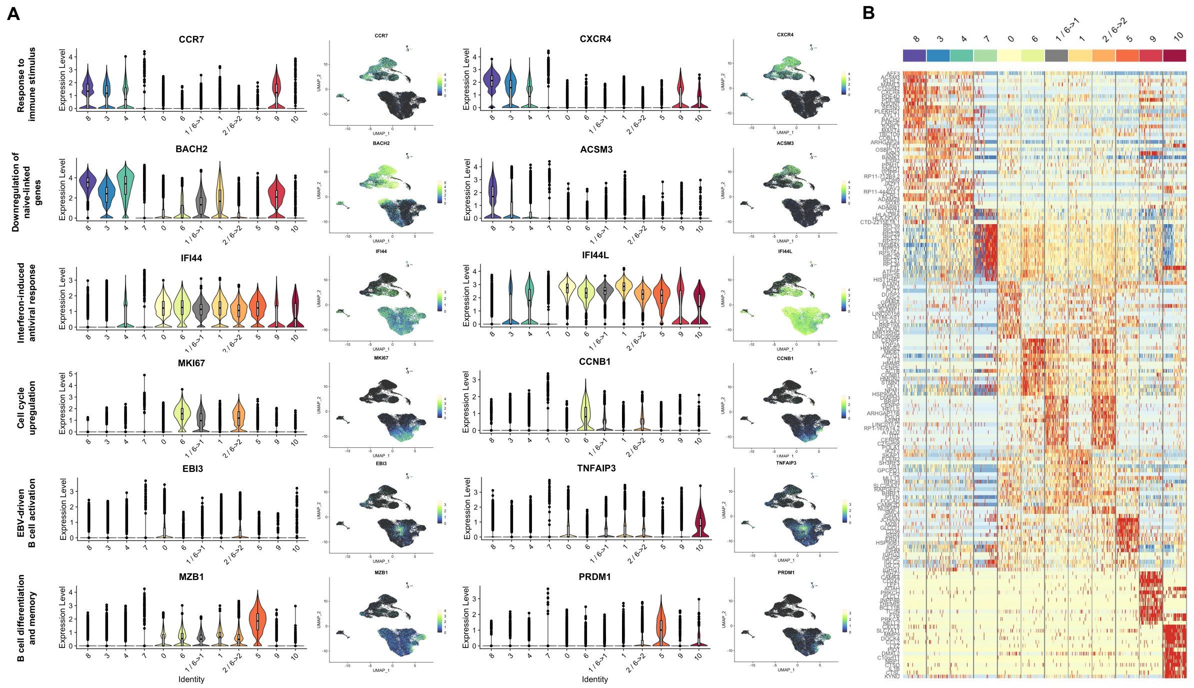


**Figure S3. Top marker gene analysis in TX1242**

(A) Phenotype-resolved violin plots and UMAP visualization of select marker genes by response class in TX1242.

(B) Expression of the top 15 markers per cluster in 100 randomly sampled cells per cluster (as in Fig. 1F for TX1241).

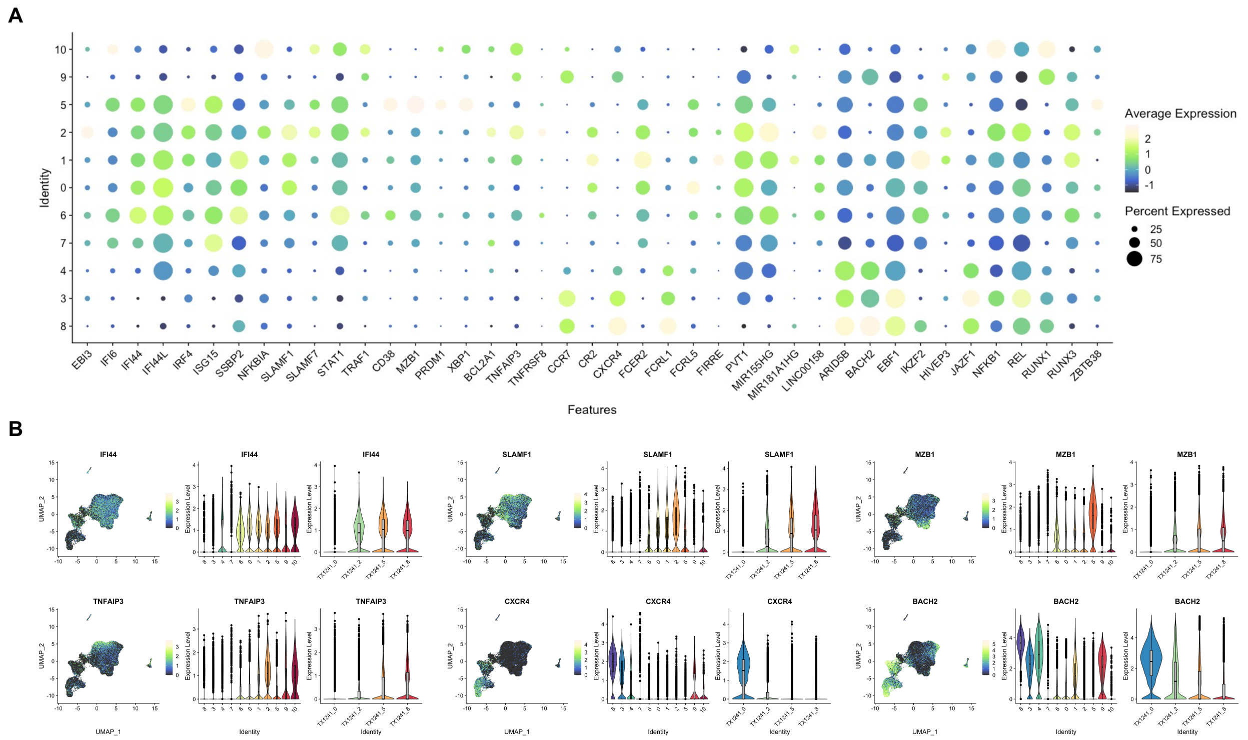


**Figure S4. Cluster- and time-resolved profiles for select DEGs of interest.**
(A) Average and cluster fractional expression of select DEGs.
(B) UMAPs and violin plots (by cluster and timepoint) of expression for select DEGs in (A).

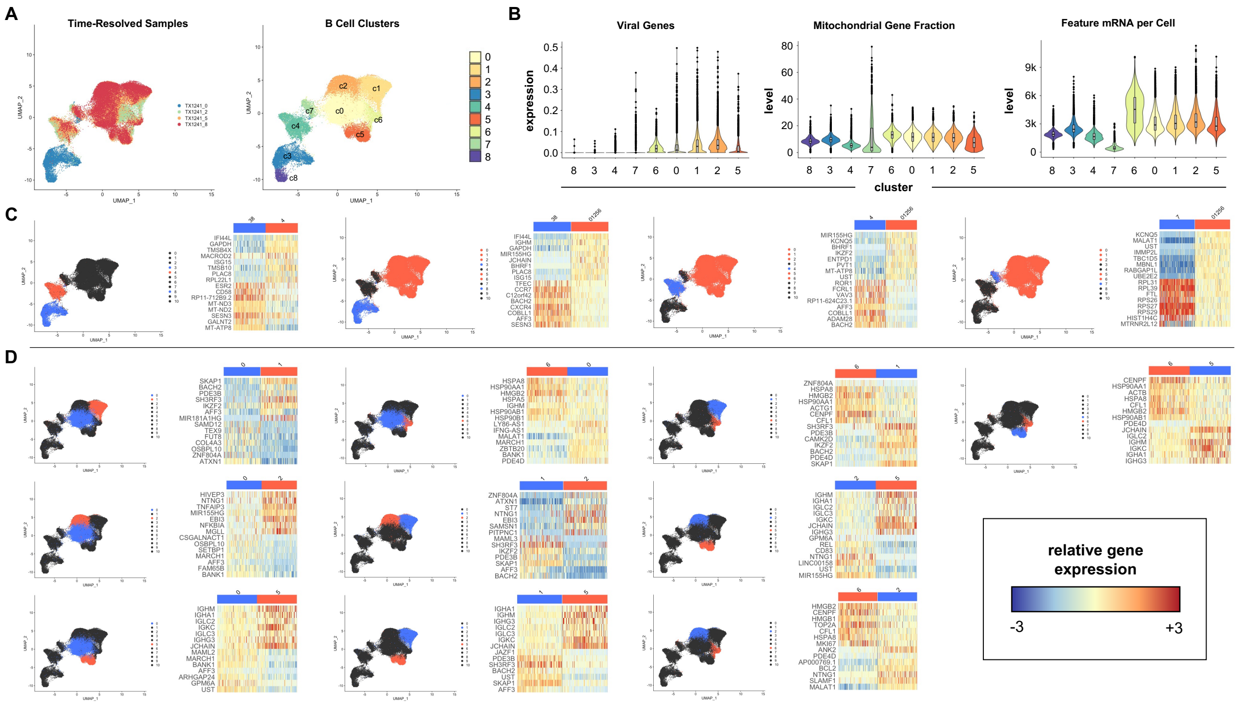

**Figure S5. High-resolution dissection of infected B cell phenotypes**
(A and B) Overview of global gene expression trends by phenotype. Fractions of viral transcripts were captured via alignment to the hg38 reference genome with type 1 EBV genome (NC_007605) concatenated as an extra chromosome.
(C) Pairwise differential gene expression between selected cluster groups. Clusters 3 and 8 are uninfected cells (day 0). Clusters 4 and 7 are two distinct EBV$^{+}$ phenotypes with variable hallmarks of arrest. Clusters 0,1,2,5,and 6 are distinct EBV$^{+}$ phenotypes with variable hallmarks of progressive infection (i.e., non-arrested).
(D) All pairwise combinations of differential gene expression for non-arrested EBV$^{+}$ phenotypes (clusters 0,1,2,5, and 6).

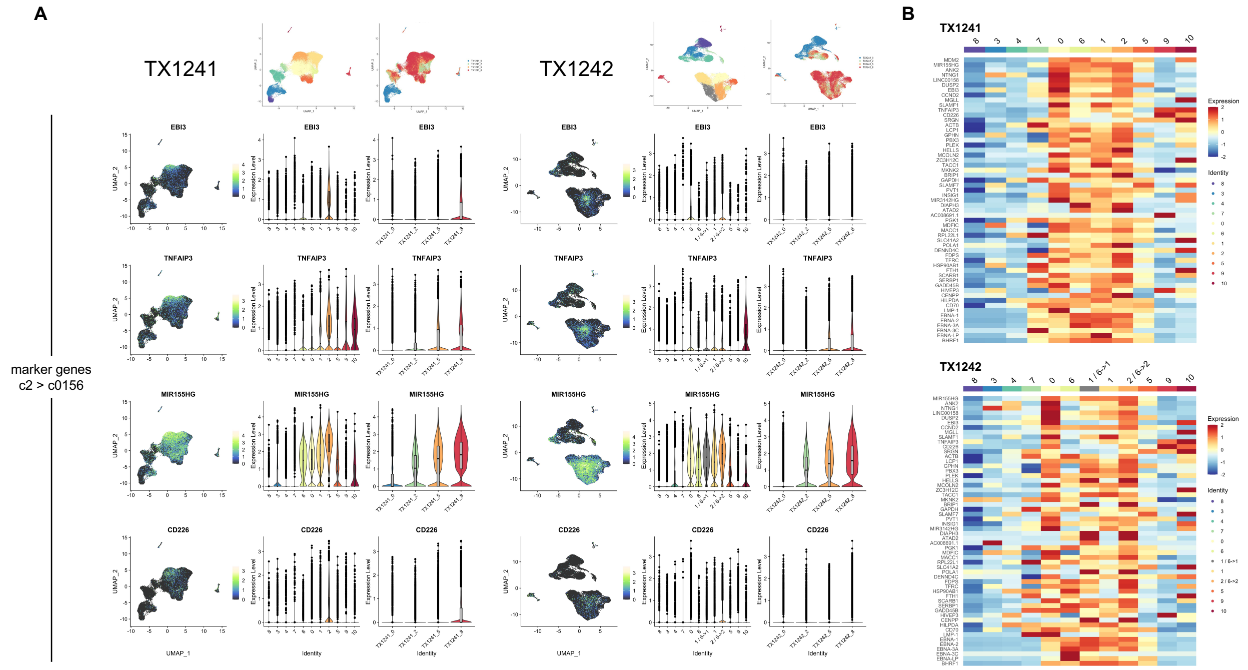


**Figure S6. Cluster 2 expression across both donors (EBV+ NF-κB activated state).**
(A) UMAP and violin plots for select c2*Δc0156* DEGs in TX1241 and TX1242.
(B) Phenotype average expression of the top 40 c2*Δc0156* DEGs (and viral genes of interest) in TX1241 and TX1242.

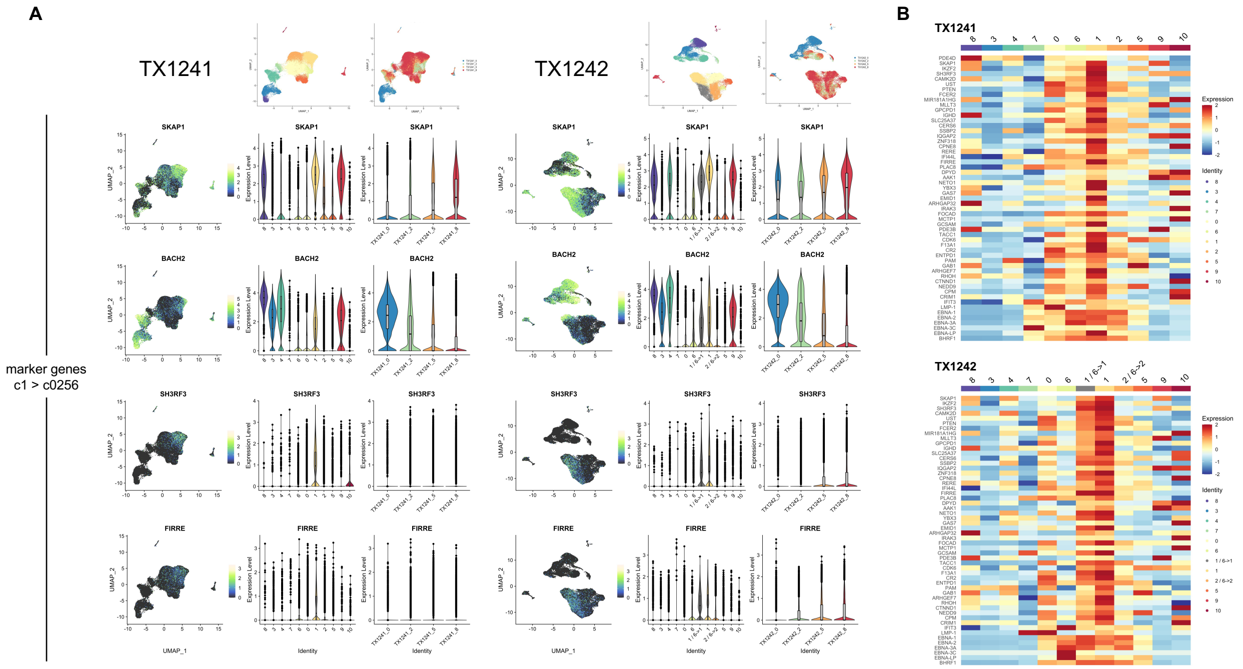


**Figure S7. Cluster 1 expression across both donors (EBV$^{+}$ naïve-linked activation intermediate).**
(A) UMAP and violin plots for select c1*Δc0256* DEGs in TX1241 and TX1242.
(B) Phenotype average expression of the top 40 c1*Δc0256* DEGs (and viral genes of interest) in TX1241 and TX1242.

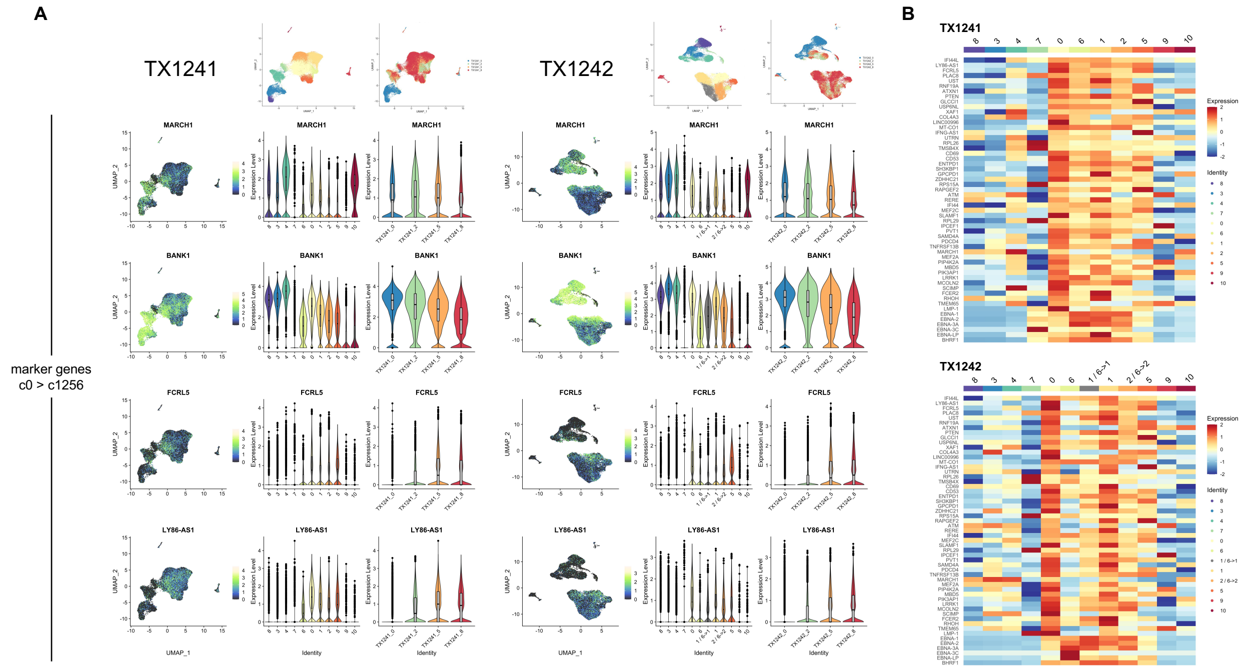


**Figure S8. Cluster 0 expression across both donors ($EBV^+$ activation intermediate).**
(A) UMAP and violin plots for select c0*Δc1256* DEGs in TX1241 and TX1242.
(B) Phenotype average expression of the top 40 c0*Δc1256* DEGs (and viral genes of interest) in TX1241 and TX1242.

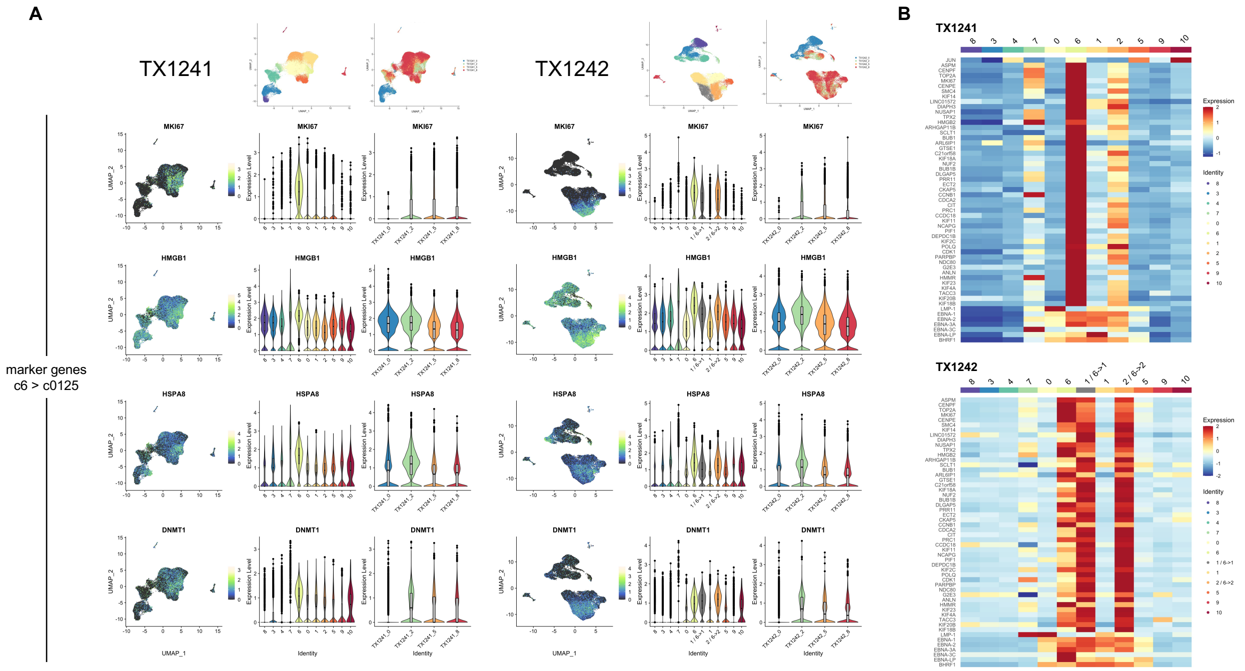


**Figure S9. Cluster 6 expression across both donors (EBV$^+$ hyperproliferation state).**
(A) UMAP and violin plots for select c6*Δc0125* DEGs in TX1241 and TX1242.
(B) Phenotype average expression of the top 40 c6*Δc0125* DEGs (and viral genes of interest) in TX1241 and TX1242.

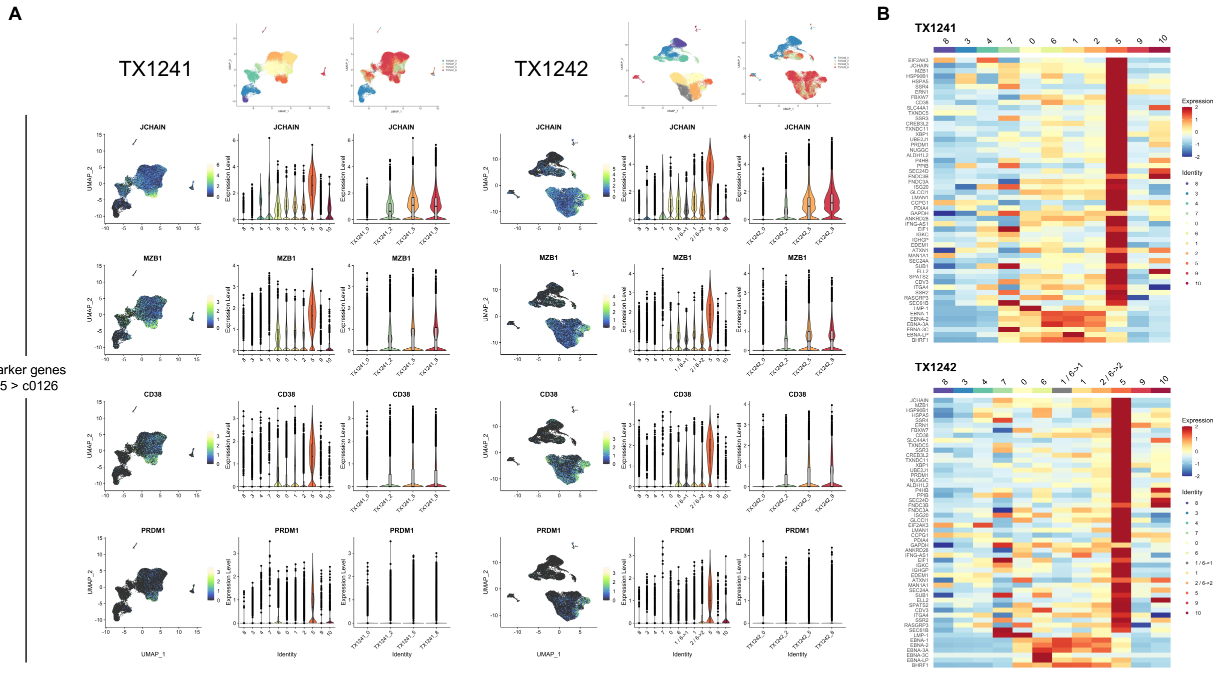


**Figure S10. Cluster 5 expression across both donors (EBV$^+$ differentiated plasmablast state).**

(A) UMAP and violin plots for select c5*Δc0126* DEGs in TX1241 and TX1242.

(B) Phenotype average expression of the top 40 c5*Δc0126* DEGs (and viral genes of interest) in TX1241 and TX1242.

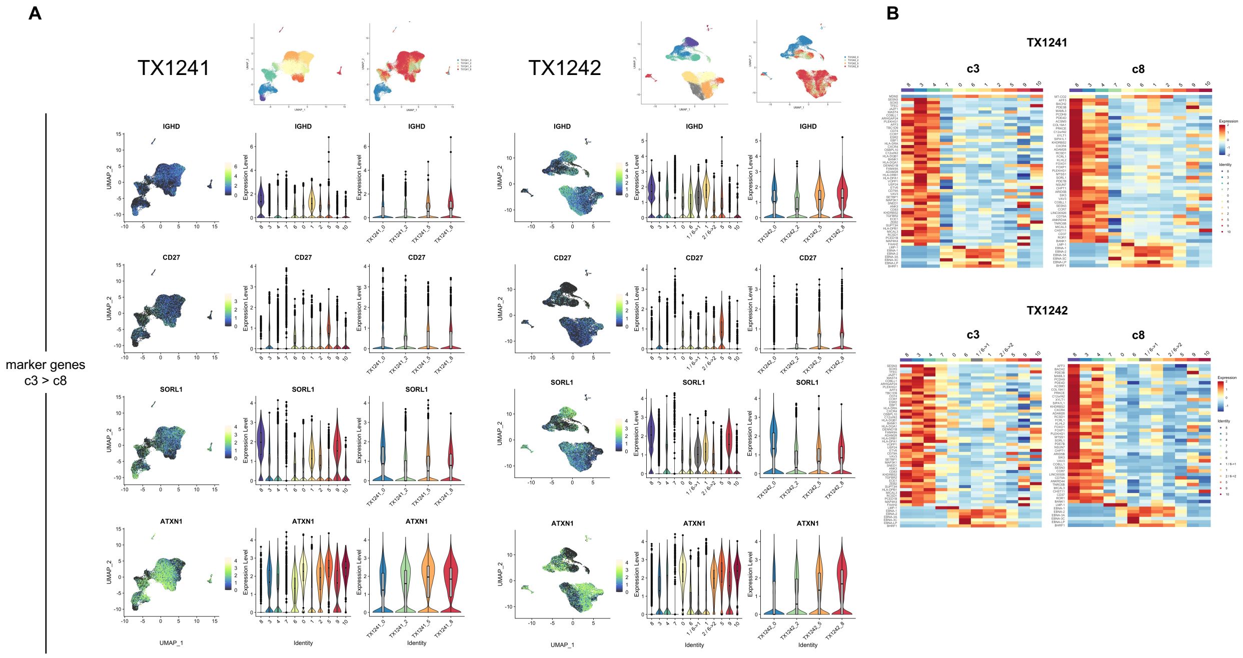


**Figure S11. Cluster 3 and 8 expression across both donors (Uninfected memory and naïve B cells).**
(A) UMAP and violin plots for select c3*Δc8*DEGs in TX1241 and TX1242.
(B) Phenotype average expression of the top 40 c3*Δc8* DEGs (and viral genes of interest) in TX1241 and TX1242.

**Cluster 8: expression markers**

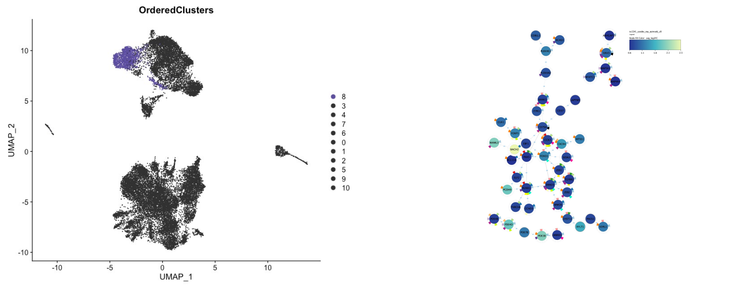


| GO ID | GO process | FDR | # genes |
|---|---|---|---|
| GO.0048583 | regulation of response to stimulus | 2.46E-8 | 30 |
| GO.2000113 | negative regulation of cellular macromolecule biosynthetic process | 8.18E-6 | 16 |
| GO.0002682 | regulation of immune system process | 1.04E-5 | 16 |
| GO.0048704 | embryonic skeletal system morphogenesis | 1.8E-4 | 5 |
| GO.0023057 | negative regulation of signaling | 2.1E-4 | 13 |
| GO.0051241 | negative regulation of multicellular organismal process | 2.6E-4 | 12 |
| GO.0006198 | cAMP catabolic process | 2.6E-4 | 3 |
| GO.0050854 | regulation of antigen receptor-mediated signaling pathway | 4.2E-4 | 4 |
| GO.0051347 | positive regulation of transferase activity | 4.6E-4 | 9 |
| GO.0030595 | leukocyte chemotaxis | 5.2E-4 | 5 |
| GO.0051240 | positive regulation of multicellular organismal process | 9.4E-4 | 13 |
| GO.0048731 | system development | 9.8E-4 | 22 |
| GO.0002768 | immune response-regulating cell surface receptor signaling pathway | 9.8E-4 | 6 |
| GO.0035239 | tube morphogenesis | 0.0016 | 8 |
| GO.0051223 | regulation of protein transport | 0.0017 | 8 |
| GO.0030155 | regulation of cell adhesion | 0.0017 | 8 |
| GO.1903243 | negative regulation of cardiac muscle hypertrophy in response to stress | 0.0025 | 2 |
| GO.0048584 | positive regulation of response to stimulus | 0.0025 | 14 |
| GO.0046649 | lymphocyte activation | 0.0028 | 6 |
| GO.0001954 | positive regulation of cell-matrix adhesion | 0.0031 | 3 |
| GO.0046626 | regulation of insulin receptor signaling pathway | 0.0039 | 3 |
| GO.1901699 | cellular response to nitrogen compound | 0.0043 | 7 |
| GO.0051249 | regulation of lymphocyte activation | 0.0043 | 6 |
| GO.0002062 | chondrocyte differentiation | 0.0052 | 3 |
| GO.0001817 | regulation of cytokine production | 0.0059 | 7 |
| GO.0044249 | cellular biosynthetic process | 0.0061 | 21 |
| GO.0030111 | regulation of Wnt signaling pathway | 0.0068 | 5 |
| GO.0070482 | response to oxygen levels | 0.009 | 5 |
| GO.1901888 | regulation of cell junction assembly | 0.0102 | 3 |
| GO.0071731 | response to nitric oxide | 0.0104 | 2 |
| GO.0032460 | negative regulation of protein oligomerization | 0.0104 | 2 |
| GO.0002407 | dendritic cell chemotaxis | 0.0104 | 2 |
| GO.0048589 | developmental growth | 0.0107 | 5 |
| GO.0031346 | positive regulation of cell projection organization | 0.0109 | 5 |
| GO.0051896 | regulation of protein kinase B signaling | 0.0119 | 4 |
| GO.0030099 | myeloid cell differentiation | 0.0119 | 4 |
| GO.0050767 | regulation of neurogenesis | 0.0123 | 7 |
| GO.0034067 | protein localization to Golgi apparatus | 0.0125 | 2 |
| GO.0022603 | regulation of anatomical structure morphogenesis | 0.0129 | 8 |
| GO.0030206 | chondroitin sulfate biosynthetic process | 0.0146 | 2 |
| GO.0006366 | transcription by RNA polymerase II | 0.0162 | 7 |
| GO.0051491 | positive regulation of filopodium assembly | 0.0175 | 2 |
| GO.0051282 | regulation of sequestering of calcium ion | 0.0181 | 3 |
| GO.0080134 | regulation of response to stress | 0.0188 | 9 |
| GO.0051150 | regulation of smooth muscle cell differentiation | 0.02 | 2 |
| GO.0043552 | positive regulation of phosphatidylinositol 3-kinase activity | 0.0209 | 2 |
| GO.0050853 | B cell receptor signaling pathway | 0.0219 | 2 |

**Cluster 3: expression markers**

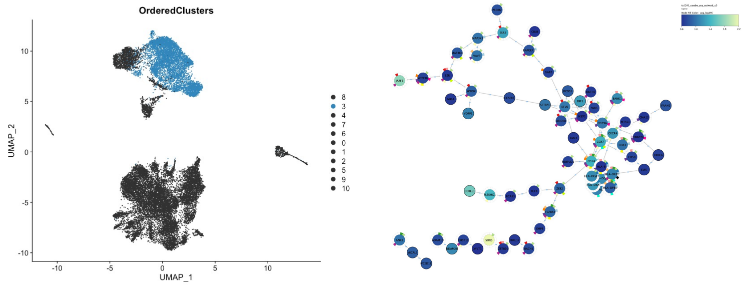


| GO ID | GO process | FDR | # genes |
|---|---|---|---|
| GO.0002684 | positive regulation of immune system process | 2.45E-6 | 17 |
| GO.0060333 | interferon-gamma-mediated signaling pathway | 2.97E-6 | 7 |
| GO.0048518 | positive regulation of biological process | 3.77E-6 | 39 |
| GO.0022407 | regulation of cell-cell adhesion | 3.05E-5 | 10 |
| GO.0001817 | regulation of cytokine production | 2.4E-4 | 11 |
| GO.0043410 | positive regulation of MAPK cascade | 2.8E-4 | 10 |
| GO.0000122 | negative regulation of transcription by RNA polymerase II | 0.0018 | 11 |
| GO.0071823 | protein-carbohydrate complex subunit organization | 0.0019 | 2 |
| GO.0050670 | regulation of lymphocyte proliferation | 0.0019 | 6 |
| GO.0002521 | leukocyte differentiation | 0.0021 | 7 |
| GO.0044093 | positive regulation of molecular function | 0.0023 | 16 |
| GO.0002381 | immunoglobulin production involved in immunoglobulin mediated immune response | 0.0025 | 3 |
| GO.0001775 | cell activation | 0.0025 | 12 |
| GO.0048523 | negative regulation of cellular process | 0.0028 | 28 |
| GO.0009966 | regulation of signal transduction | 0.0029 | 22 |
| GO.1902105 | regulation of leukocyte differentiation | 0.0041 | 6 |
| GO.0006366 | transcription by RNA polymerase II | 0.0041 | 10 |
| GO.0090087 | regulation of peptide transport | 0.0047 | 9 |
| GO.0070372 | regulation of ERK1 and ERK2 cascade | 0.0049 | 6 |
| GO.0032649 | regulation of interferon-gamma production | 0.0054 | 4 |
| GO.0002606 | positive regulation of dendritic cell antigen processing and presentation | 0.0059 | 2 |
| GO.0009653 | anatomical structure morphogenesis | 0.0074 | 16 |
| GO.0048701 | embryonic cranial skeleton morphogenesis | 0.0079 | 3 |
| GO.0010628 | positive regulation of gene expression | 0.0082 | 15 |
| GO.0032872 | regulation of stress-activated MAPK cascade | 0.0084 | 5 |
| GO.0002792 | negative regulation of peptide secretion | 0.0087 | 4 |
| GO.0046598 | positive regulation of viral entry into host cell | 0.0089 | 2 |
| GO.0002664 | regulation of T cell tolerance induction | 0.0101 | 2 |
| GO.0050854 | regulation of antigen receptor-mediated signaling pathway | 0.0104 | 3 |
| GO.0032147 | activation of protein kinase activity | 0.0109 | 6 |
| GO.0019883 | antigen processing and presentation of endogenous antigen | 0.0117 | 2 |
| GO.0038095 | Fc-epsilon receptor signaling pathway | 0.0138 | 3 |
| GO.0042113 | B cell activation | 0.014 | 4 |
| GO.0016045 | detection of bacterium | 0.014 | 2 |
| GO.0007009 | plasma membrane organization | 0.0145 | 3 |
| GO.0120036 | plasma membrane bounded cell projection organization | 0.0163 | 10 |
| GO.0002407 | dendritic cell chemotaxis | 0.0186 | 2 |
| GO.0045597 | positive regulation of cell differentiation | 0.0211 | 9 |
| GO.0032388 | positive regulation of intracellular transport | 0.0216 | 4 |
| GO.0050900 | leukocyte migration | 0.0225 | 5 |

**Figure S12. GO process networks for uninfected resting naïve (c8) and memory (c3) B cells.**

**Cluster 4: expression markers**

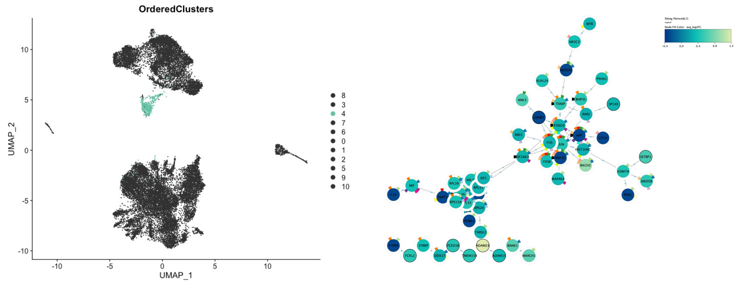


| GO ID | GO process | FDR | # genes |
|---|---|---|---|
| GO.0006413 | translational initiation | 3.52E-14 | 14 |
| GO.0010629 | negative regulation of gene expression | 1.03E-12 | 28 |
| GO.0044267 | cellular protein metabolic process | 7.26E-9 | 33 |
| GO.0010038 | response to metal ion | 2.59E-5 | 9 |
| GO.0051591 | response to cAMP | 3.9E-4 | 5 |
| GO.0051412 | response to corticosterone | 8.4E-4 | 3 |
| GO.0009605 | response to external stimulus | 8.7E-4 | 18 |
| GO.0014070 | response to organic cyclic compound | 0.0012 | 11 |
| GO.2000112 | regulation of cellular macromolecule biosynthetic process | 0.0023 | 25 |
| GO.0006366 | transcription by RNA polymerase II | 0.0023 | 10 |
| GO.0007568 | aging | 0.0027 | 6 |
| GO.0000122 | negative regulation of transcription by RNA polymerase II | 0.0028 | 10 |
| GO.1900449 | regulation of glutamate receptor signaling pathway | 0.0056 | 3 |
| GO.0043065 | positive regulation of apoptotic process | 0.0074 | 8 |
| GO.2000649 | regulation of sodium ion transmembrane transporter activity | 0.0076 | 3 |
| GO.0009612 | response to mechanical stimulus | 0.0076 | 5 |
| GO.0006417 | regulation of translation | 0.0076 | 6 |
| GO.0034599 | cellular response to oxidative stress | 0.0091 | 5 |
| GO.0010739 | positive regulation of protein kinase A signaling | 0.0102 | 2 |
| GO.0045637 | regulation of myeloid cell differentiation | 0.0133 | 5 |
| GO.0035458 | cellular response to interferon-beta | 0.0158 | 2 |
| GO.1903209 | positive regulation of oxidative stress-induced cell death | 0.0176 | 2 |
| GO.0051090 | regulation of DNA-binding transcription factor activity | 0.0176 | 6 |
| GO.0048468 | cell development | 0.0176 | 12 |
| GO.0032880 | regulation of protein localization | 0.0176 | 9 |
| GO.0000380 | alternative mRNA splicing, via spliceosome | 0.0176 | 2 |
| GO.1902165 | regulation of intrinsic apoptotic signaling pathway in response to DNA damage by p53 class mediator | 0.0185 | 2 |
| GO.0035994 | response to muscle stretch | 0.0197 | 2 |
| GO.0031103 | axon regeneration | 0.0197 | 2 |
| GO.0030224 | monocyte differentiation | 0.0197 | 2 |
| GO.2001014 | regulation of skeletal muscle cell differentiation | 0.0208 | 2 |
| GO.0045947 | negative regulation of translational initiation | 0.0208 | 2 |
| GO.0000185 | activation of MAPKKK activity | 0.0208 | 2 |
| GO.0001774 | microglial cell activation | 0.0224 | 2 |
| GO.0010628 | positive regulation of gene expression | 0.0256 | 13 |
| GO.0080135 | regulation of cellular response to stress | 0.0257 | 7 |
| GO.0014888 | striated muscle adaptation | 0.0307 | 2 |
| GO.0006952 | defense response | 0.0314 | 10 |
| GO.0033032 | regulation of myeloid cell apoptotic process | 0.0333 | 2 |
| GO.0034660 | ncRNA metabolic process | 0.0337 | 6 |

**Cluster 7: expression markers**

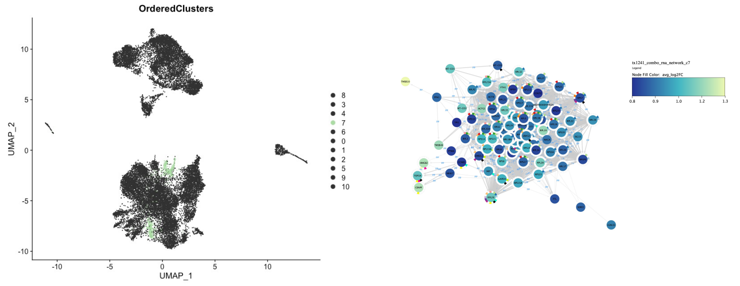


| GO ID | GO process | FDR | # genes |
|---|---|---|---|
| GO.0006614 | SRP-dependent cotranslational protein targeting to membrane | 1.76E-135 | 74 |
| GO.0002181 | cytoplasmic translation | 7.03E-55 | 34 |
| GO.0042254 | ribosome biogenesis | 8.65E-37 | 35 |
| GO.0042255 | ribosome assembly | 1.3E-22 | 17 |
| GO.0042273 | ribosomal large subunit biogenesis | 5.99E-20 | 16 |
| GO.0042274 | ribosomal small subunit biogenesis | 1.02E-16 | 14 |
| GO.0000462 | maturation of SSU-rRNA from tricistronic rRNA transcript (SSU-rRNA, 5.8S rRNA, LSU-rRNA) | 1.19E-6 | 6 |
| GO.0010608 | posttranscriptional regulation of gene expression | 8.77E-6 | 13 |
| GO.1904667 | negative regulation of ubiquitin protein ligase activity | 1.03E-5 | 4 |
| GO.0019646 | aerobic electron transport chain | 4.74E-5 | 4 |
| GO.0000470 | maturation of LSU-rRNA | 1.9E-4 | 4 |
| GO.1902229 | regulation of intrinsic apoptotic signaling pathway in response to DNA damage | 9.3E-4 | 4 |
| GO.0034101 | erythrocyte homeostasis | 0.0013 | 5 |
| GO.0000461 | endonucleolytic cleavage to generate mature 3'-end of SSU-rRNA from (SSU-rRNA, 5.8S rRNA, LSU-rRNA) | 0.0022 | 2 |
| GO.1901798 | positive regulation of signal transduction by p53 class mediator | 0.0034 | 3 |
| GO.2001244 | positive regulation of intrinsic apoptotic signaling pathway | 0.0035 | 4 |
| GO.0006880 | intracellular sequestering of iron ion | 0.0071 | 2 |
| GO.0034250 | positive regulation of cellular amide metabolic process | 0.0085 | 5 |
| GO.1903580 | positive regulation of ATP metabolic process | 0.0104 | 3 |
| GO.0031145 | anaphase-promoting complex-dependent catabolic process | 0.012 | 3 |
| GO.0043902 | positive regulation of multi-organism process | 0.013 | 8 |
| GO.0032075 | positive regulation of nuclease activity | 0.014 | 2 |
| GO.0007093 | mitotic cell cycle checkpoint | 0.0158 | 5 |
| GO.0051338 | regulation of transferase activity | 0.0164 | 13 |
| GO.0006265 | DNA topological change | 0.0167 | 2 |
| GO.0071897 | DNA biosynthetic process | 0.0194 | 4 |
| GO.0071824 | protein-DNA complex subunit organization | 0.0236 | 6 |
| GO.0010942 | positive regulation of cell death | 0.0251 | 10 |
| GO.0061844 | antimicrobial humoral immune response mediated by antimicrobial peptide | 0.0277 | 4 |
| GO.0071156 | regulation of cell cycle arrest | 0.0308 | 4 |
| GO.0071428 | rRNA-containing ribonucleoprotein complex export from nucleus | 0.0318 | 2 |
| GO.0017148 | negative regulation of translation | 0.0357 | 4 |
| GO.0090305 | nucleic acid phosphodiester bond hydrolysis | 0.0394 | 6 |

**Figure S13. GO process networks for EBV$^{+}$ arrested cell states (c4 and c7).**

Cluster 0: expression markers

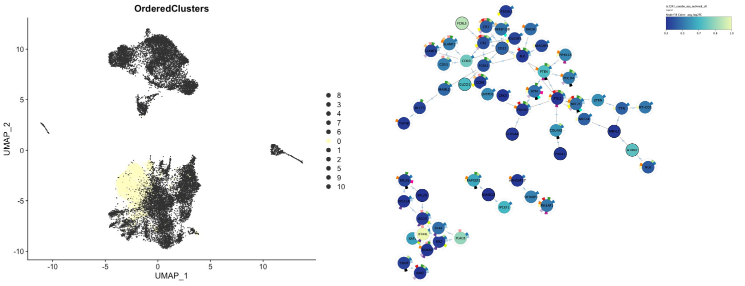


| | GO ID | GO process | FDR | # genes |
|---|---|---|---|---|
| | GO.0009607 | response to biotic stimulus | 1.62E-6 | 19 |
| | GO.0050896 | response to stimulus | 2.64E-6 | 45 |
| | GO.0007166 | cell surface receptor signaling pathway | 9.1E-5 | 21 |
| | GO.0048584 | positive regulation of response to stimulus | 1.3E-4 | 20 |
| | GO.0002252 | immune effector process | 0.0013 | 12 |
| | GO.0002253 | activation of immune response | 0.0018 | 8 |
| | GO.0002250 | adaptive immune response | 0.0018 | 7 |
| | GO.1904706 | negative regulation of vascular smooth muscle cell proliferation | 0.0026 | 3 |
| | GO.1902531 | regulation of intracellular signal transduction | 0.0026 | 16 |
| | GO.0016310 | phosphorylation | 0.0035 | 13 |
| | GO.0006959 | humoral immune response | 0.0061 | 6 |
| | GO.0031347 | regulation of defense response | 0.0082 | 9 |
| | GO.0009615 | response to virus | 0.0082 | 6 |
| | GO.0014066 | regulation of phosphatidylinositol 3-kinase signaling | 0.0111 | 4 |
| | GO.0035051 | cardiocyte differentiation | 0.0124 | 4 |
| | GO.0048468 | cell development | 0.014 | 13 |
| | GO.0002521 | leukocyte differentiation | 0.0142 | 6 |
| | GO.0043065 | positive regulation of apoptotic process | 0.0149 | 8 |
| | GO.0000165 | MAPK cascade | 0.015 | 6 |
| | GO.0048513 | animal organ development | 0.016 | 19 |
| | GO.0010038 | response to metal ion | 0.0166 | 6 |
| | GO.0019221 | cytokine-mediated signaling pathway | 0.0193 | 8 |
| | GO.0002923 | regulation of humoral immune response mediated by circulating imm... | 0.0193 | 2 |
| | GO.0002430 | complement receptor mediated signaling pathway | 0.0193 | 2 |
| | GO.0030888 | regulation of B cell proliferation | 0.0228 | 3 |
| | GO.0007568 | aging | 0.0232 | 5 |
| | GO.0007167 | enzyme linked receptor protein signaling pathway | 0.0232 | 8 |
| | GO.0060337 | type I interferon signaling pathway | 0.0243 | 3 |
| | GO.0034655 | nucleobase-containing compound catabolic process | 0.0244 | 6 |
| | GO.0050774 | negative regulation of dendrite morphogenesis | 0.0254 | 2 |
| | GO.0043517 | positive regulation of DNA damage response, signal transduction by p... | 0.0254 | 2 |
| | GO.0051241 | negative regulation of multicellular organismal process | 0.0265 | 10 |

Cluster 6: expression markers

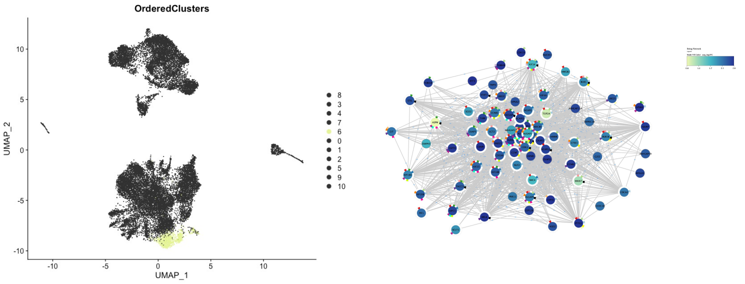


| | GO ID | GO process | FDR | # genes |
|---|---|---|---|---|
| | GO.0051301 | cell division | 4.61E-54 | 53 |
| | GO.0140014 | mitotic nuclear division | 2.7E-30 | 26 |
| | GO.0007346 | regulation of mitotic cell cycle | 3.53E-24 | 34 |
| | GO.0000226 | microtubule cytoskeleton organization | 1.91E-23 | 29 |
| | GO.0051983 | regulation of chromosome segregation | 4E-22 | 19 |
| | GO.0051276 | chromosome organization | 7.6E-22 | 38 |
| | GO.0000910 | cytokinesis | 3E-16 | 14 |
| | GO.0090068 | positive regulation of cell cycle process | 1.11E-14 | 19 |
| | GO.1902749 | regulation of cell cycle G2/M phase transition | 2.82E-14 | 16 |
| | GO.0044772 | mitotic cell cycle phase transition | 5.07E-14 | 18 |
| | GO.0051640 | organelle localization | 4.41E-13 | 23 |
| | GO.0051225 | spindle assembly | 7.41E-12 | 11 |
| | GO.0051303 | establishment of chromosome localization | 9.6E-12 | 11 |
| | GO.0051302 | regulation of cell division | 2.15E-11 | 14 |
| | GO.0031145 | anaphase-promoting complex-dependent catabolic process | 2.33E-11 | 9 |
| | GO.0010948 | negative regulation of cell cycle process | 2.69E-11 | 16 |
| | GO.0006323 | DNA packaging | 5.94E-11 | 14 |
| | GO.1903046 | meiotic cell cycle process | 9.03E-11 | 13 |
| | GO.0045841 | negative regulation of mitotic metaphase/anaphase transition | 1.87E-9 | 7 |
| | GO.0070507 | regulation of microtubule cytoskeleton organization | 3.18E-9 | 12 |
| | GO.0008283 | cell population proliferation | 2.12E-8 | 19 |
| | GO.0000022 | mitotic spindle elongation | 1.41E-7 | 5 |
| | GO.0022607 | cellular component assembly | 2.55E-7 | 33 |
| | GO.0007018 | microtubule-based movement | 3.19E-7 | 12 |

**Figure S14. GO process networks for EBV$^+$ activation intermediate (c0) and hyperproliferation (c6).**

Cluster 1: expression markers

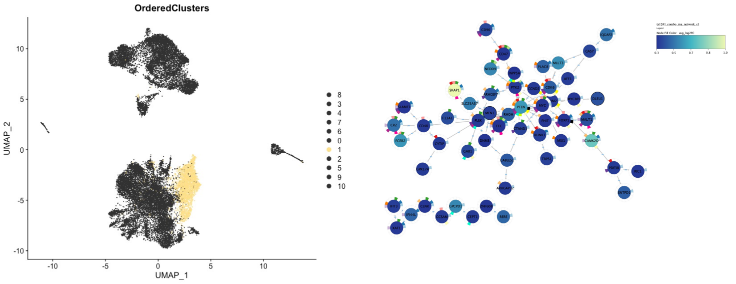


| | GO ID | GO process | FDR | # genes |
|---|---|---|---|---|
| | GO.0065007 | biological regulation | 9.23E-5 | 51 |
| | GO.0002376 | immune system process | 7.3E-4 | 21 |
| | GO.0007229 | integrin-mediated signaling pathway | 9.1E-4 | 5 |
| | GO.0007166 | cell surface receptor signaling pathway | 9.1E-4 | 19 |
| | GO.0002682 | regulation of immune system process | 0.0026 | 14 |
| | GO.0045655 | regulation of monocyte differentiation | 0.0032 | 3 |
| | GO.0051239 | regulation of multicellular organismal process | 0.0039 | 20 |
| | GO.1905809 | negative regulation of synapse organization | 0.0041 | 3 |
| | GO.0030155 | regulation of cell adhesion | 0.0048 | 9 |
| | GO.0035556 | intracellular signal transduction | 0.0051 | 14 |
| | GO.0042127 | regulation of cell population proliferation | 0.0077 | 14 |
| | GO.0098542 | defense response to other organism | 0.0092 | 10 |
| | GO.0048585 | negative regulation of response to stimulus | 0.0118 | 13 |
| | GO.0080134 | regulation of response to stress | 0.0125 | 12 |
| | GO.0006650 | glycerophospholipid metabolic process | 0.0188 | 6 |
| | GO.0070493 | thrombin-activated receptor signaling pathway | 0.0195 | 2 |
| | GO.0046855 | inositol phosphate dephosphorylation | 0.0195 | 2 |
| | GO.0010666 | positive regulation of cardiac muscle cell apoptotic process | 0.0217 | 2 |
| | GO.0002429 | immune response-activating cell surface receptor signaling pat... | 0.0226 | 5 |
| | GO.1903362 | regulation of cellular protein catabolic process | 0.0239 | 5 |
| | GO.0071375 | cellular response to peptide hormone stimulus | 0.0239 | 5 |
| | GO.0030097 | hemopoiesis | 0.0239 | 7 |
| | GO.0051270 | regulation of cellular component movement | 0.0246 | 9 |
| | GO.2000045 | regulation of G1/S transition of mitotic cell cycle | 0.0262 | 4 |
| | GO.0051093 | negative regulation of developmental process | 0.0275 | 9 |
| | GO.0002252 | immune effector process | 0.0285 | 9 |
| | GO.0022603 | regulation of anatomical structure morphogenesis | 0.0303 | 9 |
| | GO.0006796 | phosphate-containing compound metabolic process | 0.0304 | 14 |
| | GO.0002902 | regulation of B cell apoptotic process | 0.0304 | 2 |

Cluster 2: expression markers

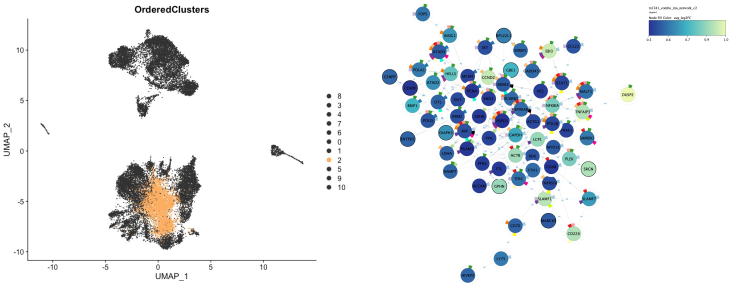


| | GO ID | GO process | FDR | # genes |
|---|---|---|---|---|
| | GO.0002376 | immune system process | 2.35E-8 | 31 |
| | GO.0006260 | DNA replication | 3.8E-6 | 10 |
| | GO.0019674 | NAD metabolic process | 4.89E-6 | 7 |
| | GO.0080090 | regulation of primary metabolic process | 1.49E-5 | 43 |
| | GO.0080134 | regulation of response to stress | 1.79E-5 | 19 |
| | GO.0050776 | regulation of immune response | 1.79E-5 | 16 |
| | GO.0042981 | regulation of apoptotic process | 5.73E-5 | 19 |
| | GO.0008283 | cell population proliferation | 5.73E-5 | 13 |
| | GO.0010033 | response to organic substance | 8.3E-5 | 26 |
| | GO.0044419 | interspecies interaction between organisms | 8.47E-5 | 13 |
| | GO.0001817 | regulation of cytokine production | 9.31E-5 | 12 |
| | GO.0045321 | leukocyte activation | 1.2E-4 | 14 |
| | GO.0042542 | response to hydrogen peroxide | 1.4E-4 | 6 |
| | GO.0046651 | lymphocyte proliferation | 2.8E-4 | 5 |
| | GO.0002822 | regulation of adaptive immune response based on somatic recombination of immune receptors ... | 3E-4 | 6 |
| | GO.0033209 | tumor necrosis factor-mediated signaling pathway | 3.5E-4 | 5 |
| | GO.0071897 | DNA biosynthetic process | 5.8E-4 | 5 |
| | GO.0016043 | cellular component organization | 6.3E-4 | 34 |
| | GO.0042770 | signal transduction in response to DNA damage | 7.6E-4 | 5 |
| | GO.0019220 | regulation of phosphate metabolic process | 7.9E-4 | 17 |
| | GO.0051707 | response to other organism | 8.6E-4 | 14 |
| | GO.0030029 | actin filament-based process | 9.9E-4 | 9 |
| | GO.0002833 | positive regulation of response to biotic stimulus | 9.9E-4 | 7 |
| | GO.0002250 | adaptive immune response | 0.001 | 7 |
| | GO.2000349 | negative regulation of CD40 signaling pathway | 0.0011 | 2 |
| | GO.0051240 | positive regulation of multicellular organismal process | 0.0011 | 16 |
| | GO.0050864 | regulation of B cell activation | 0.0011 | 5 |
| | GO.0009628 | response to abiotic stimulus | 0.0011 | 13 |
| | GO.1901992 | positive regulation of mitotic cell cycle phase transition | 0.0014 | 4 |
| | GO.0071157 | negative regulation of cell cycle arrest | 0.0016 | 3 |
| | GO.0006272 | leading strand elongation | 0.0017 | 2 |
| | GO.0051726 | regulation of cell cycle | 0.0018 | 13 |
| | GO.0009894 | regulation of catabolic process | 0.0021 | 11 |
| | GO.0044283 | small molecule biosynthetic process | 0.0022 | 9 |
| | GO.0016445 | somatic diversification of immunoglobulins | 0.0024 | 3 |
| | GO.0051235 | maintenance of location | 0.0026 | 5 |
| | GO.0008284 | positive regulation of cell population proliferation | 0.0028 | 11 |

**Figure S15. GO process networks for EBV$^+$ naïve-linked intermediate (c1) and NF-κB activation (c2).**

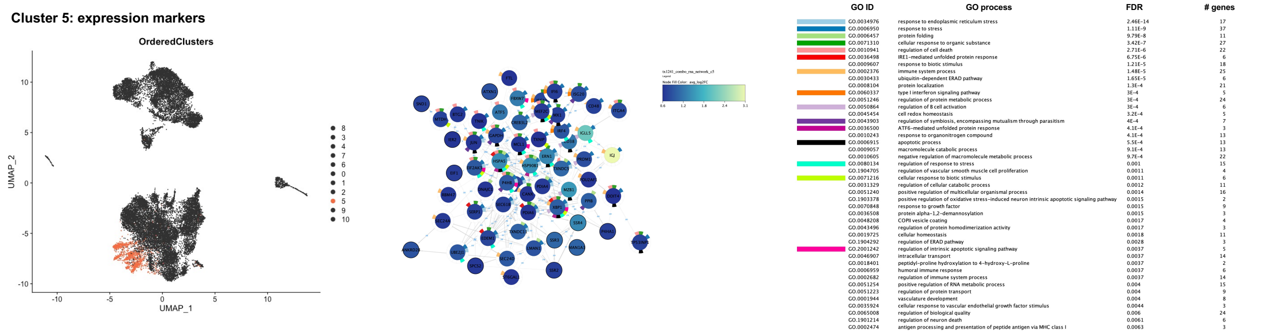


**Figure S16. GO process network for EBV$^+$ differentiated plasmablast state (c5).**

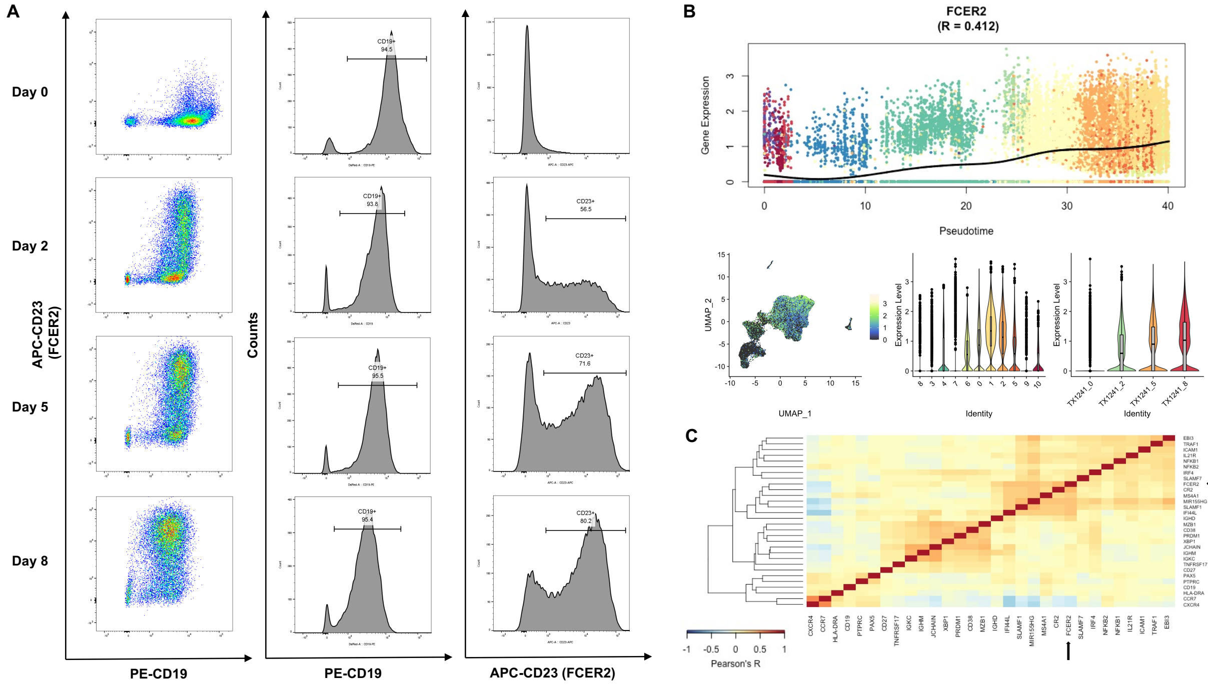


**Figure S17. Flow cytometry validation and expression dynamics of EBV-induced *FCER2* (CD23).**

(A) Protein-level validation of FCER2/CD23 upregulation following EBV infection of enriched B cells, assayed via CD19 positivity. CD19$^+$ fractions accounted for >93% of cells at all timepoints. FACS data (shown for TX1241) were consistent across donors.

(B) Expression of *FCER2* mRNA in pseudotime, as measured from scRNA-seq. A spline interpolant fit is shown for expression in pseudotime. *FCER2* mRNA expression is also presented in UMAP and violin plots to visualize cluster- and time-resolved variation.

(C) Correlation heatmap (Pearson) for *FCER2* relative to other genes of interest.

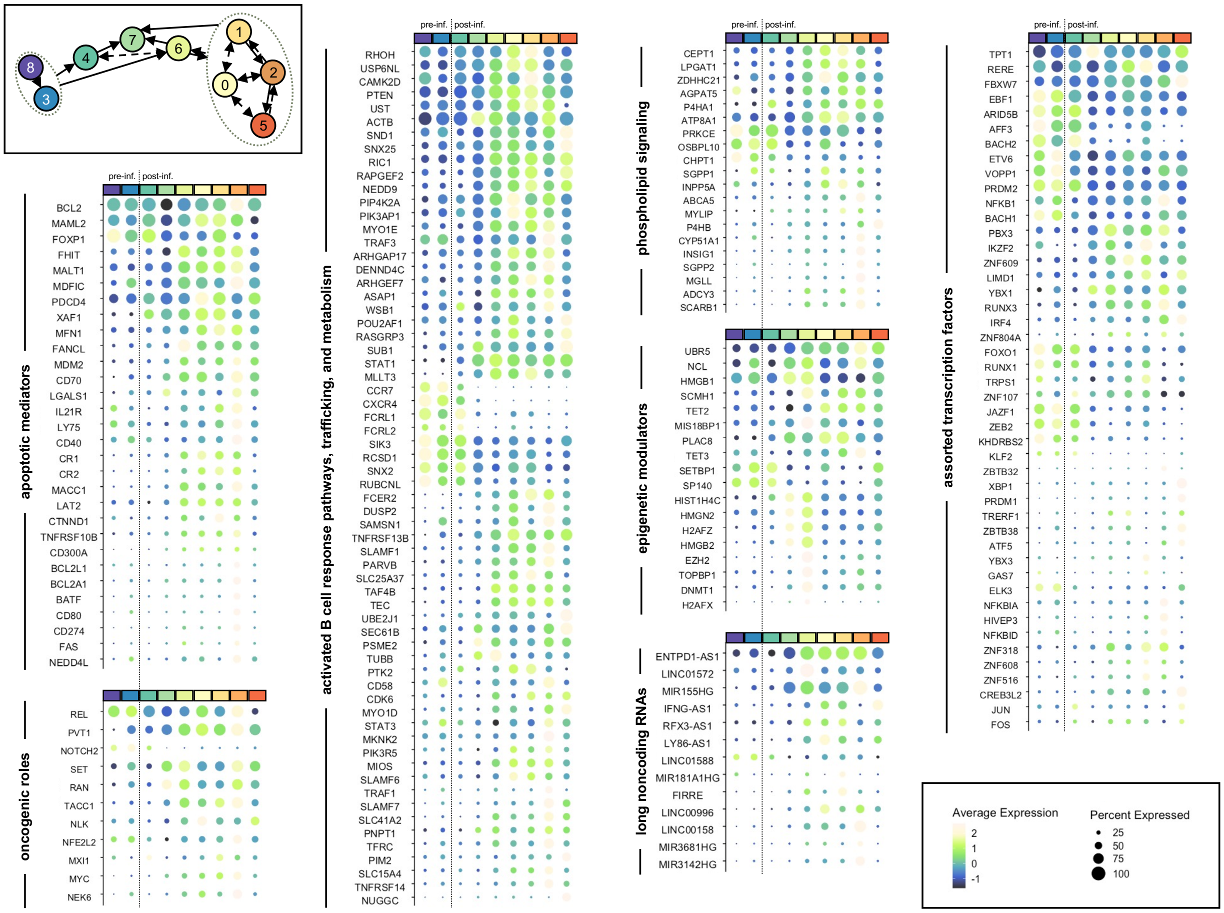


**Figure S18. Gene sets of interest resolved by model phenotype.**

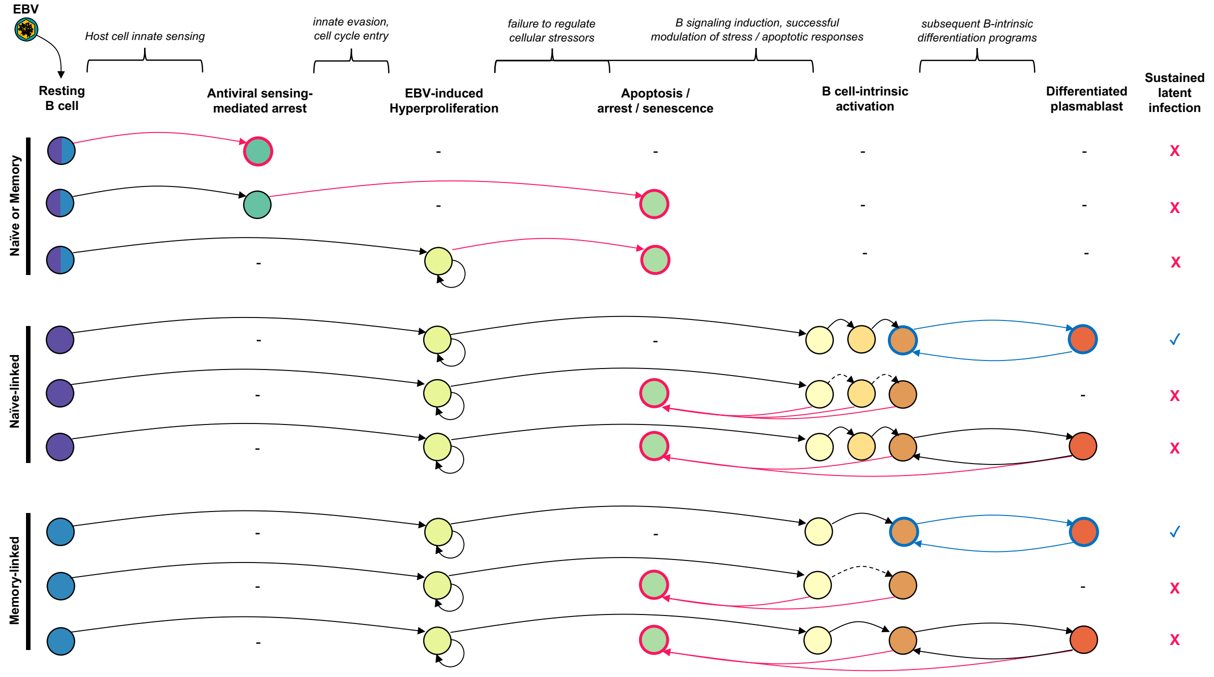


**Figure S19. Example B cell fate trajectories accounted for within the model of early EBV infection.**

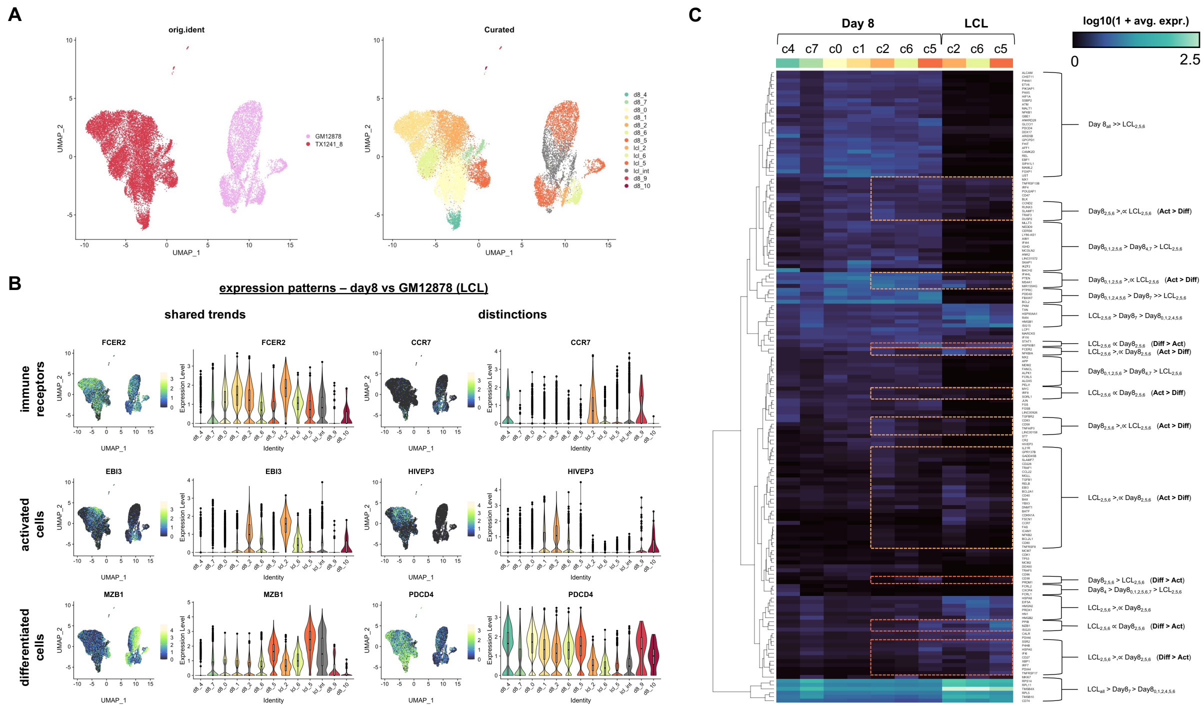


**Figure S20. Integration of Day 8 scRNA-seq with LCL scRNA-seq (GM12878).**
(A) Dimensionally reduced projection of merged Day 8 (red) + GM12878 (pink) scRNA-seq dataset with mapped expression phenotypes. Phenotype color coding is preserved for ease of interpretation.
(B) Comparison of activated and differentiated phenotypes (c2 and c5) for select genes of interest in early infection versus LCL. Day 8 and LCL samples share similarities with respect to characteristic markers of activated versus differentiated phenotypes (e.g., *EBI3* in activated cells and *MZB1* in differentiated cells). Notably, expression of *CCR7* is ablated at Day 8 but eventually recovers within the activated cell state within LCLs (Messinger et al., 2019). Conversely, the activation marker *HIVEP3* is expressed at Day 8 but absent within the LCL activation phenotype. A similar difference between Day 8 and LCL differentiated phenotypes is observed for the tumor suppressor *PDCD4*.
(C) Heatmap of phenotype-resolved gene expression patterns between Day 8 and LCL datasets. Genes (rows) are hierarchically clustered by pattern. Select gene sets are annotated based on their shared or divergent signatures between activation and differentiation phenotypes in Day 8 versus GM12878.

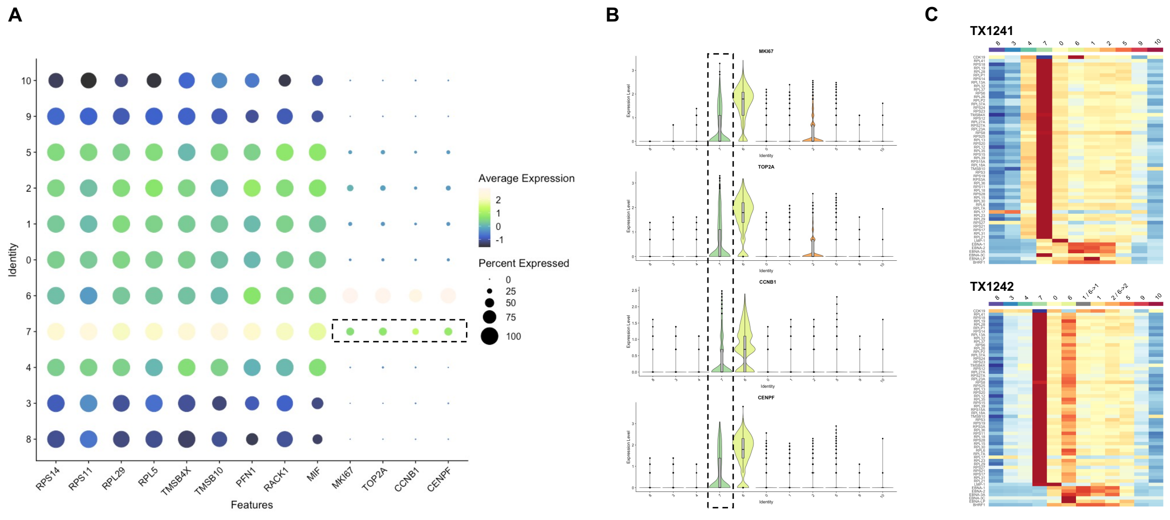


**Figure S21. Expression of stress-associated genes and cell cycle markers in c7.**
(A) Average and cluster fractional expression of c7 markers (ribosomal, actin sequestration, apoptotic, and cell cycle genes). Cells in c7 uniformly express high levels of ribosomal subunit genes, actin sequestering stress response proteins, and select apoptotic regulators. However, only 25-30% of cells in c7 express elevated cell cycle markers.
(B) Violin plots for select cell cycle markers. Cells in c7 exhibits the second-highest overall expression of cell cycle markers, superseded only by hyperproliferative cells (c6).
(C) Average expression of the top 40 markers (and viral genes of interest) for cluster 7 in both donors.

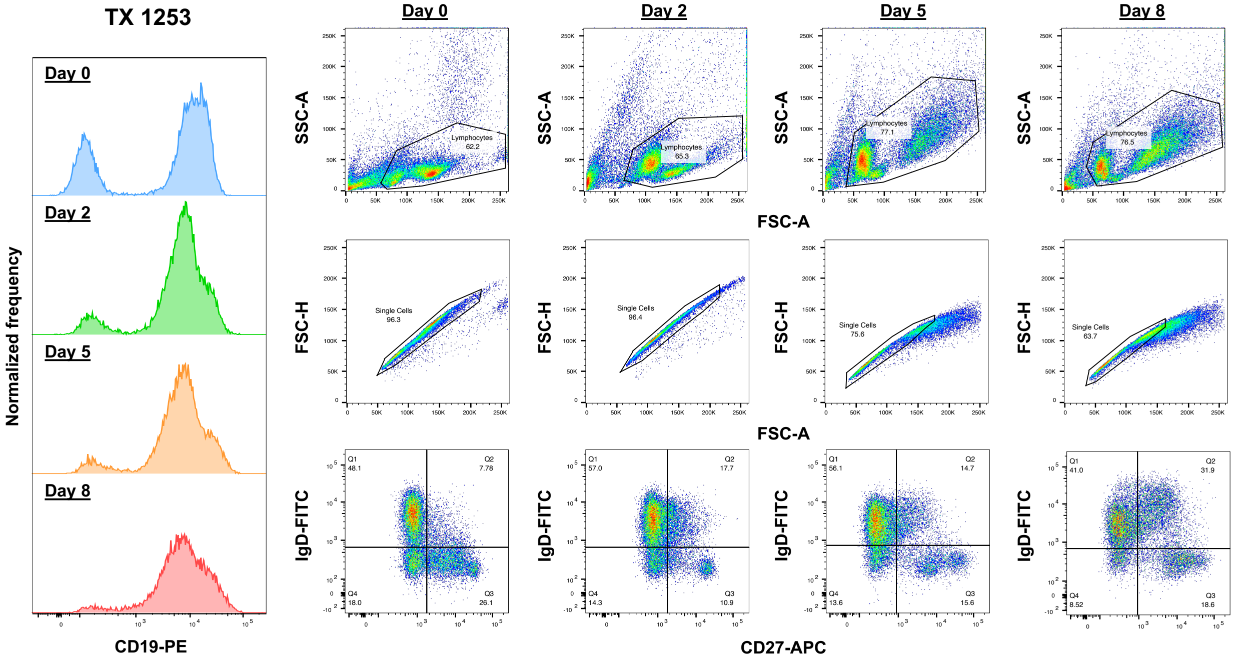


**Figure S22. B cell enrichment, gating strategy, and naïve and memory status during early infection in a third donor.**

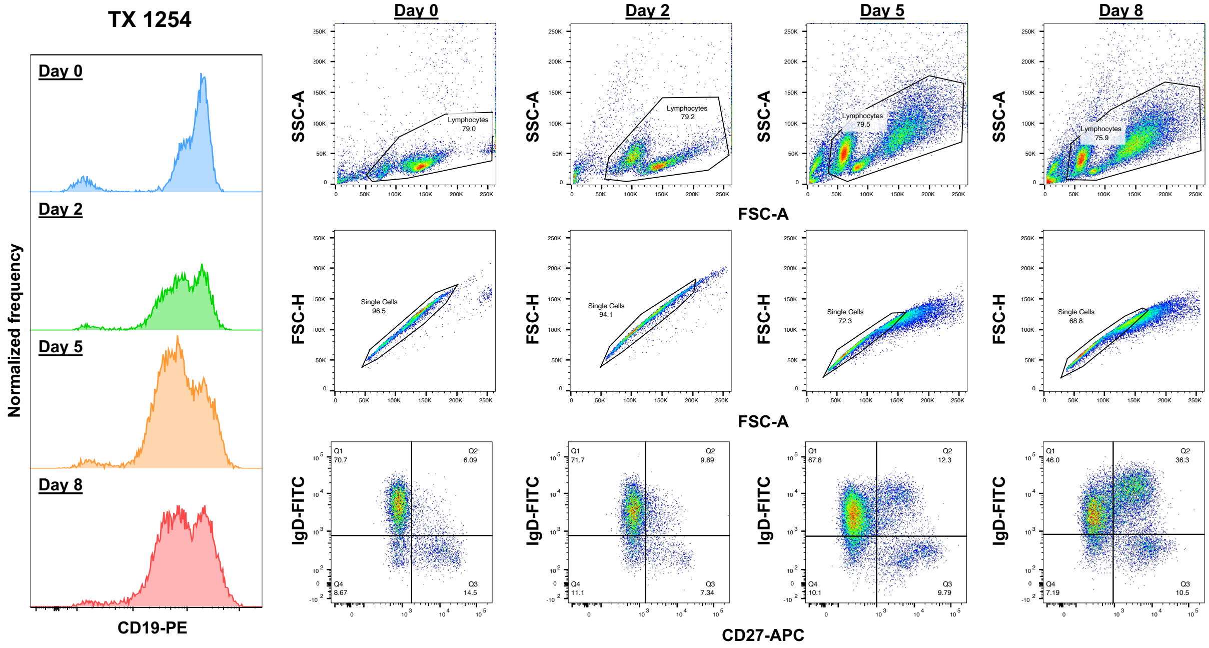


**Figure S23. B cell enrichment, gating strategy, and naïve and memory status during early infection in a fourth donor.**

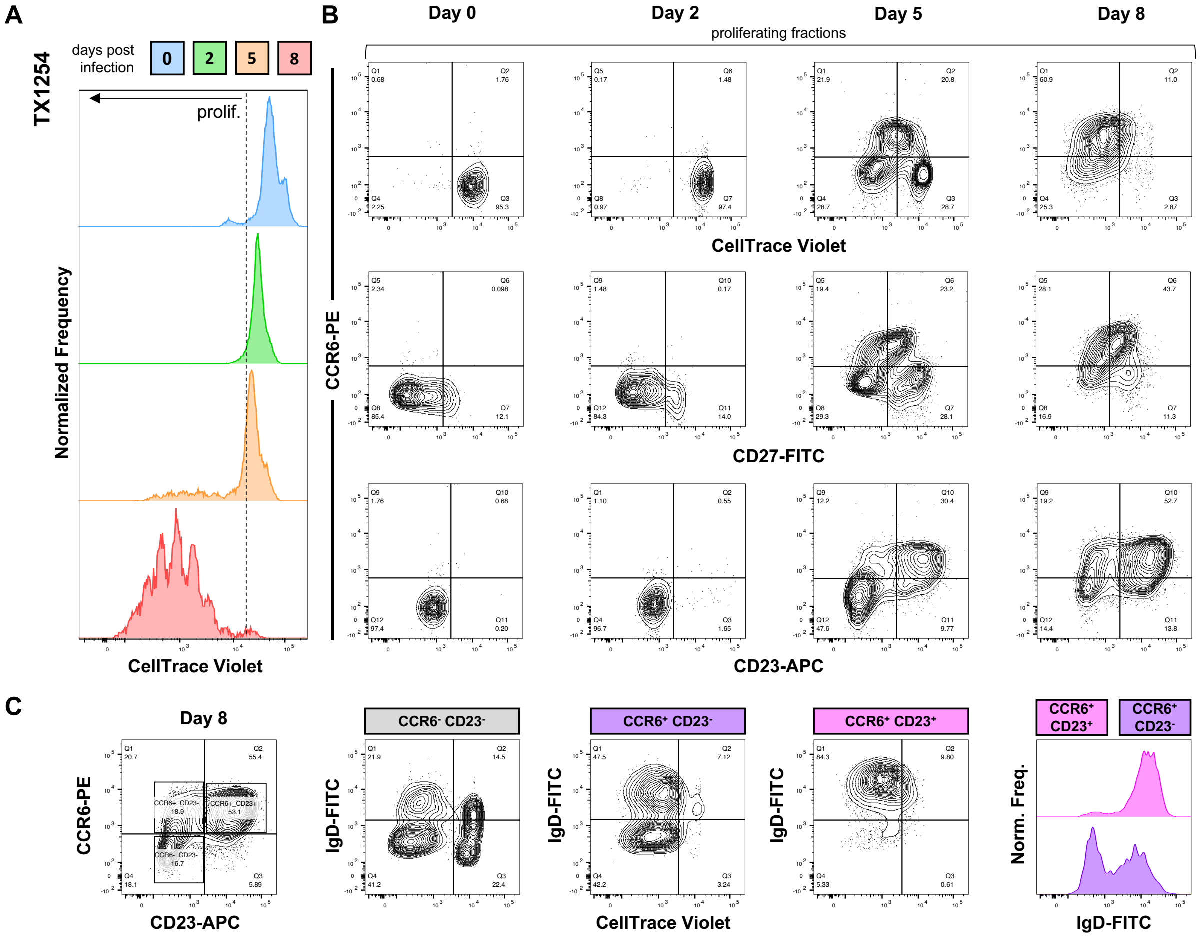


**Figure S24. Analysis of CCR6$^{+}$ AP-eMBC cells in proliferating fractions during early infection in the fourth donor.**
(A) CellTrace Violet staining to track cell proliferation following EBV infection.
(B) Time-resolved CCR6 status versus proliferation (top row), memory (CD27, middle row), and EBV-induced activation (CD23/FCER2, bottom row).
(C) Naïve vs memory and proliferative status of CCR6$^{+}$ cells by CD23 status.

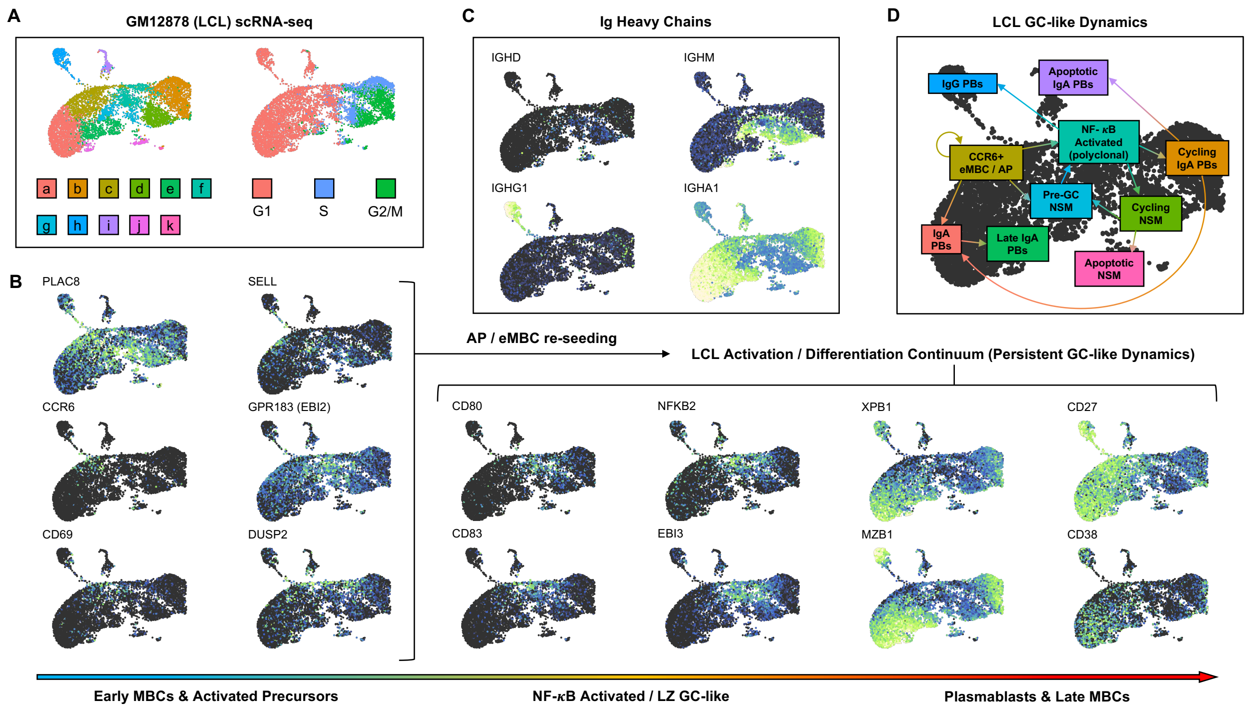


**Figure S25. An AP-eMBC phenotype is retained in EBV-immortalized LCLs.**
(A) Cluster and cell cycle scoring UMAPs of GM12878 scRNA-seq.
(B) Ordered progression of AP-like expression markers through GC BC and post-GC effector phenotypes.
(C) Immunoglobulin heavy chain classes in GM12878.
(D) Proposed network of annotated phenotypes and transitions in LCL based on early infection data and GC dynamics.

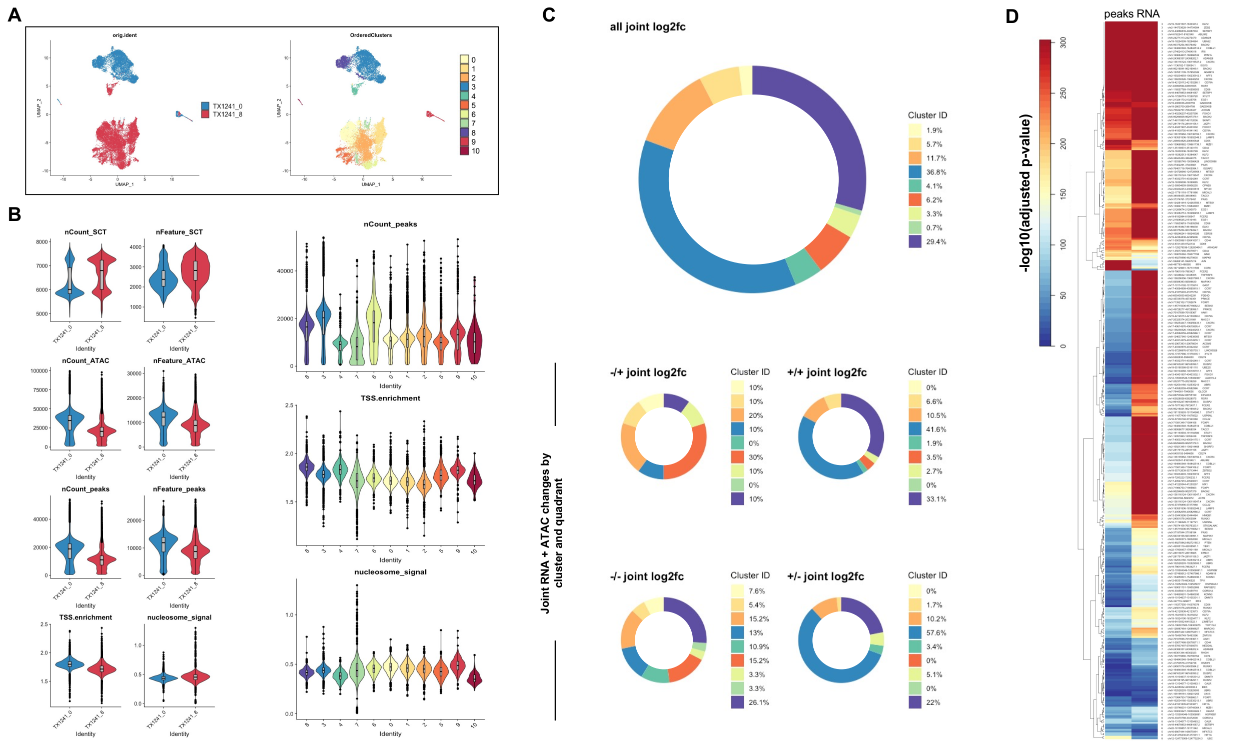


**Figure S26. Joint multimodal assay QC and DAP-linked DEG pattern summary by cluster.** (A) UMAP of integrated scRNA + scATAC data color-coded by day and cell phenotype.
(B) QC and global trends in multimodal single-cell assay. SCT = RNA expression data normalized using sctransform R package, (Hafemeister and Satija, 2019); ATAC = Tn5 insertion sites; peaks = MACS2 called ATAC peaks; TSS = transcription start site; nCount_peaks = number of total peaks; nFeature_peaks = number of unique peaks; nucleosome_signal < 1 for QC filter.
(C) Phenotypic summary of all-vs-one DAP-linked DEGs. All identified DAP-linked DEGs are summarized (top panel) and broken down by observed joint regulatory patterns, expressed as the directionality of DAP and DEG changes within a given phenotype (e.g., -/+ = decreased accessibility and upregulated expression within clusters specified by color coding).
(D) Log-transformed adjusted p values for DAPs (and their linked gene) by cluster, as calculated in Signac.

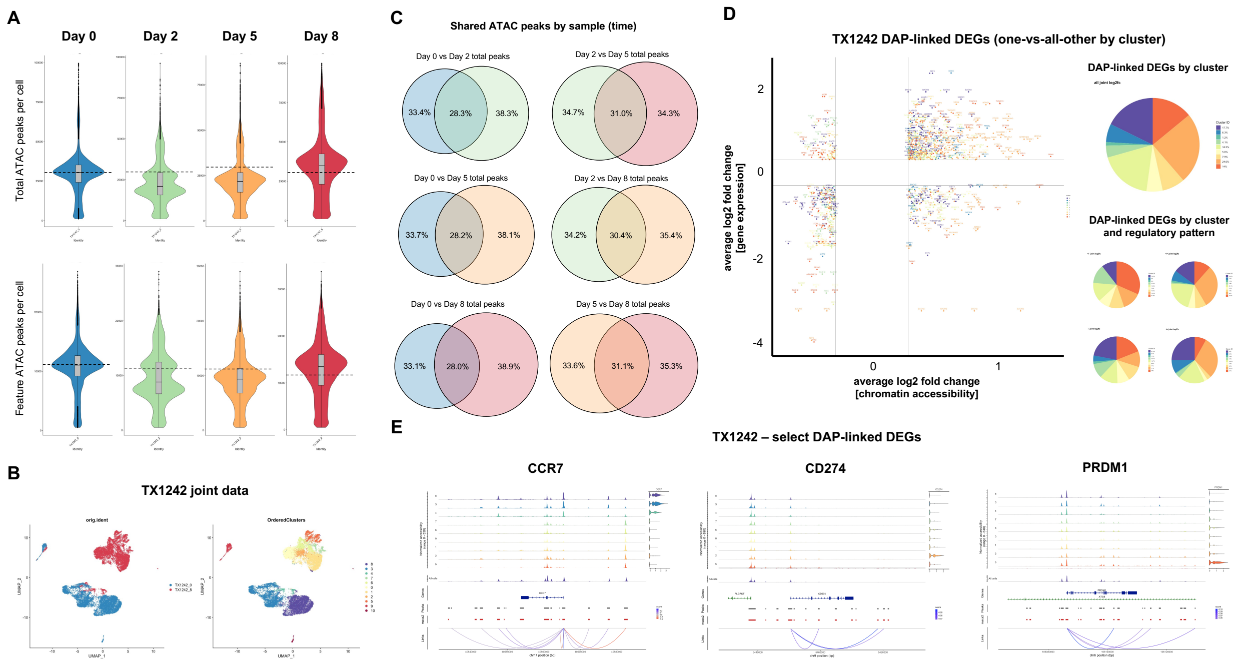


**Figure S27**. **TX1242 ATAC summary and joint multimodal analysis**.
(A) Global changes in ATAC peaks and features per cell by day. Dashed lines demarcate the mean peak per cell at Day 0.
(B) Sample- and cluster-resolved joint multimodal UMAP for TX1242.
(C) Overview of shared ATAC peaks between all pairs of sampled timepoints.
(D) Overview of significant DAP-linked DEGs and joint regulatory patterns.
(E) Joint multimodal plots for select DAP-linked DEGs (also identified in TX1241).

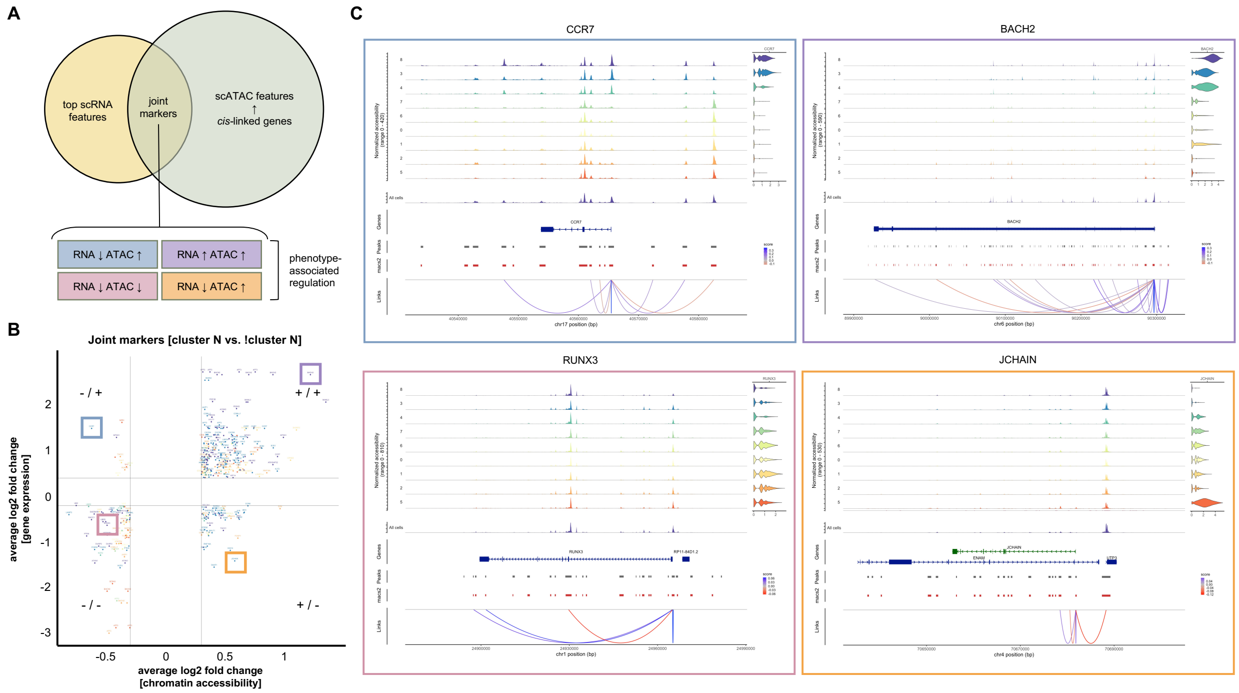


**Figure S28. Example DAP-linked DEGs with variable joint regulatory patterns.**
(A) Schematic of joint marker identification (DAPs linked to DEGs found via one-vs-all other phenotype comparisons) and the four regulatory patterns observed in DAP-linked DEGs.
(B) Scatterplot of significant DAP/DEG linkages by phenotype. Gene expression log2 fold changes (y axis) are plotted against accessibility log2 fold changes (x axis) and color-coded by cluster. Each point corresponds to a DAP. Multiple DAPs may be linked to the same DEG.
(C) Genes of interest representing each of the four joint regulatory patterns. In many cases, a given DAP-linked DEG exhibits multiple regulatory patterns (e.g., multiple peaks, including positive and negative regulatory linkages).

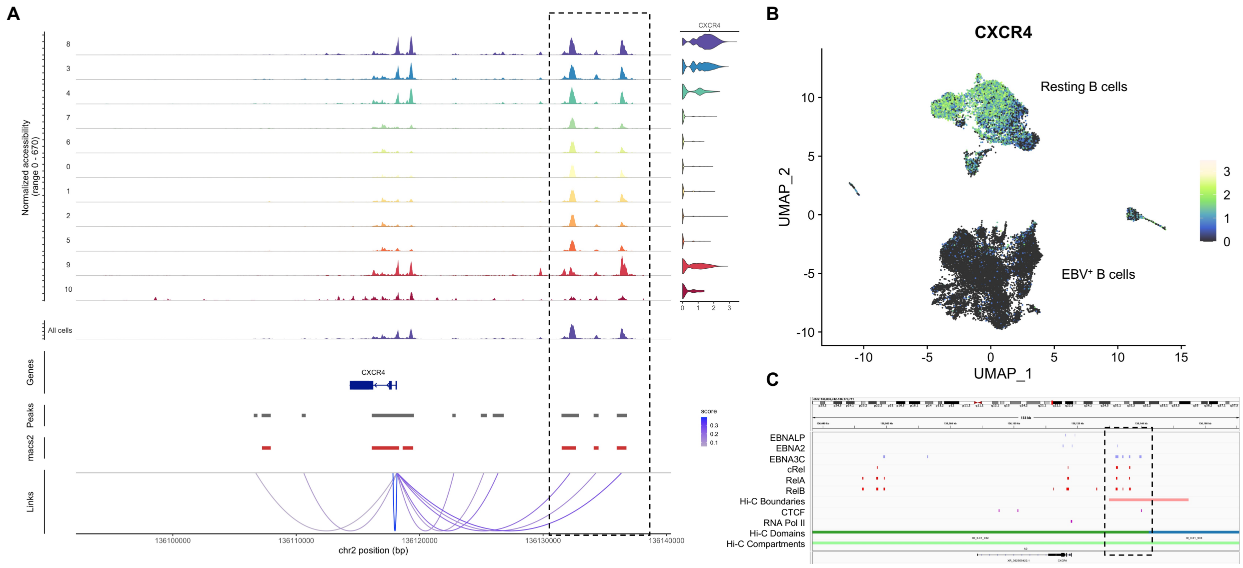


**Figure S29. *CXCR4* – a DAP-linked DEG with associated host and viral transcriptional co-activator binding sites.**
(A) Joint coverage plot for *CXCR4*. Phenotype-resolved scATAC profiles and matched scRNA expression data are depicted for the *CXCR4* locus. Statistically significant linkages to called peaks were identified at various points relative to the gene TSS.
(B) Expression of *CXCR4* visualized in integrated multimodal single-cell UMAP.
(C) Overlap of multiple upstream regulatory peaks with EBNA and Rel family binding sites. This region also corresponds with a local topologically associated domain (TAD) boundary within a euchromatic compartment based on alignment to ensemble Hi-C data for GM12878.

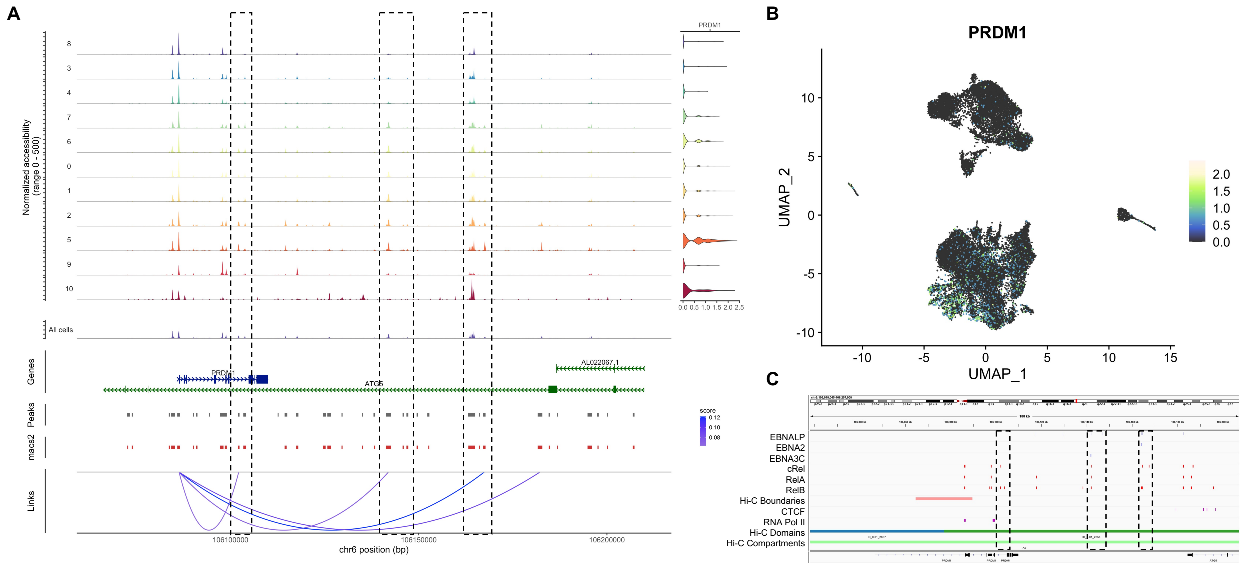


**Figure S30. *PRDM1* – a DAP-linked DEG with associated host and viral transcriptional co-activator binding sites.**

(A) Joint coverage plot for *PRDM1*. Phenotype-resolved scATAC profiles and matched scRNA expression data are depicted for the *PRDM1* locus. Statistically significant linkages to called peaks were identified at various points downstream of the gene TSS.

(B) Expression of *PRDM1* visualized in integrated multimodal single-cell UMAP.

(C) Overlap of multiple downstream regulatory peaks with EBNA and Rel family binding sites. This region is adjacent to a local topologically associated domain (TAD) boundary within a euchromatic compartment based on alignment to ensemble Hi-C data for GM12878.

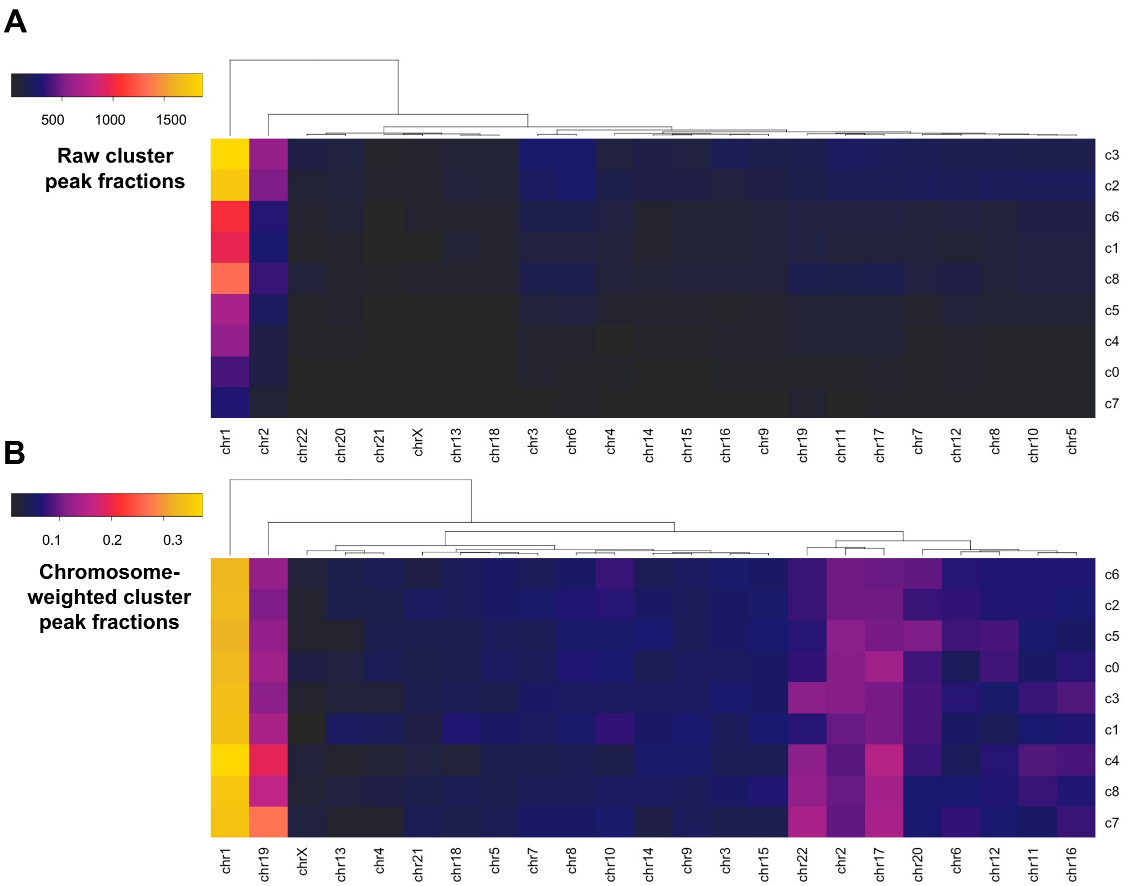


**Figure S31. Called peaks by chromosome and cell phenotype.**
(A) Raw peak counts by chromosome for each cluster. The majority of peaks are on large, gene-rich chromosomes (chr1, chr2).
(B) Chromosome length-normalized peak proportions by cluster. Raw peak counts were normalized by chromosome lengths (# of bases) and expressed as proportions of all peaks within a given phenotype (i.e., each row sum = 100% of peaks within the cluster). This visualization reveals that non-arrested $EBV^+$ phenotypes (c0, c1, c2, c5, and c6) exhibit reduced peak accessibility within chromosome 22 and increased accessibility within chromosome 20 relative to resting B cells. Notably, while overall accessibility decreases significantly in $EBV^+$ arrested states (c4 and c7), the only clear change in chromosome-resolved peak frequency is a slight increase in accessibility within chromosome 19 relative to resting cells.

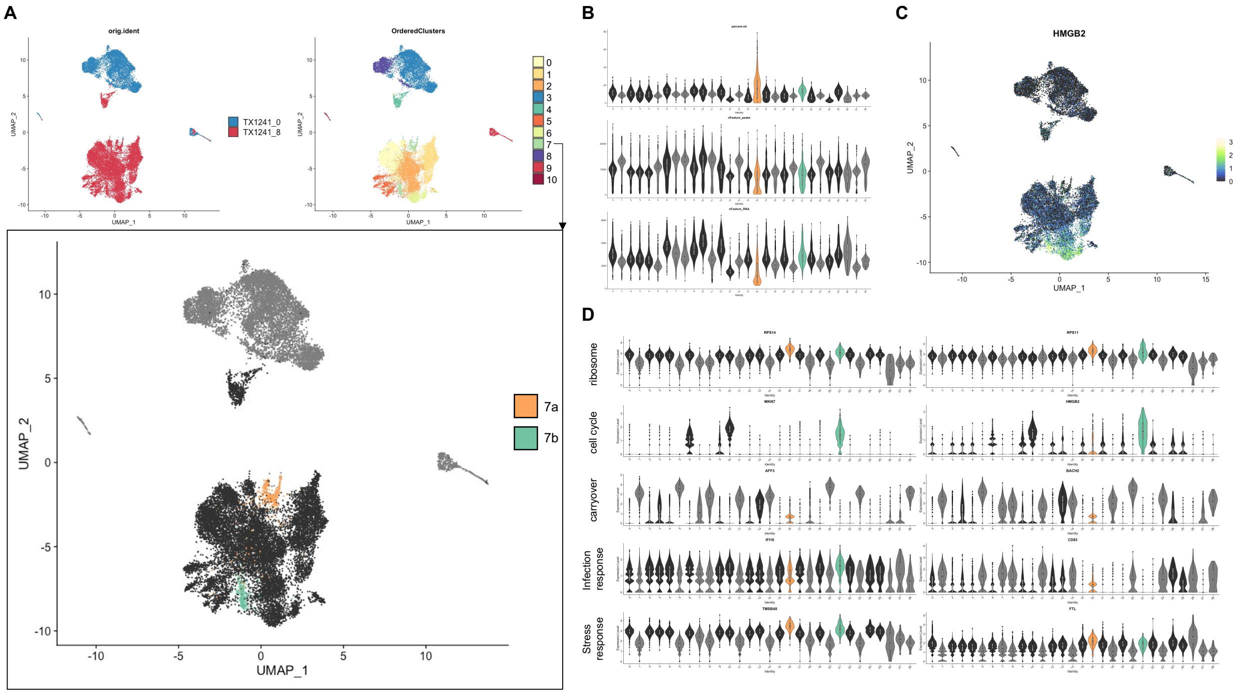


**Figure S32. High-resolution multimodal dissection of c7 (EBV$^{+}$ arrest phenotype).**
(A) UMAP of integrated scRNA + scATAC data. Cells color-coded by day (top left panel) and phenotype reveal two separate c7 populations within the 2D UMAP projection (top right panel). Upon higher resolution clustering, c7 separates into 7a (orange) and 7b (green) (bottom panel). All high resolution resting cell clusters and EBV$^{+}$ cell clusters not in 7a or 7b are also presented (gray and black cell groups, respectively).
(B) Mitochondrial expression, called peak features, and gene expression features within high-resolution clusters. 7a exhibits higher levels (and a broader range) of mitochondrial gene expression than 7b. Conversely, 7b exhibits more unique feature RNAs than 7a. Both 7a and 7b exhibit fewer feature ATAC peaks than resting cell phenotypes (gray).
(C) Joint UMAP of *HMGB2* expression, a gene involved in cytosolic DNA sensing and chromatin remodeling. Loss of *HMGB2* is implicated in heterochromatin formation. *HMGB2* transcript levels are markedly higher in 7b versus 7a.
(D) Expression of key marker genes by class within 7a versus 7b. Both 7a and 7b display elevated expression of ribosomal subunit genes involved in ribosome biogenesis-mediated senescence. Unlike 7a, 7b exhibits strong expression of cell cycle markers (consistent with its proximity to c6). Differential expression of *HMGB2* in 7a vs 7b implicates its roles in foreign DNA sensing (in 7a) and cell cycle-associated chromatin remodeling (7b). 7a exhibits expression of genes carried over from resting cells, while 7b does not, indicating earlier temporal occurrence of 7a (i.e., derivation from innate arrested cells (c4)). 7a also exhibits higher levels of *CD83* which is downregulated upon infection. 7a and 7b display similar elevated levels of stress response-associated genes (*TMSB4X*, *FTL*).

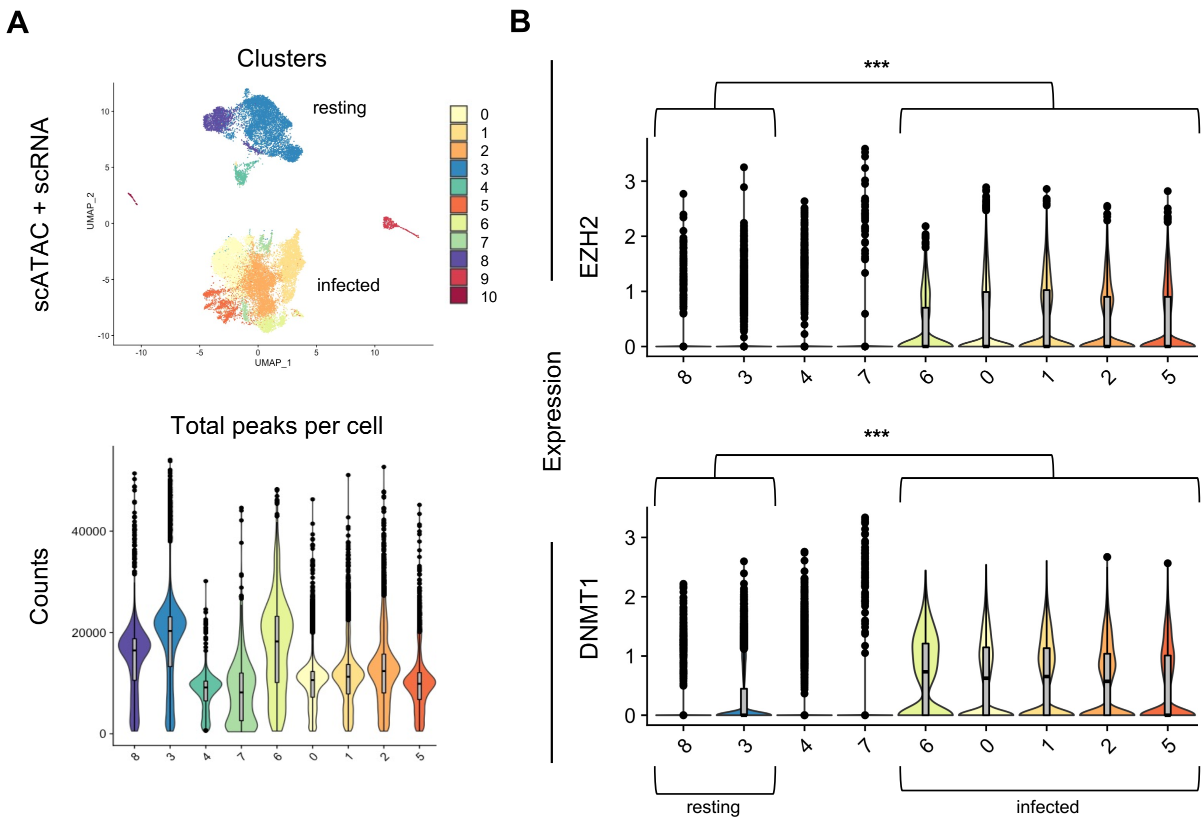


**Figure S33. Post-infection heterochromatin induction is concomitant with upregulated expression of genes involved in polycomb repression.**

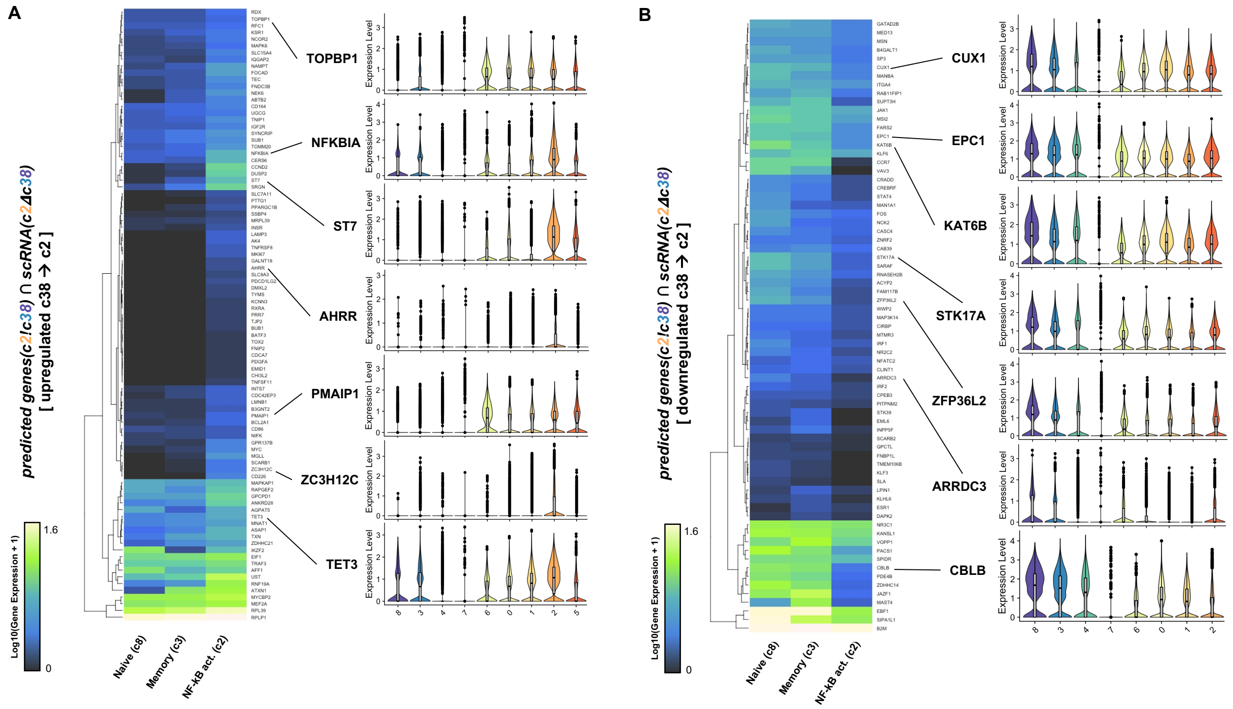


**Figure S34. Predicted *c2!c38* DAP-linked genes identified as *c2Δc38* DEGs (TX1241).**
(A) Predicted c2!c38 DAP-linked DEGs with *c38* → *c2* upregulated expression.
(B) Predicted c2!c38 DAP-linked DEGs with *c38* → *c2* downregulated expression.

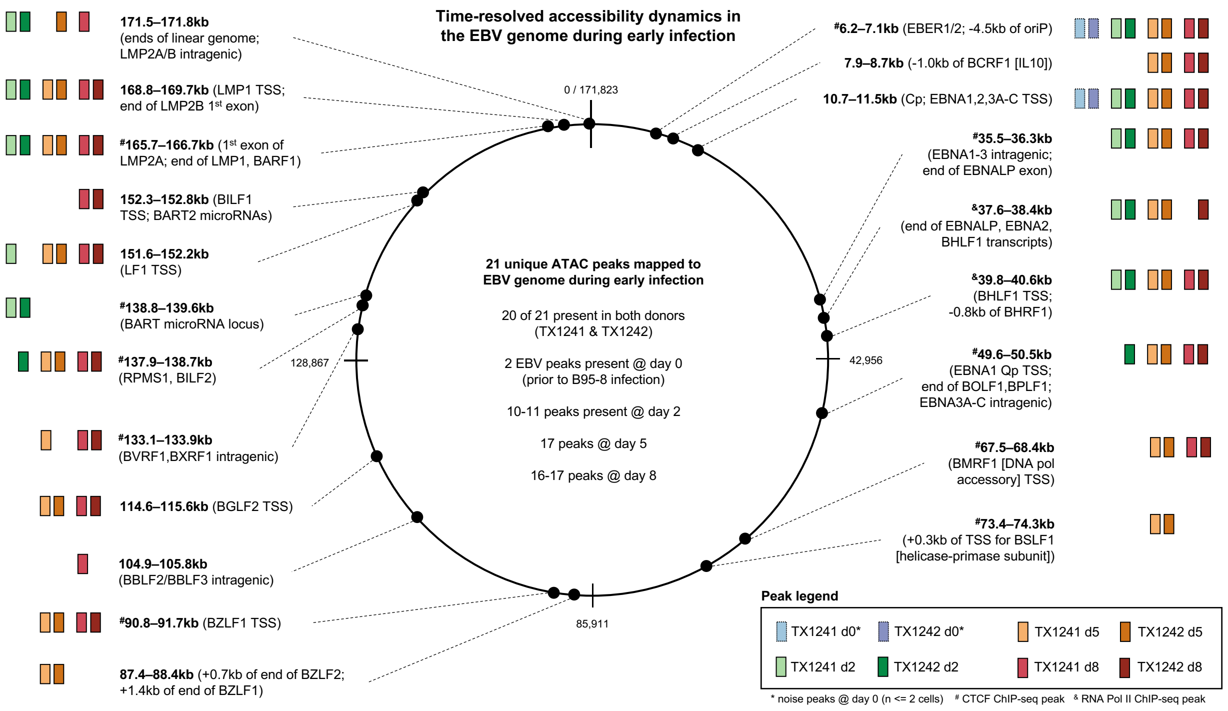


**Figure S35. Time-resolved accessibility dynamics in the EBV genome.**

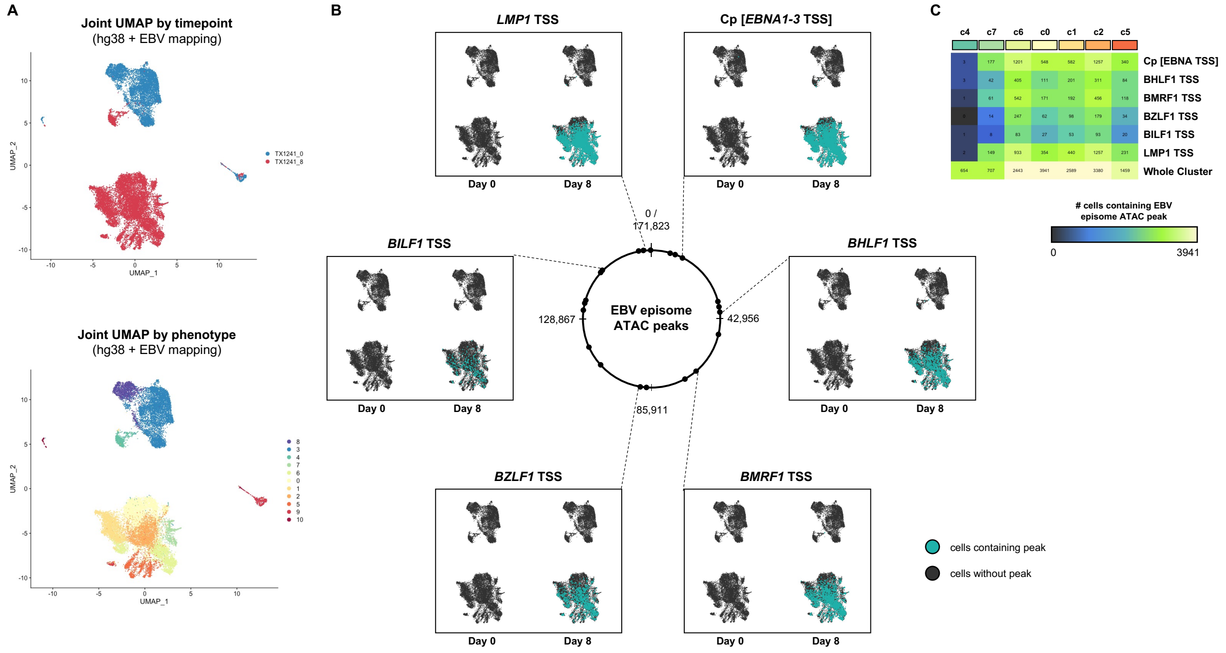


**Figure S36. Phenotype-resolved viral episome accessibility differences at key loci.**
(A) Sample- and cluster-resolved joint multimodal UMAP from data mapped to hg38+EBV reference.
(B) UMAP depiction of cells exhibiting viral episome accessibility at key loci of interest.
(C) Number of cells within each cluster (phenotype) exhibiting episomal ATAC peaks at the loci of interest in (B). The total number of cells per cluster is provided in the bottom row of the heatmap.

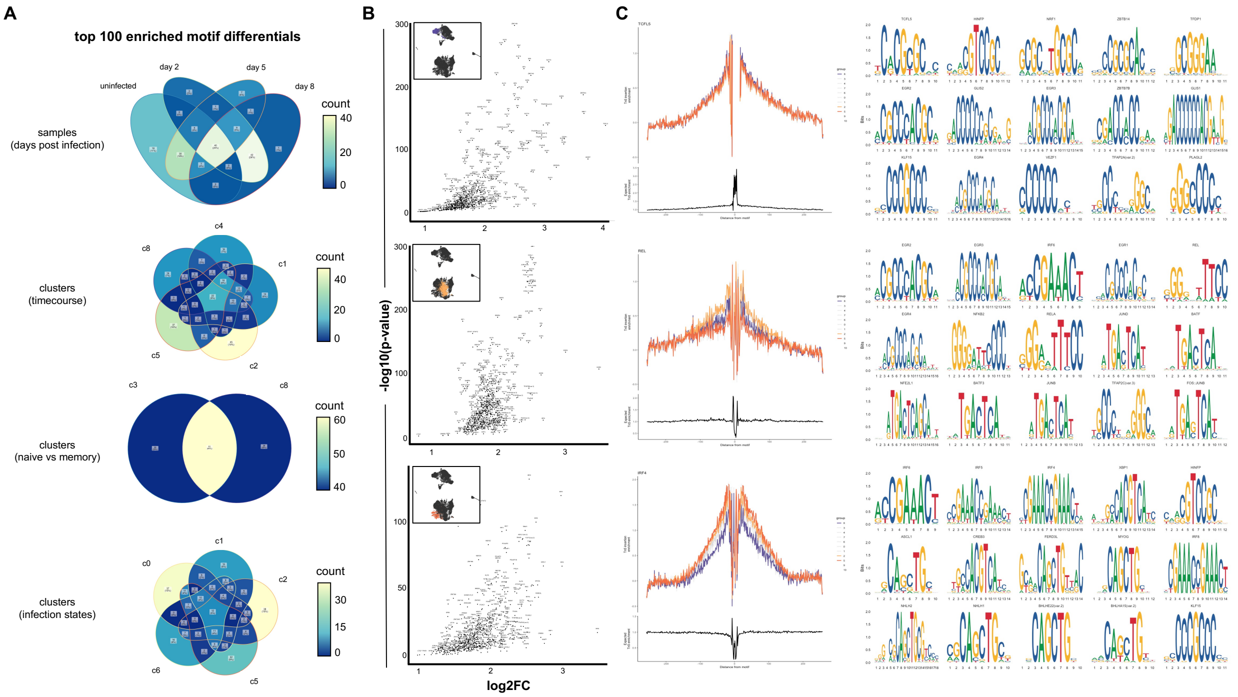


**Figure S37. Differential motif enrichment in ATAC peaks by timepoint and phenotype (TX1241).**
(A) Venn diagrams of differences in the top 100 enriched motifs by timepoint and phenotype.
(B) Volcano plots of enriched TF binding motifs for select clusters (top row = naïve cells (c8); middle row = activated cells (c2); bottom row = differentiated cells (c5)).
(C) Example enrichment of TF footprint by phenotype and the top 15 motifs per cluster.

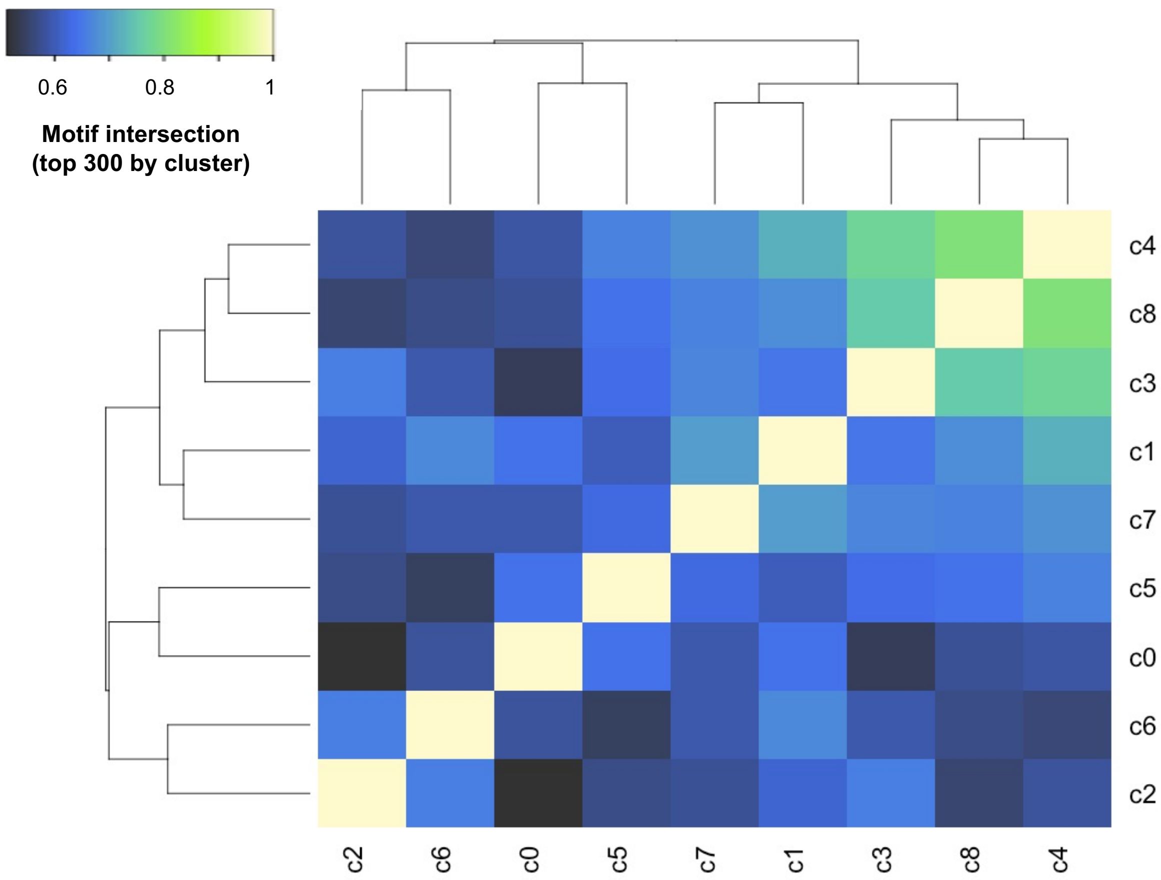


**Figure S38. TF binding motif correlation by phenotype.**

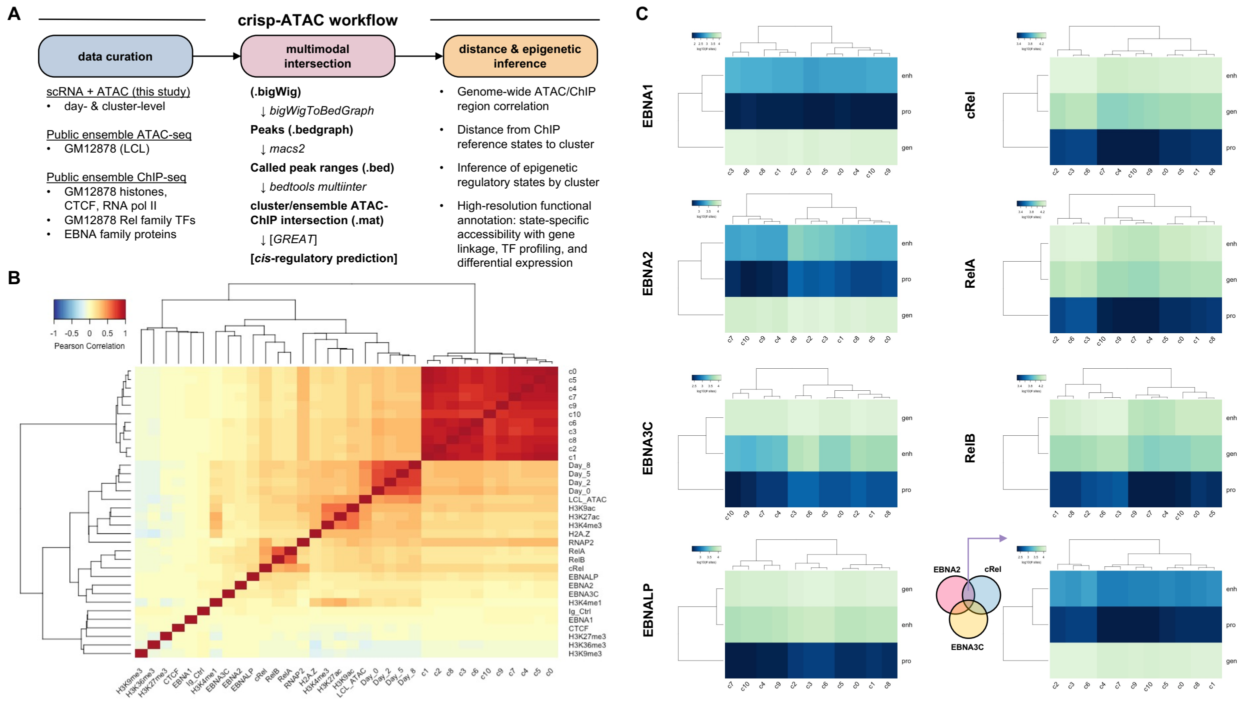


**Figure S39. crisp-ATAC workflow and initial testing.**
(A) Schematic of the inputs, steps, and outputs of crisp-ATAC.
(B) Correlation heatmap (Pearson) of scATAC, ensemble ATAC, and ensemble ChIP-seq datasets. Differences in correlation arise between modalities and single-cell versus ensemble measurements.
(C) Phenotype-resolved crisp-ATAC identification of enhancer (enh), promoters (pro) and actively transcribed genes (gen).

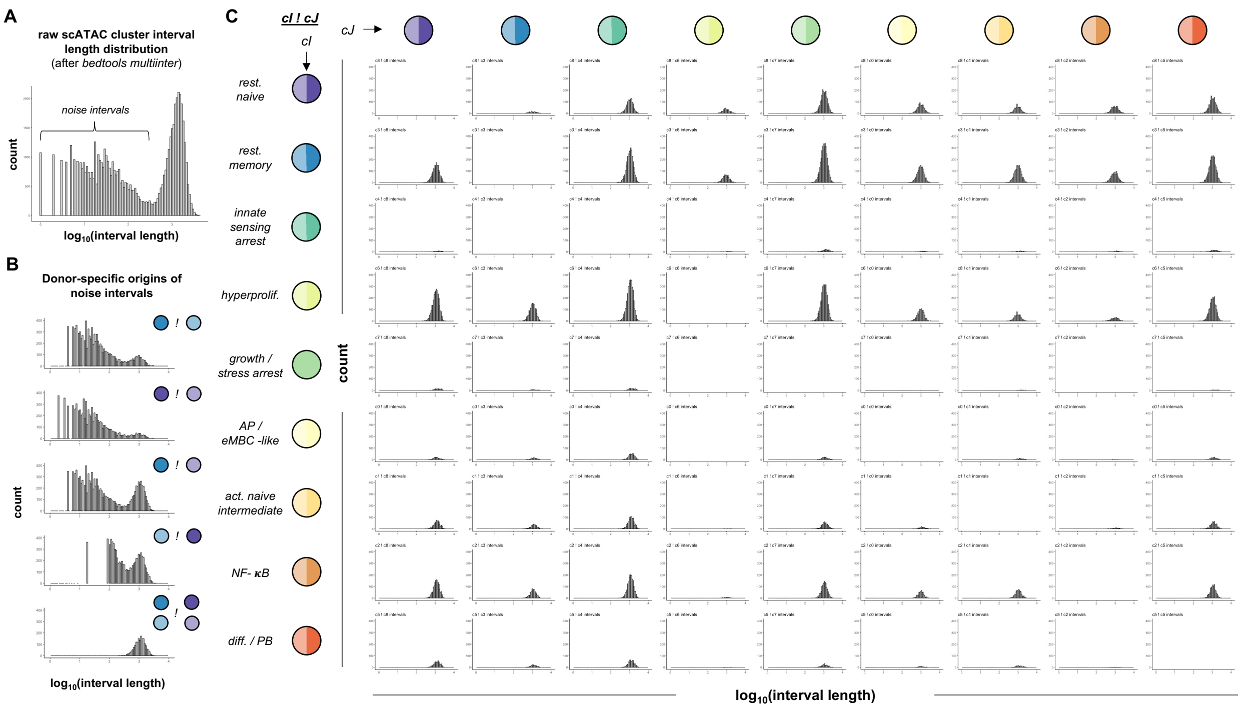


**Figure S40. crisp-ATAC intersection interval characterization across scATAC clusters.** (A) Interval size distribution resulting from multiple intersection of all scATAC cluster bed files across both biological replicates (donors). Called differential peak intervals less than ~100 nucleotides among clusters were considered as noise.
(B) Example highlighting donor variation in called peaks as the predominant source of noise intervals. In the example, memory cell ATAC peaks from donor 1 (dark blue) and donor 2 (light blue) were compared against each other as well as against naïve cell ATAC peaks from each donor (dark purple and light purple, respectively). Noise intervals are successfully filtered by constraining differential intervals to those consistent across both donors (bottom panel).
(C) Intersection peak interval length distributions and frequency across all pairwise cluster comparisons taking both donors into account. The x and y axes ranges are identical across the histograms to facilitate quantitative comparisons of peak frequency across phenotypes (distribution height). Across all cluster comparisons, the median length of a unique peak interval identified from crispATAC was ~1,150 nucleotides.

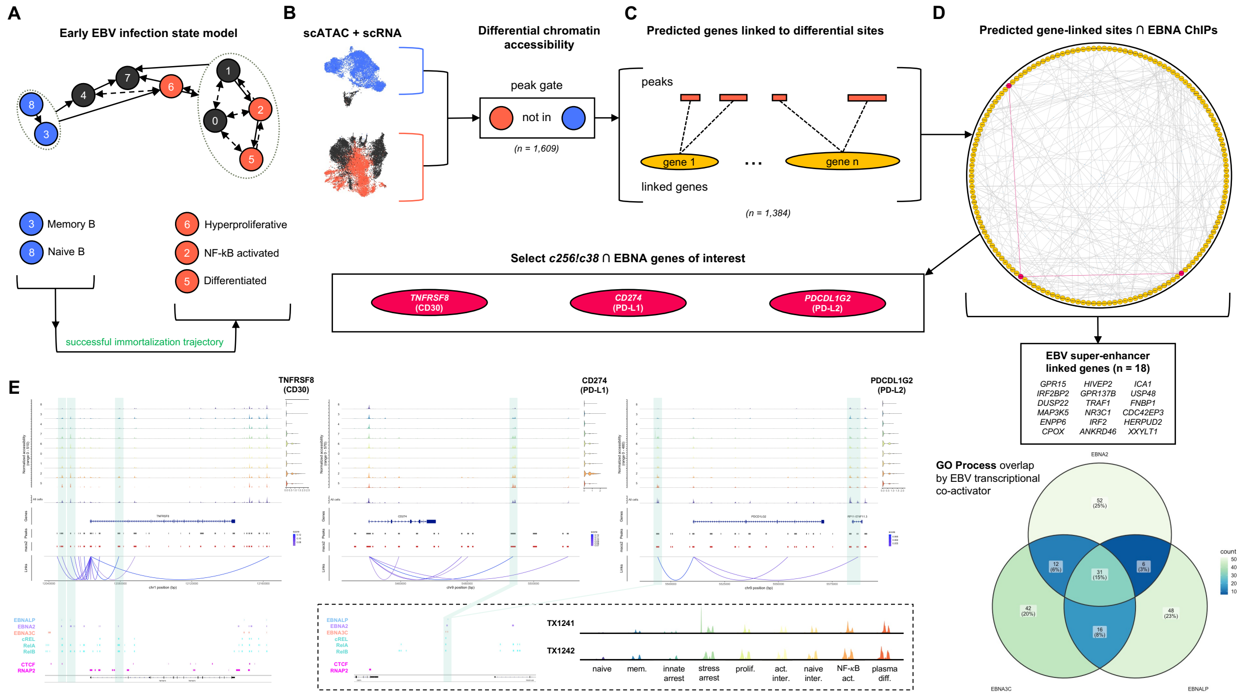


**Figure S41. crisp-ATAC analysis of DAP-linked DEGs associated with viral transcriptional co-activator binding sites in successful infection trajectory states.**
(A) Schematic overview of model states used in comparison (*c256 vs c38* peaks in both donors).
(B) Gating to identify *c256!c38* DAPs (n = 1,609).
(C) Prediction of DAP-linked genes (n = 1,384).
(D) GO network of *c256!c38* DAP-linked DEGs at EBNA-associated sites and overlap with EBVSE-linked genes (18 of 167). The Venn diagram depicts GO process enrichment by EBNA for associated DAP-linked DEGs.
(E) Joint coverage plots for each gene of interest (*PDCDL1G2, CD274,* and *TNFRSF8*) with EBNA-associated linked sites highlighted.

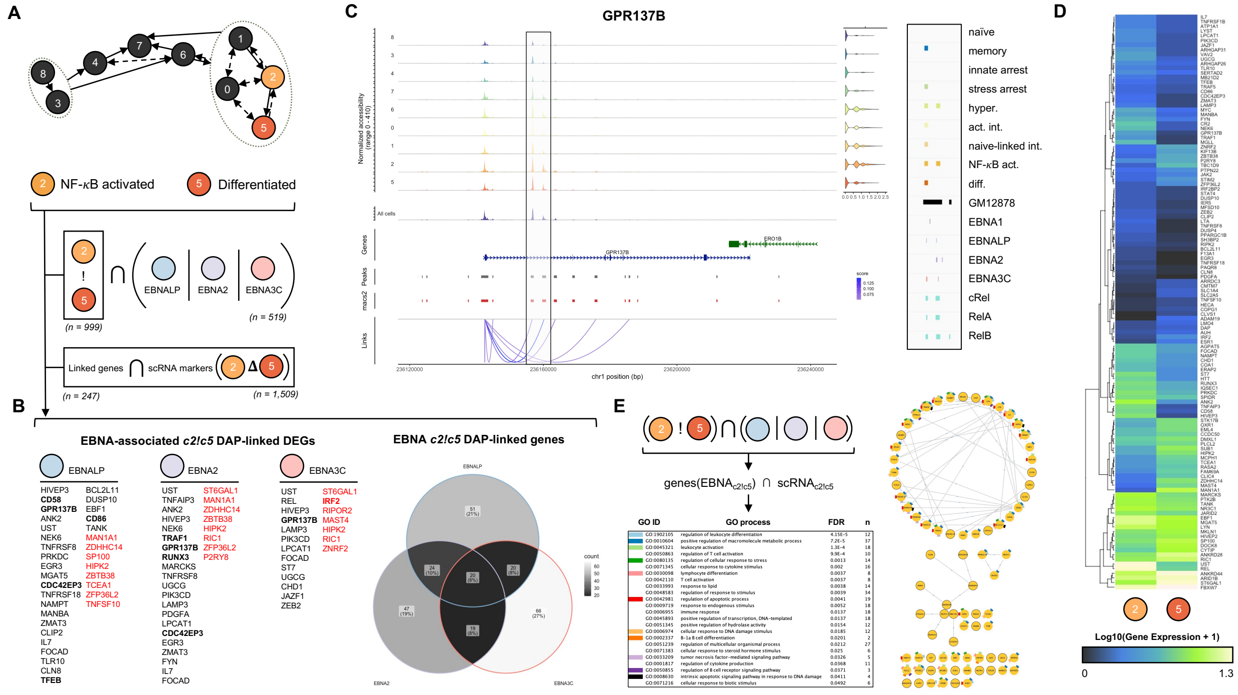


### Figure S42. crisp-ATAC analysis of EBNA site DAP-linked DEGs in activated versus differentiated EBV$^+$ B cells

(A) Schematic overview of model states used in comparison (*c2 vs c5* peaks in both donors) with gating to identify *c2!c5* DAPs (n = 999) overlapping at least one EBNA binding site (n = 519). Predicted DAP-linked genes (n = 247) were intersected with empirically measured DEGs (n = 1,509) from scRNA-seq.

(B) Identification of EBNA-associated c2!c5 DAP-linked DEGs with empirical validation. Bold text denotes EBVSE-linked genes. Black and red genes are respectively upregulated and downregulated along the *c2→c5* trajectory. The Venn diagram shows the frequency of single- and multiple- EBNA site overlaps for these genes.

(C) Joint accessibility and expression plot for *GPR137B*, an empirically validated EBNA-associated (EBVSE-linked) *c2!c5* DAP-linked DEG. Peak calls for scATAC + scRNA clusters as well as EBNA and Rel ChIP-seq at select significant linked sites are highlighted.

(D) Average expression of identified EBNA-associated DAP-linked DEGs between c2 and c5.

(E) GO process term enrichment and network for EBNA-associated *c2!c5* DAP-linked DEGs identified using crisp-ATAC.

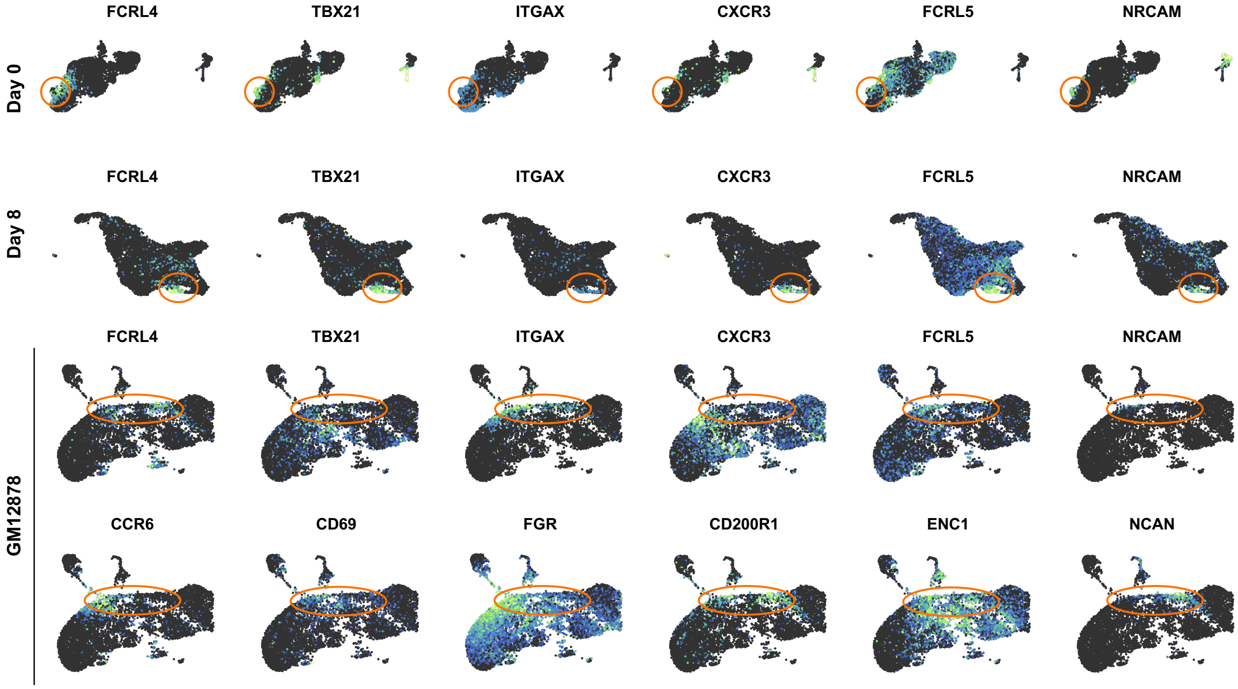


**Figure S43. Conservation of a Tissue-Like Memory (TLM) B cell phenotype through early EBV infection and apparent presence of EBV-induced TLM phenotype in LCLs.**

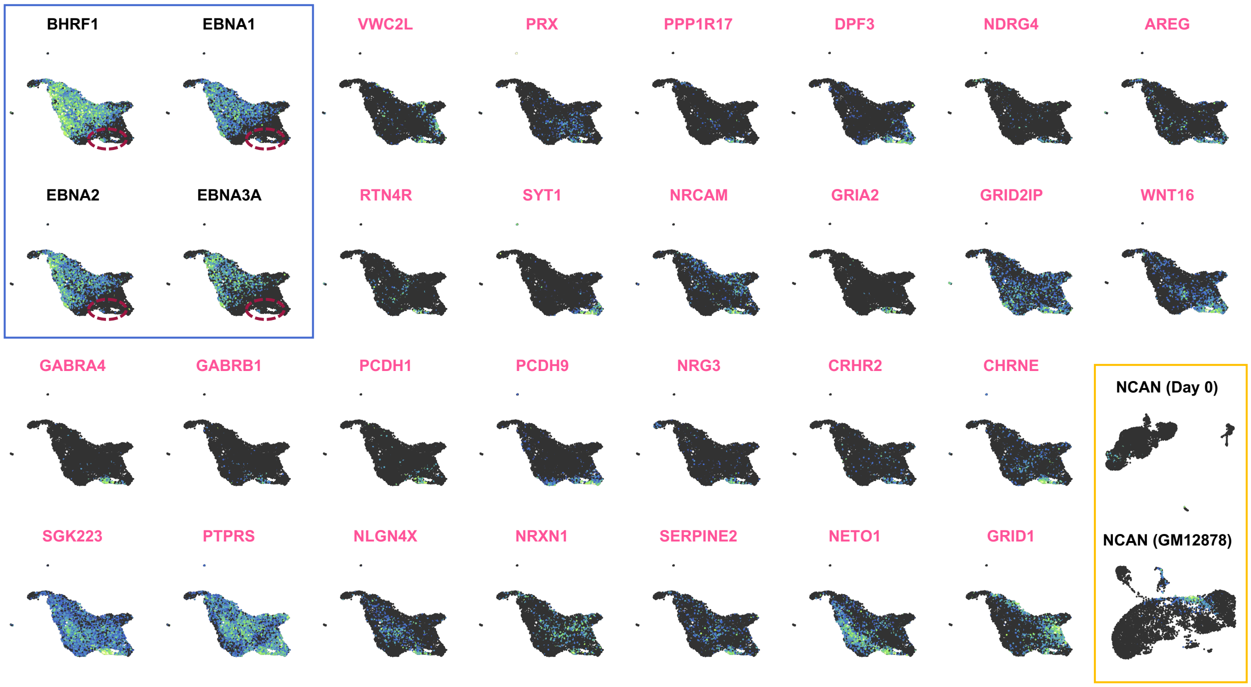


**Figure S44. Expression of EBV genes and neural cell adhesion molecules and signaling receptors in Tissue-Like Memory (TLM) B cells at Day 8 of early infection.** *BHRF1* and several EBV latency genes are shown in the blue box inset. The purple dashed oval highlights the location of FCRL4$^{+}$ TLM-like B cells at Day 8. Genes with known adhesion and signaling functions in the brain that exhibit upregulated expression of FCRL4+ TLM-like B cells at Day 8 are presented (pink labels). Expression of *NCAN* is shown at Day 0 in FCRL4+ TLM-like B cells and in the GM12878 LCL.

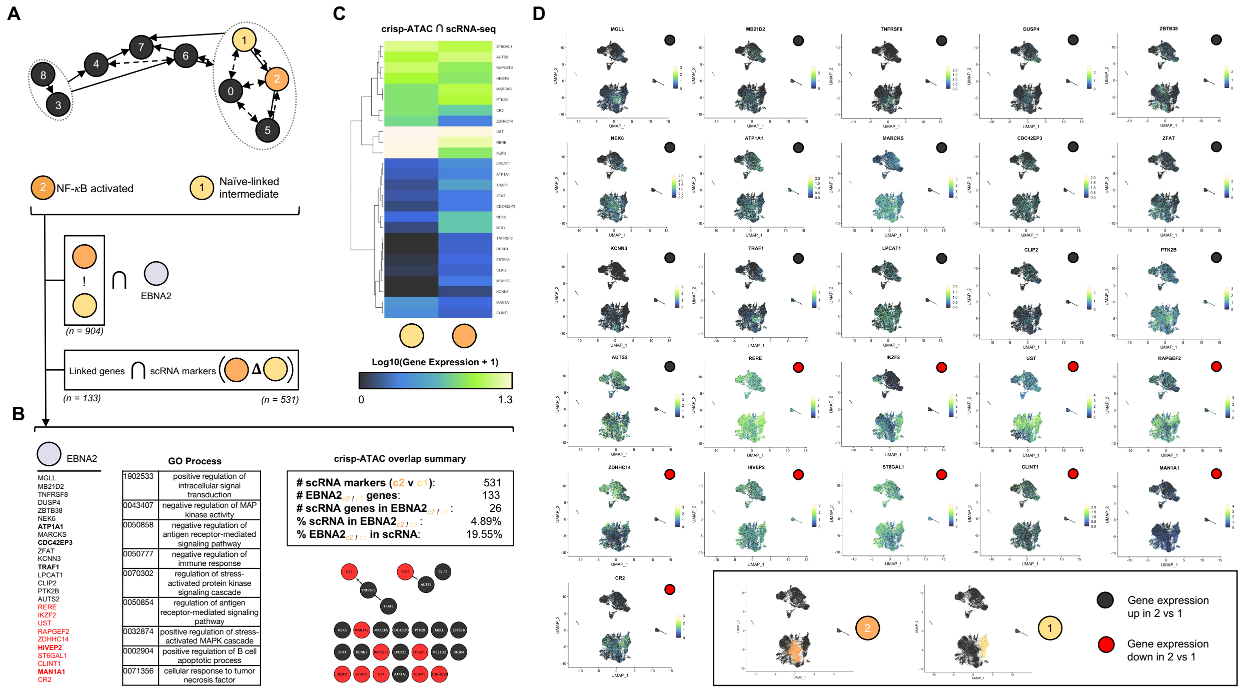


**Figure S45. crisp-ATAC analysis of DAP-linked DEGs associated with viral transcriptional co-activator binding sites in two activated states (TX1241).**

(A) Schematic overview of model states used in comparison (*c2 vs c1*). 904 DAPs were identified and intersected with EBNA2 ChIP data, resulting in 133 predicted linked genes. These gene predictions were cross-referenced with scRNA markers for the corresponding comparison (n = 531).

(B) 26 EBNA2-associated DAP-linked DEGs were identified, including five EBVSE-linked genes. These included 16 upregulated genes and 10 downregulated genes.

(C) Average expression of 26 EBNA2-associated DAP-linked DEGs in c1 cs c2.

(D) Gene expression UMAPs for each of the 26 identified genes.

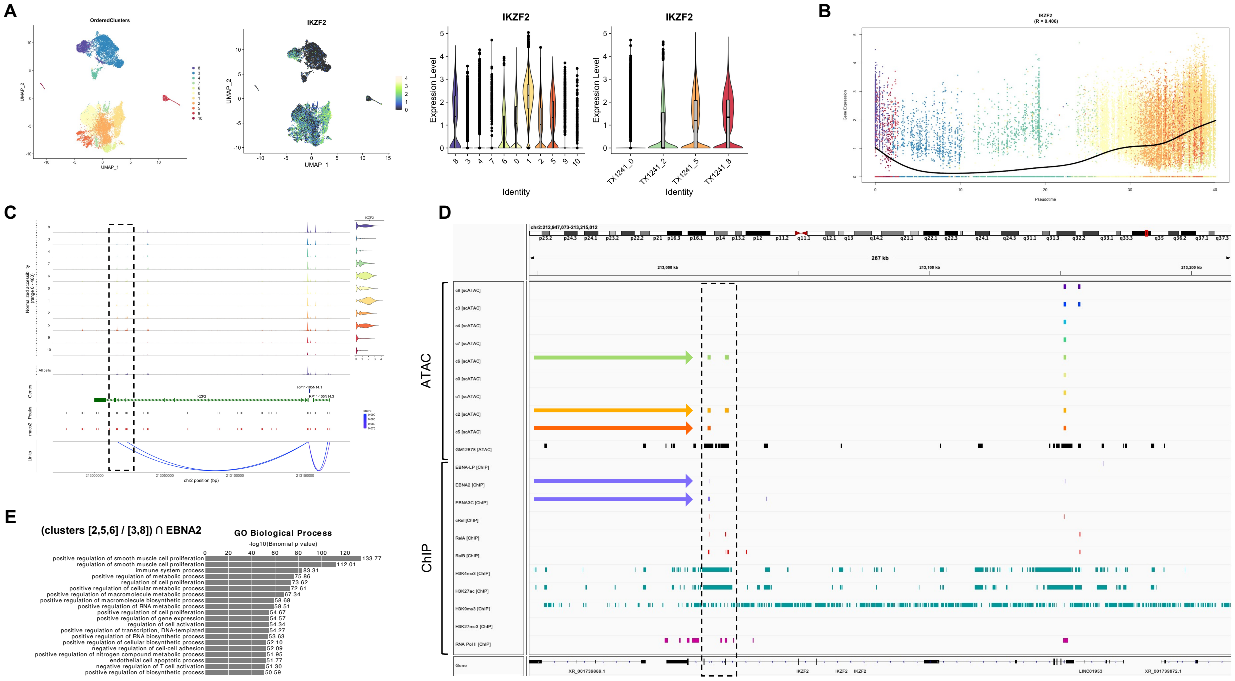


**Figure S46. Gene of interest analysis: *IKZF2*.**
(A) Expression characteristics of *IKZF2*.
(B) *IKZF2* expression in pseudotime.
(C) Joint coverage plot showing expression-linked peaks.
(D) EBNA and Rel-associated DAPs linked to *IKZF2* expression.
(E) Top GO term enrichment from the crisp-ATAC recipe used to identify *IKZF2* as a gene of interest.

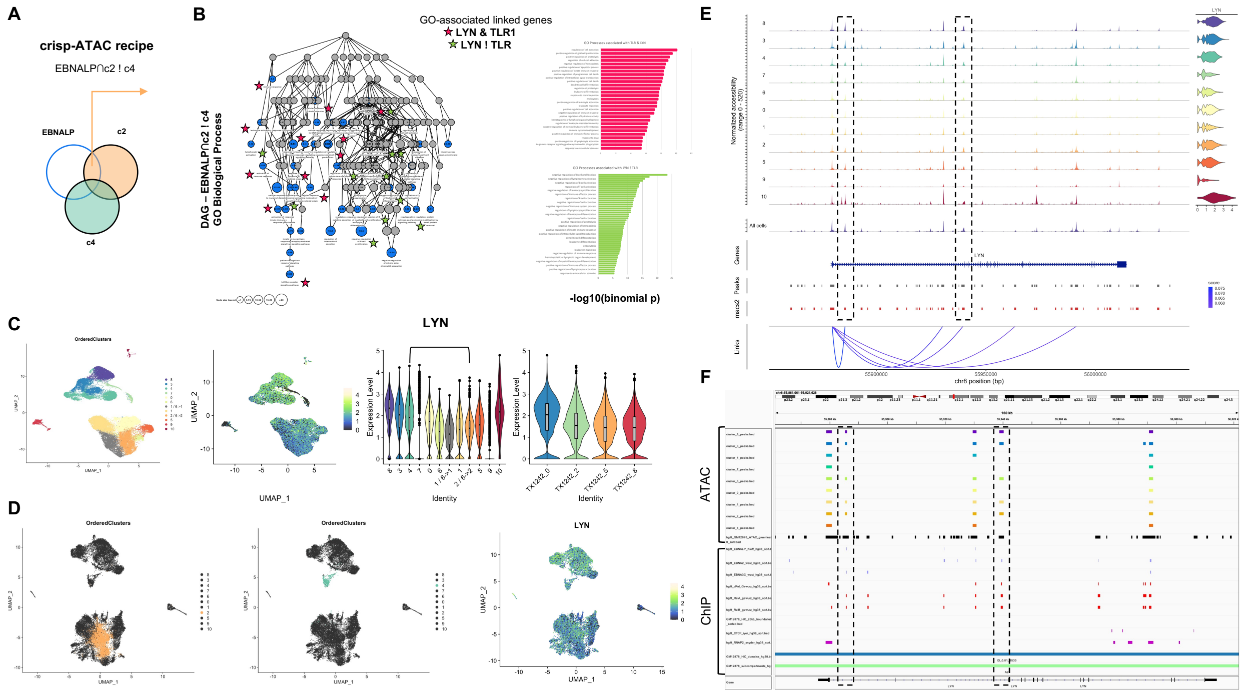


**Figure S47. Gene of interest analysis: *LYN*.**
(A) Schematic of used crisp-ATAC recipe.
(B) GO network (directed acyclic graph) with all *LYN*-associated terms denoted (with or without TLR gene associations as well).
(C) Expression profiling of *LYN*.
(D) Multimodal profiling of LYN expression and accessibility in *c2!c4* comparison.
(E) Expression-linked DAPs within the *LYN* locus.
(F) Overlap of expression-linked DAPs with EBNA and Rel family binding sites in the *LYN* locus.

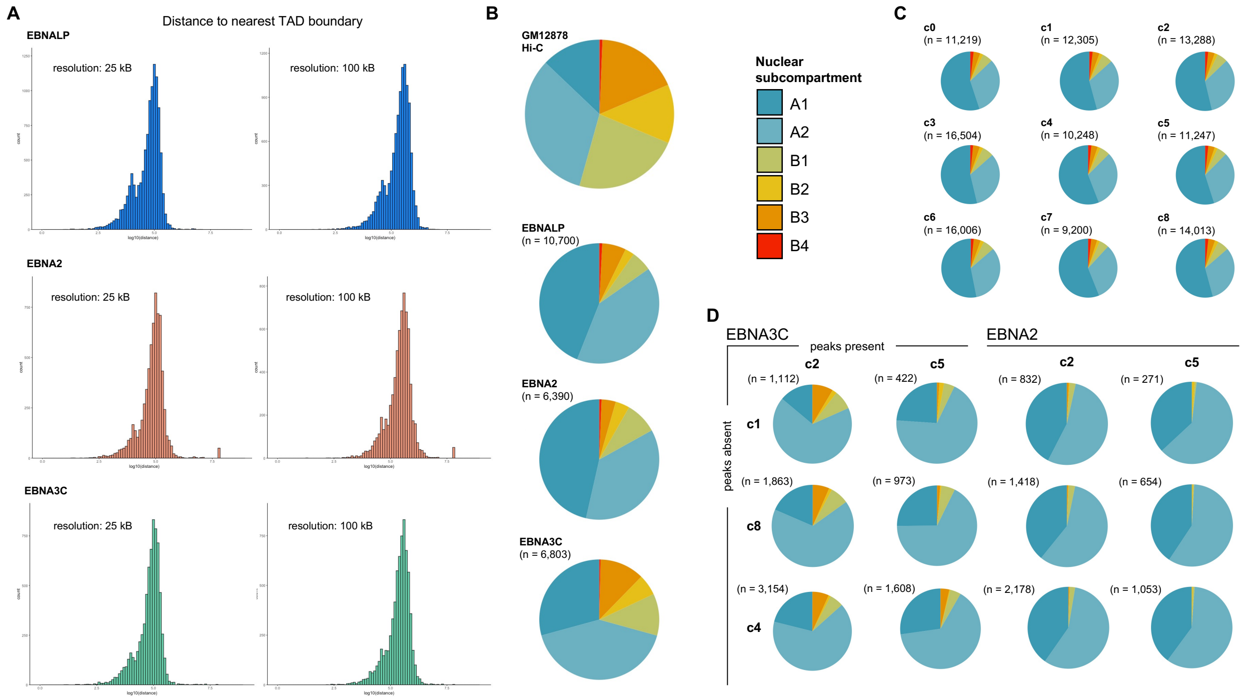


**Figure S48. Integration of Hi-C data using crisp-ATAC.**

(A) EBNA binding sites by distance to the nearest TAD identified at to Hi-C resolutions.

(B) Nuclear chromatin subcompartment proportions represented in GM12878 ensemble ATAC data (top chart) and in EBNA ChIP datasets (bottom 3 charts).

(C) Chromatin subcompartment breakdown by cell phenotype.

(D) Subcompartment frequencies of EBNA-associated DAPs between select cluster pairs.

| | TX1241 Day 0 | TX1241 Day 2 | TX1241 Day 5 | TX1241 Day 8 | TX1242 Day 0 | TX1242 Day 2 | TX1242 Day 5 | TX1242 Day 8 |
|---|---|---|---|---|---|---|---|---|
| **# cells (pre-QC)** | 8934 | 11087 | 20000 | 16161 | 13228 | 12252 | 12051 | 11914 |
| **# cells (post-QC)** | 8376 | 9212 | 19310 | 15373 | 12113 | 10601 | 11268 | 10938 |
| **# RNA features** | 36601 | 36601 | 36601 | 36601 | 36601 | 36601 | 36601 | 36601 |
| **# ATAC features** | 128651 | 178841 | 166703 | 176456 | 141025 | 168357 | 163878 | 170488 |
| **Mean unique RNAs / cell** | 2417 | 3198 | 2738 | 2893 | 2315 | 3150 | 3245 | 3337 |
| **Median unique RNAs / cell** | 2305 | 3281 | 2621 | 2761 | 2141 | 3199 | 3232 | 3274 |
| **Mean total RNAs / cell** | 6093 | 10317 | 7277 | 8151 | 5502 | 9692 | 9629 | 9976 |
| **Median total RNAs / cell** | 5420 | 9736 | 6377 | 7136 | 4714 | 8859 | 9015 | 9188 |
| **Mean unique peaks / cell** | 11185 | 10619 | 6439 | 8831 | 10646 | 9602 | 9296 | 12844 |
| **Median unique peaks / cell** | 12168 | 10403 | 6025 | 8767 | 11043 | 8834 | 9345 | 13441 |
| **Mean total peaks / cell** | 32091 | 27281 | 14769 | 21650 | 30286 | 23700 | 22802 | 34715 |
| **Median total peaks / cell** | 34675 | 26330 | 13297 | 20577 | 30906 | 21153 | 22019 | 35057 |
| **Mean mitochondrial gene %** | 9.49 | 10.45 | 9.78 | 11.03 | 6.81 | 9.89 | 9.29 | 8.53 |
| **Median mitochondrial gene %** | 9.23 | 10.15 | 9.58 | 10.75 | 6.59 | 9.52 | 9.03 | 8.14 |

**Table S1. Summary statistics and QC for single-cell multiomics data (all donors and timepoints).**

| Data type | Resolution | Target(s) | Cell type | Original publication | GEO Accession |
|---|---|---|---|---|---|
| ATAC-seq | single-cell (clusters) | N/A | early EBV$^+$ B lymphocytes | this study | --- |
| ATAC-seq | population ensemble | N/A | early EBV$^+$ B lymphocytes | this study | --- |
| ATAC-seq | population ensemble | N/A | LCL (GM12878) | Buenrostro *et al*, **Nat. Methods** (2013) | GSE47753 |
| ChIP-seq | population ensemble | EBNA1 | LCL | Tempera *et al*, **J. Virology** (2016) | GSE73887 |
| ChIP-seq | population ensemble | EBNALP | LCL (IB4) | Portal *et al*, **Proc. Nat. Acad. Sci.** (2013) | GSE49338 |
| ChIP-seq | population ensemble | EBNA2, EBNA3C | BL (Mutu III) | McClellan *et al*, **PLoS Pathogens** (2013) | GSE47629 |
| ChIP-seq | population ensemble | IgG (control) | LCL (GM12878) | Gunnell *et al*, **Nucleic Acids Res.** (2016) | GSE76869 |
| ChIP-seq | population ensemble | cRel, RelA, RelB | LCL (GM12878) | Zhao *et al*, **Cell Reports** (2014)<br>Iannetti *et al*, **PLoS Genetics** (2014) | GSE55105 |
| ChIP-seq | population ensemble | CTCF | LCL (GM12878) | Lee *et al*, **Genome Res** (2012) | GSE32883 |
| ChIP-seq | population ensemble | RNA Pol II | LCL (GM12878) | Raha *et al*, **Proc. Nat. Acad. Sci.** (2010) | GSE19550 |
| ChIP-seq | population ensemble | H2AZ, H3K4me1, H3K4me3, H3K9ac, H3K9me3, H3K27ac, H3K27me3, H3K36me3 | LCL (GM12878) | ENCODE Project Consortium, **Nature** (2012) | GSE29611 |
| Hi-C | population ensemble | N/A | LCL (GM12878) | Rao *et al*, **Cell** (2014) | GSE63525 |

**Table S2. ATAC-seq, ChIP-seq, and Hi-C datasets used in this work for crisp-ATAC.**
