## Supplementary Text for "Fate-resolved gene regulatory signatures of individual B lymphocytes in the early stages of Epstein-Barr Virus infection"

### Supplementary Information

### Supplementary Notes – Introduction

- LMP2A also promotes evasion by downregulating host factors that positively regulate surface expression of antigen-presenting MHC class II molecules (Lin et al., 2015; Rancan et al., 2015) and inhibiting interferon (IFN) responses along with LMP2B (Shah et al., 2009).

### Supplementary Notes – Results

#### Effects of cell cycle marker regression on cluster identification

Without cell cycle marker regression, all states except a discrete c2 were found via unsupervised methods. However, we identified two clusters in non-regressed datasets (c1/6→1, c2/6→2; shown for TX1242) as transitional phenotypes based on their hybrid transcriptomes (**Figure S1**). Each of these transitions is accounted for in our cell state model.

#### Overview of one-vs-all other cluster gene markers

As expected, many genes known to be modulated in EBV infection were identified as high-variance features from one-vs-all-other state comparisons. These included IFN targets and interleukin family genes (*EBI3, IFI6, IFI16, IFI44, IFI44L, IRF4, ISG15, SSBP2*) (Dooley et al., 2018; Klein et al., 2006; Liang et al., 2005; Pflanz et al., 2002; Schoggins et al., 2011; Wang et al., 2017a; Xu et al., 2008); genes involved in B cell activation (*NFKBIA, SLAMF1, SLAMF7, STAT1, TRAF1*) (Greenfeld et al., 2015; Hoffmann et al., 2002; Kim et al., 2016; Tsitsikov et al., 2001; Wood et al., 2007; Wuerzberger-Davis et al., 2011; Yoon et al., 2020) and differentiation

(*CD38, MZB1, PRDM1/BLIMP1, XBP1*) (Mrozek-Gorska et al., 2019; Nutt et al., 2015; Rosenbaum et al., 2014; Terstappen et al., 1990; Vrzalikova et al., 2011); apoptotic regulators (*BCL2A1, TNFAIP3, TNFRSF8*) (Fries et al., 1996; Izumi, 1997; Messinger et al., 2019; Price et al., 2017); chemokine and immune response-linked receptors (*CCR7, CR2, CXCR4, FCER2/CD23, FCRL1, FCRL5*) (Bonnefoy et al., 1995; Dement-Brown et al., 2012; Ghobrial et al., 2004; Nakayama et al., 2002; Nemerow et al., 1985; Price et al., 2012); non-coding RNAs (*FIRRE, PVT1, MIR155HG, MIR181A1HG, LINC00158*) (Hacisuleyman et al., 2014; Linnstaedt et al., 2010; Lu et al., 2008; Shi et al., 2019; Wood et al., 2018; Yang et al., 2020); and a broad array of TFs (*ARID5B*, *BACH2, EBF1, IKZF2, HIVEP3, JAZF1, NFKB1, REL, RUNX1, RUNX3, ZBTB38*) (Allen et al., 2002; Goodings-Harris et al., 2018; Huang et al., 2014; Hunter et al., 2016; Kobiita et al., 2020; Lu et al., 2016; Miotto et al., 2018; Muto et al., 1998; Park et al., 2019; Pozner et al., 2018; Schwartz et al., 2016; Spender et al., 2002; Swaminathan et al., 2013; Zhou et al., 2015) (**Figure S3**). The lymphoid oncogenic mediator *AFF3/LAF4* (Lefevre et al., 2015; Ma and Staudt, 1996; Wang et al., 2017b) was significantly downregulated upon EBV infection, as was the growth suppressor *SESN3*. The hydrolase *MACROD2*, a marker of the ATM-mediated DNA damage response, was significantly upregulated from c3 and c8 to c4 (Golia et al., 2017). Relative to other post-infection states, c2 exhibited elevated expression of the adhesion molecule *NTNG1* (Yaguchi et al., 2014) and *HIVEP3* (Hicar et al., 2001). Cells in c7 also exhibited elevated expression of genes associated with pro- and anti-apoptotic regulation (*RACK1*, *MIF*) (Mamidipudi and Cartwright, 2009; Nguyen et al., 2003).

All expression differences identified through cluster comparisons cited herein and subsequently were statistically significant after accounting for multiple hypothesis testing (Bonferroni corrected, p-adjusted < $1 \times 10^{-30}$). Average fold changes (log2FC), cluster frequencies, and statistics for genes referenced in this work are provided in a supplementary file.

<u>Details of pair- and group-wise cluster comparisons for phenotype annotation</u>

Relative to resting B cells (c3, c8), top markers of cells in c4 included elevated levels of genes involved in type I IFN response (*IFI44L, ISG15*) (DeDiego et al., 2019; Kanda et al., 1999; Schoggins et al., 2011) and inhibition of actin polymerization via sequestration (*TMSB4X, TMSB10*) (Safer et al., 1990). Conversely, c4 exhibited reduced mitochondrial transcripts (*MT-ND3, MT-ND2, MD-ATP8*) relative to c3 and c8. Comparison of Day 0 clusters (c3, c8) with those enriched at Day 5 and Day 8 (c0, c1, c2, c5, c6) revealed the largest significant changes observed over the sampled timecourse. These included post-infection elevation of *IFI44L*, *ISG15*, *MIR155HG*, immunoglobulin (Ig) genes (*IGHM*, *JCHAIN*), and a highly expressed EBV gene,

*BHRF1*. Genes downregulated post-infection included *CXCR4*, *BACH2*, and the EBNA3C target *COBLL1* (Gillman et al., 2018). Similar comparisons were made between c4 and c7 versus c0, c1, c2, c5, and c6 (**Figure S5**).

Because c0, c1, c2, c5, and c6 comprised most post-infection states, we generated pairwise expression comparisons among them (**Figure S5**). Notably, c2 was enriched for host markers of EBV-induced B cell activation (*EBI3*, *TNFAIP3*, *MIR155HG*, *CD226*) (Dudziak et al., 2003; Fries et al., 1996; Linnstaedt et al., 2010; Lu et al., 2008; Wood et al., 2018; Yoon et al., 2020), the tumor suppressor gene *ST7* (Zenklusen et al., 2001), exclusive expression of *TNFRSF8/CD30* (Kanavaros et al., 1992; Smith et al., 1993; Tazzari, 1999), and several long non-coding RNAs (lncRNAs; *LINC00158*, *MALAT1*) (Gutschner et al., 2013; Liu et al., 2021; Wood et al., 2018). Viral *LMP-1* and host NF-κB targets were also higher in c2 relative to other states (**Figure S6**). Among infected cells, c1 was similar to c2 but displayed greater (and in some cases, near-unique) expression of certain genes. These included *SKAP1* (Raab et al., 2010), *AFF3*, and *BACH2*, which appeared to carry over from elevated levels in resting cells within cluster 8. Notably, c1 exhibited higher transcript levels of *CDKN2A*/*p16INK4a* (*p16*), an EBNA3C-repressed target whose expression blocks transformation (Skalska et al., 2013). Cells in c1 also had the highest expression of the TF *IKZF2* (Park et al., 2019), near-exclusive expression of the acute lymphoblastic leukemia-associated gene *SH3RF3*/*POSH2* (Wang et al., 2015; Zhang et al., 2020), and *FIRRE*, a cMYC-induced lncRNA that plays key roles in epigenetic mediation of nuclear architecture (Hacisuleyman et al., 2014; Shi et al., 2019; Yang et al., 2015) (**Figure S7**). Cells in c0 exhibited elevated levels of *MARCH1* (Wu et al., 2020), *BANK1* (Georg et al., 2020; Kozyrev et al., 2008) linked to carryover from resting cells, *FCRL5*, and *KLHL29* (Dhanoa et al., 2013) (**Figure S8**). We found c6 contained hallmarks of the cell cycle, proliferation-induced stress, and survival mediation (*MKI67*, *HMGB1, HSPA8, DNMT1*) (Chen et al., 2007; José-Enériz et al., 2008; Pallier et al., 2003; Stricher et al., 2013; Štros et al., 2009; Yamada and Maruyama, 2007), consistent with the cluster's high cell cycle score and transcript content (**Figure S9**). Upregulated markers in c5 included numerous plasmablast-like features including polyclonal Ig heavy and light chain genes (*IGHM*, *IGHA1*, *IGHG3*, *IGLC2*, *IGLC3, JCHAIN*) (Calame, 2001; Gerster et al., 1986; Young et al., 1994) and high levels of *MZB1*, *CD38*, *XBP1*, and *PRDM1*/*BLIMP1* (**Figure S10**).

Given the cluster-level specificity of transcript similarity for certain genes between resting and infected cells (carryover), we examined heterogeneity in resting cells (c3, c8). Cells in c3 were consistent with resting memory B cells (*IGHD*$^{-}$, IGHM$^{+}$, *CD27*$^{+}$) while c8 had a naïve B cell profile (*IGHD*$^{+}$, *IGHM*$^{+}$, *CD27*$^{-}$) (Agematsu et al., 2000; Calame, 2001). Resting B cell features including

*SKAP1*, *ATXN1*, and *SORL1* (Uhlen et al., 2019) were differentially expressed between naïve and memory cells, which facilitated lineage tracing among post-infection phenotypes, most evident as linkage between c8 and c1 (**Figure S11**).

GO processes for one-vs-all-other cluster gene networks (**Figures S12-S16**)

Top GO process terms enriched in cluster 8 were related to regulation of immune receptor-mediated signaling (GO.0023057, FDR=2.1e-4; GO.0050854, FDR=4.2e-4; GO.0002768, FDR=9.8e-4) and negative regulation of macromolecule biosynthesis (GO.2000113, FDR=8.16e-6). GO enrichment in cluster 3 included regulation of cytokine production (GO.0001817, FDR=2.4e-4) and differentiated leukocyte processes (GO.0002521, FDR=2.1e-3; GO.0002381, FDR=2.5e-3; GO.1902105, FDR=4.1e-3). Cluster 4 was enriched for gene expression downregulation (GO.0010629, FDR=1.03e-12; GO.0000122, FDR=2.8e-3), IFN and stress responses (GO.0051412, FDR=8.4e-4; GO.0009605, FDR=8.7e-4; GO.0035458, FDR=1.58e-2), and pro-apoptotic processes (GO.0043065, FDR=7.4e-3; GO.1903209, FDR=1.76e-2). Cluster 7 was also defined by pro-apoptotic pathways (GO.1901798, FDR=3.4e-3; GO.2001244, FDR=3.5e-3) as well as DNA damage-induced mitotic arrest (GO.1902229, FDR=9.3e-4; GO.0007093, FDR=1.58e-2) and near-exclusive enrichment of ribosome biogenesis terms (GO.0042254, FDR=8.65e-37; GO.0042273, FDR=5.99e-20; GO.0042274, FDR=1.02e-16). Cluster 0 exhibited clear hallmarks of antiviral response (GO.0009607, FDR=1.62e-6; GO.0048584, FDR=1.3e-4, GO, GO.0009615, FDR=8.2e-3; GO.0031347, FDR=8.2e-3) and immune activation signaling (GO.0007166, FDR=9.1e-5; GO.0002235, FDR=1.8e-3; GO.1902531, FDR=2.6e-3). Cluster 6 was distinguished by cell division (GO.0051301, FDR=4.61e-54; GO.0140014, FDR=2.7e-30; GO.0007346, FDR = 3.53e-24). Cluster 1 displayed a diverse cellular milieu enriched for surface receptor signaling (GO.0007166, FDR=9.1e-4; GO.0002429, FDR=2.26e-2), immune defense responses (GO.0002682, FDR=2.6e-3; GO.0098542, FDR=9.2e-3), cell proliferation (GO.00042127, FDR=7.7e-3; GO.2000045, FDR=2.62e-2), and cellular protein catabolism (GO.1903362, FDR=2.39e-2). Likewise, cluster 2 exhibited diverse (but more specific) term enrichment for B cell activation signaling (GO.0033209, FDR=3.5e-4; GO.2000349, FDR=1.1e-3; GO.0050864, FDR=1.1e-3), cell proliferation, DNA damage, negative arrest regulation (GO.0006260, FDR=3.8e-6; GO.0008283, FDR=5.73e-5; GO.0042770, FDR=7.6e-4; GO.1901992, FDR=1.4e-3; GO.0071157, FDR=1.6e-3), and response to another organism (GO.0044419, FDR=8.47e-5; GO.0051707, FDR=8.6e-4). Lastly, cluster 5 displayed biotic (GO.0009607, FDR=1.21e-5; GO.0060337, FDR=3e-4) and apoptotic responses (GO.0010941, FDR=2.71e-6; GO.0006195, FDR=5.5e-4) and was uniquely enriched among all clusters for protein folding quality control responses (GO.0034976, FDR=2.46e-14; GO.0006457, FDR=9.79e-8; GO.0036498, FDR=6.75e-6, GO.0036500, FDR=4e-4).

Details of gene regulatory patterns in select genes (**Figures S28-S30**)

Post-infection downregulation of the chemokine receptor genes *CCR7* and *CXCR4* were each significantly linked to a balance of accessible sites with positive and negative regulatory ATAC sites. For *CCR7*, these included negative regulatory sites approximately +1 kb, +2 kb, and -7 kb from the gene TSS and positive regulatory sites approximately +18 kb, -12 kb, and 0 kb from the TSS (**Figure 28C**). Notably, *CXCR4* was linked to positive regulatory sites at +18 kb and +130 kb that became inaccessible after infection and coincide with EBNA3C binding sites identified in public ChIP-seq datasets (**Figure S29**). Other genes exhibiting significant, phenotype-specific mélanges of regulatory links included the TFs *BACH2* (+/+ in $EBV^-$ naïve cells; -/- in $EBV^+$ differentiated cells) and *RUNX3* (-/+ in $EBV^+$ activated B cells; -/- in $EBV^-$ naïve cells) (**Figure 28C**). Curiously, the IgM crosslinker gene *JCHAIN* (+/- in $EBV^-$ naïve cells; -/+ in $EBV^+$ differentiated cells), was significantly linked (correlation score=-.12, p<0.05) to an upstream negative regulatory site (-6 kb) that coincided with an EBNALP / cREL / RelA / RelB binding site and the TSS of an adjacent gene (*UTP3*, which exhibited neither significant differential accessibility nor expression). In addition to sites linked to *JCHAIN* (which undergoes essential transcriptional upregulation during plasmablast formation) (Yagi and Koshland, 1981), we identified phenotype-specific, DAPs coincident with EBNALP / [EBNA3C] / [EBNA2] / Rel sites approximately 12 kb, 55 kb, and 67 kb downstream of the TSS for *PRDM1*/*BLIMP1*, a key transcriptional regulator of B cell differentiation (**Figure S30**).

Differentially enriched binding motifs by time and phenotype

24% of the top 100 enriched TF motifs were conserved across timepoints (multi-phenotype ensembles), whereas only 9% of the top 100 enriched motifs were common when parsed by phenotypes corresponding to EBV infection of naïve cells (**Figure S37A**, top two panels). As expected, the most enriched TF motifs in all phenotypes were associated with regulation of RNA Pol II transcription (GO.0000122, GO.0006357). 61 motifs were shared between resting memory and naïve B cells, accounting for 44% of all unique top 100 enriched motifs present within these two cell states (n=139). Top enriched motifs within naïve cells included sites for TFs associated with earlier developmental and hematopoietic processes (GO.1903708, FDR=6.4e-4; GO.0009888, FDR=2.73e-6) relative to top motifs in memory cells. By comparison, only 5% of the top 100 motifs by cluster were common across non-arrested infected states. Notable gene ontology terms for TFs with enriched motif accessibility in the NF-κB activated B cell state (c2) included responses to stress (GO.0043620, FDR=5.07e-7) and other organisms (GO.0051707, FDR=5.02e-7), including type I IFN induction (GO.0060337, FDR=2.2e-4). The EBV$^+$ plasmablast state was likewise enriched for IFN binding sites (GO.0060337, FDR=4.83e-8) as well as motifs for TFs involved in cell differentiation (GO.0045595, FDR=2.99e-7) and unfolded protein stress (GO.0006990, FDR=1.33e-2).

#### crisp-ATAC testing and epigenetic pattern quantification

A typical crisp-ATAC pipeline begins with curation of population ensemble ChIP-seq and ATAC-seq datasets for host and/or virus TF binding sites and epigenetic marks from a reference state (in our case the EBV$^+$ LCL, GM12878). Publicly available datasets used for crisp-ATAC in this study are provided in **Table S2**. Curated data are prepared as called peak ranges (.bed) with MACS2 (Liu, 2014), which are then intersected with state-specific ATAC peaks using *bedtools* (Quinlan and Hall, 2010) to identify common genomic ranges. Combinatorial logic gates (crisp-ATAC recipes) are applied to the resulting intersection matrix to identify phenotype-resolved TF-associated DAPs and their epigenetic signatures. Genomic intervals that pass all imposed gates are analyzed with GREAT (McLean et al., 2010) to predict DAP-linked *cis*-regulated genes. When matched scRNA and scATAC data are available, output DAP-linked gene predictions can be cross-referenced with empirical DEGs from the same phenotype comparison.

We initially evaluated correlations among ATAC phenotypes and LCL ensemble ATAC- and ChIP data (**Figure S39B**). Genomic range vectors for single-cell clusters were more correlated with each other ($0.75<R<0.93$) than with any ensemble dataset, likely due to global differences in data sparsity, modality, and biological targets. However, cluster ATAC profiles were mildly positively correlated with activating epigenetic marks ($0.18<R<0.28$), various host and viral TFs ($0.02<R<0.21$), and RNA Pol II ChIP sites ($0.318<R<0.331$).

Intersection of scATAC peaks with viral EBNA1 and epigenetic signature ChIP-seq peaks revealed negligible differences in the frequency of EBNA1-associated enhancers (enh), promoters (pro), or transcribed gene regions (gen) among phenotypes. Recipes were readily extensible to evaluate multiplexed TF binding sites. For example, c2, c3, and c6 had a group average of 1209+/-66 genome-wide accessible intervals associated with cRel and EBNA2 but not EBNA3C binding [(EBNA2$\cap$cREL)/EBNA3C], which was 50% greater than in clusters 4,7,9, and 10 (806+/-43) and 30% more than in clusters 0,1,5, and 8 (930+/-46) (**Figure S39C**, right column).

#### EBNA-specific GO from successful infection trajectory – *c256!c38* crisp-ATAC (**Figure S41**)

EBNA2-specific *c256!c38* site-linked genes were enriched for negative regulation of lymphocyte activation (GO.0051250, -log10(p)=41.15), cell differentiation (GO.0045596, -log10(p)=40.58), and leukocyte cell adhesion (GO.1903038, -log10(p)=47.93); and positive regulation of cell motility (GO.2000147, -log10(p) = 35.70), phosphorylation (GO.0042307, -log10(p)=38.76), and leukocyte-mediated immunity (GO.0002444, -log10(p)=33.74). EBNA3C-specific *c256!c38* site-linked genes corresponded to negative regulation of B cell proliferation (GO.0030889, -log10(p)=30.21), activation, (GO.0050869, -log10(p)=25.44), regulation of germinal center formation (GO.0002634, -log10(p)=30.14), and negative regulation of TF binding (GO.0043433, -log10(p)=35.85). EBNALP-specific intersected *c256!c38* site-linked gene sets were enriched for positive regulation of lymphocyte activation (GO.0051251, -log10(p)=9.64), proliferation (GO.0050671, -log10(p)=9.53), and regulation of *TNF* family cytokine production (GO.1903555, -log10(p)=16.44).

Additional crisp-ATAC recipes and results (**Figures S45-S48**)

*c2!c5*: Other EBNA site-linked genes with large *c2!c5* expression fold changes included *MARCKS, CYTIP, STK17B, OXR1, HIVEP3, ANK2, FBXW7, MGLL, ST7, LTA, and ZBTB38* (**Figure S45**).

*c2!c1*: We used crisp-ATAC to finely dissect differences in predicted EBNA2 regulatory relationships between two closely related cell states – the canonical EBV-induced NF-κB activated state (c2) and the naïve-linked activation intermediate (c1). A total of 904 EBNA2-associated *c2!c1* DAPs were identified and linked to 133 unique genes (**Figure S45A**). 19.6% of $EBNA2_{c2!c1}$ predicted genes (26 of 133) were also *c2!c1* DEGs measured using scRNA-seq, accounting for 4.89% of all significant *scRNA(c2!c1)* DEGs. The 26 EBNA2-associated DAP-linked DEGs (16 upregulated, 10 downregulated) were involved in GO processes known to be essential for successful establishment of latent EBV infection (**Figure S45B-D).** Further, four upregulated genes (*ATP1A, MARCKS, CDC42EP3, TRAF1*) and one downregulated gene (*HIVEP2*) were associated with EBVSEs. We confirmed the presence of EBNA2 DAPs for individual scRNA-matched gene predictions.

Analyses of other genes of interest (*IKZF2, LYN*) identified using distinct crisp-ATAC recipes were also demonstrated (**Figures 46-47**).

Extension of crisp-ATAC concept for integration of Hi-C reference data

We extended the crisp-ATAC concept to integrate all-vs-all chromatin conformation capture data (Hi-C) to study TF- and phenotype-resolved accessibility differences by their frequencies within distinct nuclear subcompartments and distances to local topologically associated domain (TAD) boundaries (**Figure S48**). Generally, differentially-accessible TF-associated sites are enriched within euchromatin (compartments A1, A2) relative to genome-wide ChIP and ensemble ATAC compartmentalization in the GM12878 cell line. There is some variation by phenotype and viral transcriptional activators, most notably as greater proportions of c2 (NF-κB phenotype) and EBNA3C peaks within heterochromatic subcompartments. In each case, the proportion of peaks within repressed compartments is still notably less than the genome-wide frequency of these compartments.

### Supplementary Notes – Discussion

c47!c38 DAP-linked DEGs: ribosomal subunit genes

We observed that RPS27A exhibited elevated expression in arrested relative to resting states that was linked to increased accessibility at multiple negative regulatory sites. Notably, *RPS27A* is a p53 target that is transcriptionally upregulated in response to DNA damage (Deisenroth and Zhang, 2010; Nosrati et al., 2014; Nosrati et al., 2015). Further, it encodes a ubiquitin fusion protein (Kirschner and Stratakis, 2000) that stabilizes viral LMP1 to promote cell proliferation (Hong et al., 2017).
